# High-resolution animal tracking with integration of environmental information in aquatic systems

**DOI:** 10.1101/2020.02.25.963926

**Authors:** Fritz A Francisco, Paul Nührenberg, Alex Jordan

## Abstract

Acquiring high resolution quantitative behavioural data underwater often involves installation of costly infrastructure, or capture and manipulation animals. Aquatic movement ecology can therefore be limited in scope of taxonomic and ecological coverage. Here we present a novel deep-learning based, multi-individual tracking approach, which incorporates Structure-from-Motion in order to determine the 3D location, body position and the visual environment of every recorded individual. The application is based on low-cost cameras and does not require the animals to be confined or handled in any way. Using this approach, single individuals, small heterospecific groups and schools of fish were tracked in freshwater and marine environments of varying complexity. Further, we established accuracy measures, resulting in positional tracking errors as low as 1.09 ± 0.47 cm (RSME) in underwater areas up to 500 m^2^. This cost-effective and open-source framework allows the analysis of animal behaviour in aquatic systems at an unprecedented resolution. Implementing this versatile approach, quantitative behavioural analysis can employed in a wide range of natural contexts, vastly expanding our potential for examining non-model systems and species.

## Background

Understanding the movement and behaviour of animals in their natural habitats is the ultimate goal of behavioural and movement ecology. By situating our studies in the natural world, we have the potential to uncover the processes of selection acting on the behaviour in natural populations in a manner that cannot be achieved through lab studies alone. The ongoing advance of animal tracking and biologging brings the opportunity to revolutionize not only the scale of data collected from wild systems, but also the types of questions that can subsequently be answered. Incorporating geographical data has already given insights, for example, into the homing behaviour of reef fish, migratory patterns of birds, or the breeding site specificity of sea turtles [10, 22, 51]. Great advances in systems biology have further been made through the study of movement ecology, understanding the decision-making processes at play within primate groups manoeuvring through difficult terrain or the collective sensing of birds traversing their physical environment [41, 52]. Unravelling these aspects of animal movement can also vastly improve management strategies [13, 14], for example in the creation of protected areas that incorporate bird migratory routes [48] or by reducing by-catch with dynamic habitat usage models of marine turtles [37].

Yet the application of techniques that meet the challenges of working in naturally complex environments is not straightforward, with practical, financial, and analytical issues often limiting the resolution or coverage of data gathered. This is especially true in aquatic ecosystems, where approaches such as Global Positioning System (GPS) tags allow only sparse positioning of animals that surface more or less frequently, or Pop-up Satellite Archival Tags (PSATs) which integrate surface positions with logged gyroscope and accelerometer data to estimate movement of larger aquatic animals [29, 54]. Not only does the spatial resolution of respective tracking systems, e.g. currently 4.9m for GPS, limit the possibilities of behavioural analyses on a fine scale, but also excludes almost all animals below a certain size class [56]. This is problematic because in aquatic ecosystems, as in terrestrial systems, life is numerically dominated by small animals [3]. Ultrasonic acoustic telemetry is one methodology useful for underwater tracking of smaller animals and those in larger groups [29, 35], but this approach is limited to a stationary site through the positioning of the acoustic receivers, and the costs, maintenance, and installation of these systems preclude their effective use in the majority of systems and for many users. These methods also require animals to be captured and equipped with tags that should not exceed 5% of the animals weight [17, 33, 35], rendering current generation GPS and PSATs problematic for small animals. While acoustic tags are small enough for injection, the increased handling time associated with these invasive measures can lead to additional stress for the animals, whereas the tag itself may disturb the animals’ natural behaviour [30]. Hence, approaches that facilitate collection of behavioural data in smaller animals, those in large groups, and those in varied aquatic habitats, are still lacking.

A lack of data becomes a fundamental problem if certain ecosystems, species, or habitat types are underrepresented in terms of adequate research, management, or discovery. Although the oceans constitute up to 90% of habitable ecosystems worldwide, as little as 5% have been explored [25, 40, 42]. Within the oceans, coastal inshore areas have the greatest species diversity, with approximately 80% of fish species (the most speciose group of vertebrates) inhabiting the shallow waters of the littoral zone [46], and providing over 75% of commercial seafood landings [20]. Coastal regions in both marine and freshwater environments are also those that are of greatest interest for eco-tourism, community fisheries, and industry, while simultaneously being most affected by habitat degradation, exploitation, and anthropogenic pollution [11, 12, 15]. Knowledge of the coastal regions is essential for establishing sanctuaries and sustainable concepts of ocean preservation [21] and movement data plays a vital role in this process, in that it gives detailed information about the location, preferred habitat and temporal distribution of organisms [33]. Yet for reasons of animal size, species abundance, and habitat complexity, most available tracking methods are poorly suited to these inshore regions.

Application of appropriate tracking and behavioural analysis techniques in a flexible, accessible, and broadly applicable manner would alleviate these limitations in systems and species coverage, improving capacity for conservation, management, and scientific understanding of natural systems across scales and conditions. In pure research terms, the application of quantitative behavioural and movement analyses in natural settings would also help bridge the gap between quantitative lab-based research and often qualitative field-based research. Recent advances in behavioural decomposition [6, 27] may then be employed in field settings, vastly improving our understanding of behaviour and movement in the wild.

Here we present an open-source, low-cost approach based on consumer grade cameras to quantify the movement and behaviour of animals of various sizes in coastal marine and freshwater ecosystems. Our approach integrates two methodologies from the field of computer vision, object detection with deep neural networks and Structure-from-Motion (SfM). Object detection has been successfully employed in terrestrial systems for animal localization, yielding highly resolved movement data through e.g. drone-based videos over broad environmental contexts [28]. While these aerial approaches may also be used in some aquatic systems, they are limited to extremely shallow water and large animals [45]. The approach we advocate allows data to be collected on any animal that can be visualized with cameras, enabling it also for smaller fish and other aquatic animals. In addition to solely providing animal trajectories, video-based observations also entail environmental data that adds the possibility to study interactions of mobile animals with their natural habitat [52]. Our approach synthesizes object detection with SfM into a coherent framework that can be deployed in a variety of systems without domain-specific expertise. SfM is commonly used for 3D environmental reconstructions, photogrammetry and camera tracking for visual effects in video editing [5, 59], and here allows the reconstruction of 3D models of the terrain through which the animals move and interact with. Our open-source analysis pathway enables subsequent calculation of movement, interactions, and postures of animals. Set-up costs can be as small as two commonly available action cameras, and the proposed method can be taken into habitats which are otherwise explored by snorkeling, diving, or with the use of remotely operated underwater vehicles (ROVs). Analysis can be performed on local GPU-accelerated machines or widely-accessible computing services (e.g. Google Colaboratory). Overall, this method provides a low-cost approach for measuring the movement and behaviour of aquatic animals that can be implemented across scales and contexts.

## Methods

Three datasets of varying complexity were used to demonstrate the versatility of the proposed method. These were chosen to range from single animals (*Conger conger*) and small heterospecific groups (*Mullus surmuletus, Diplodus vulgaris*) to schools of conspecific individuals (*Lamprologus callipterus*) under simple and complex environmental conditions, resulting in the datasets *‘single’, ‘mixed’* and *‘school’* respectively. Moreover, we provide a dataset of four repeated trials (*‘accuracy’*) to validate the accuracy of our tracking approach. This dataset was used to reconstruct the trajectories of a calibration wand and examine resulting tracking errors. The *‘single’* and *‘mixed’* datasets were created while snorkeling at the surface, using a stereo camera set-up at STARESO, Corsica (Submarine and Oceanographic Research Station). The remaining datasets were collected via SCUBA (5-8 m) with either multi or stereo camera set-ups in Lake Tanganyika, Zambia (Tanganyika Science Lodge, Mpulungu) and STARESO. While the *‘single’* and *‘mixed’* datasets were recorded with untagged fish, we attached tags made of waterproof paper (8×8 mm) anterior to the dorsal fin of the individuals for the *‘school’* dataset [23]. See Tab. 1 for a summary of the collected datasets. For a general guideline and comments on the practical implementation of our method, refer to Additional file 7.

**Table 1.** Summary of acquired datasets. *Camera distances* lists the minimum camera-to-camera distance in the setups, *tags* whether individual animals were tagged, *pose* if animal spine pose estimation was used during tracking. *Complexity* lists an estimate of overall complexity (number of individuals, environment). NA: not applicable.

| Dataset | Location | Approach | Species | Setup | Camera distances (m) | Tags | Pose | Complexity |
| --- | --- | --- | --- | --- | --- | --- | --- | --- |
| *single* | STARESO | schnorchel | *Conger conger* | stereo | 0.4 | no | yes | high |
| *mixed* | STARESO | schnorchel | *Mullus surmuletus*,<br>*Diplodus vulgaris* | stereo | 0.6 | no | yes | low |
| *school* | Tanganyika | dive | *Lamprologus callipterus* | multiple (12) | 0.2 | yes | no | medium |
| *accuracy* | STARESO | dive | NA | multiple (4) | 0.6 | NA | NA | varying |

### Automated animal detection and tracking

Since all data was collected in the form of videos, image-based animal detection was required for subsequent trajectory reconstruction and analyses. Firstly, the videos resulting from the stereo or multi-camera set-ups were synchronized using a convolution of Fourier-transformed audio signals to determine the video offsets (code available). Secondly, the synchronized videos were tracked independently using an implementation of a Mask and Region based Convolution Neural Network (Mask R-CNN) for precise object detection at a temporal resolution of either 30 Hz (*‘single’, ‘mixed’* and *‘accuracy’*) or 60 Hz (*‘school’*) [1, 26]. To this end, we trained Mask R-CNN models on small subsets of labeled video frames using transfer learning from a model that was pre-trained on the COCO dataset with more than 200K labeled images and 80 object classes [1, 38]. Our training sets contained 171, 80 and 160 labeled images for the *‘single’, ‘mixed’* and *‘school’* datasets respectively. For the *‘accuracy’* dataset, we annotated a total of 73 images. The original image resolutions of 2704 × 1520 px (*‘single’* and *‘school’*) and 3840 × 2160 px (*‘mixed’* and *‘accuracy’*) were downsampled to a maximum width of 1024 px while training and predicting to achieve better performance. After training, the models were able to accurately detect and segment the observed animals, which was visually confirmed with predictions on validation datasets.

The predicted masks were either used to estimate entire poses of the tracked animals (*‘single’, ‘mixed’*) or to calculate centroids of the tags or calibration wand ends in case of the *‘school’* and *‘accuracy’* datasets. Established morphological image processing was used to skeletonize the Mask R-CNN predictions, producing a 1 px midline for each of the detected binary masks. A fixed number of points was equidistantly distributed on these midlines as an estimation of the animals’ spine poses. Both the spine points and the tag centroids represent pixel coordinates of detected animals in further data processing. Partitioned trajectories were generated from detections with a simple combination of nearest-neighbors between subsequent frames or utilizing a cost-reduction algorithm (the Hungarian method), and filtering for linear motion over a short time window, reducing later quality control and manual track identification for continuous trajectories to a minimum. For video and image annotations, trajectory and pose visualization, manual track corrections and other trajectory utility functions, we developed a GUI based on Python and Qt5 within the lab (*‘TrackUtil’*, Additional file 4). The code for Mask R-CNN training and inference, video synchronization, fish pose estimation and automatic trajectory assignment is also available (Additional files 5 and 6). The training and tracking details are summarized in Tab. 2.

**Table 2.** Dataset parameters and accuracy metrics. *Annotations* lists how many frames were annotated for training Mask R-CNN, *rate* the frames per second of each video set, i.e. the temporal tracking resolution. *Resolution* is video resolution, 2.7k: 2704*×*1520 px, 4k: 3840*×*2160 px. *Reconstruction* metrics refer to the deviation of reconstructed camera-to-camera distances from the actual distance, *reprojection* metrics to the reprojection of triangulated 3D tracks to the original video pixel coordinates and *tracking* to the deviation of the tracked calibration wand length from its actual length. In case of the *‘accuracy’* dataset, the accuracy results are listed as the mean and standard deviation of the four repeated trials. NA: not applicable.

| Dataset | Annotations | Rate (Hz) | Resolution (px) | Accuracy metrics |  |  |  |
| --- | --- | --- | --- | --- | --- | --- | --- |
|  |  |  |  | Metric | Reconstruction (cm) | Reprojection (px) | Tracking (cm) |
| *single* | 171 | 30 | 2.7k | median | 0.30 | 11.51 | NA |
|  |  |  |  | RMSE | 1.28 | 20.97 | NA |
| *mixed* | 80 | 30 | 4k | median | 0.44 | 3.77 | NA |
|  |  |  |  | RMSE | 1.09 | 7.77 | NA |
| *school* | 160 | 60 | 2.7k | median | 0.06 | 2.57 | NA |
|  |  |  |  | RMSE | 0.30 | 3.78 | NA |
| *accuracy* | 73 | 30 | 4k | median | -0.14 ± 0.06 | 3.53 ± 1.96 | 0.14 ± 0.33 |
|  |  |  |  | RMSE | 1.34 ± 0.79 | 8.56 ± 5.21 | 1.09 ± 0.47 |

### Structure from motion

The field of computer vision has developed powerful techniques that have found applications in vastly different fields of science [19, 24, 58]. The concept of Structure-from-Motion (SfM) is one such method that addresses the large scale optimization problem of retrieving three dimensional information from planar images [39]. This approach relies on a static background scene, from which stationary features can be matched by observing them from different perspectives. This results in a set of images, in which feature-rich key points are first detected and subsequently used to compute a 3D reconstruction of the scene and the corresponding view point positions. As shown in (1) and (2), a real world 3D point *M*′ (consisting of *x, y, z*) can be projected to the image plane of an observing camera by multiplying the camera’s intrinsic matrix *K* (consisting of focal lengths *f*_*x*_, *f*_*y*_ and principal point *c*_*x*_, *c*_*y*_), with the camera’s joint rotation-translation matrix [*R|t*] and *M*′, resulting in the corresponding image point *m*′ (consisting of pixel coordinates *u, v*, scaled by *s*) [8]. By extension, this can be used to resolve the ray casting from a camera position towards the actual 3D coordinates of a point given the 2D image projection of that point with known camera parameters. Due to this projective geometry, it is not possible to infer at which depth a point is positioned on its ray from a single perspective. SfM is able to circumvent this problem by tracking mutually-observed image points (*m*′) across images of multiple camera view points. As a result, the points can be triangulated in 3D space (*M*′), representing the optimal intersections of their respective rays pointing from the cameras positions towards them. By minimizing reprojection errors, which are the pixel distances between the 3D points’ reprojections to the image planes and their original image coordinates (*u, v*), SfM is also able to numerically solve the multi-view system of the cameras relative rotation (*R*), translation (*t*) and intrinsic (*K*) matrices and to retrieve the optimal camera distortion parameters (*d*).

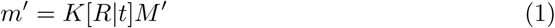

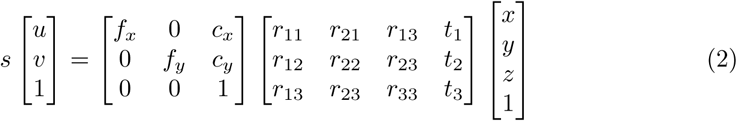

Here, SfM was incorporated into data processing in order to gain information about exact camera positions, which was done using the general-purpose and open-access pipeline COLMAP [49, 50]. The synchronized videos were resampled as image sequences with a rate of 3 Hz. In case of the *‘mixed’* dataset, we removed frames that were recorded when the cameras were stationary. The resulting image sequences served as input into the reconstruction process during which the cameras were calibrated (*K, d*) and relative extrinsic parameters (*R, t*) computed, so that all camera projections relate to a shared coordinate system. Every input image resulted in a corresponding position along the reconstructed, 3D camera path of the recording, where the number of images determined the temporal resolution of resolved camera motion. Since only a subset of all video frames were used for the reconstructions, SfM optimized a smaller number of parameters, resulting in a reduced computational load. Additionally, this could improve reconstruction accuracy, as the images still had sufficient visual overlap, but increased angles between viewpoints. Finally, the retrieved camera parameters were interpolated to match the acquisition rate of animal tracking, assuring that reference camera parameters are given for each recorded data point by simulating a continuous camera path.

### Reconstruction of animal trajectories

It is necessary to resolve the camera motion when tracking moving animals with non-stationary cameras, since the camera motion will also be represented in the pixel coordinate trajectories of the animals. With camera information (*K, d*) and relative perspective transformations (*R, t*) for the entire camera paths retrieved from SfM, and multi-view animal trajectories from the Mask R-CNN detection pipeline available, a triangulation approach similar to SfM can be used to compute 3D animal trajectories. Here, an animal’s pixel coordinates represent *m*′ (consisting of *u* and *v*) observed from more than one known view point (*R, t*), and the animals 3D positions *M*′ (*x, y, z*) can be triangulated. Positions of animals observed in exactly two cameras were triangulated using an OpenCV implementation of the direct linear transformation algorithm, while positions of animals observed in more than two cameras were triangulated using singular value decomposition following an OpenCV implementation [8, 24]. Additionally, positions of animals temporarily observed only in one camera were projected to the world coordinate frame by estimating the depth component as an interpolation of previous triangulation results. Through the recovered camera positions, the camera motion is nullified in the resulting 3D trajectories. Thus, they provide the same information as trajectories recorded with a fixed camera setup (Fig. 1). Animal trajectories and the corresponding reconstructions were scaled, so that the distances between the reconstructed camera locations equal the actual distances within the multi-view camera setup. As a result, all observations are represented on a real world scale. The code for trajectory triangulation, camera path interpolation and visualizations is bundled in a Python module (*‘multiviewtracks’*), accessible on GitHub [43].

**Figure 1.**
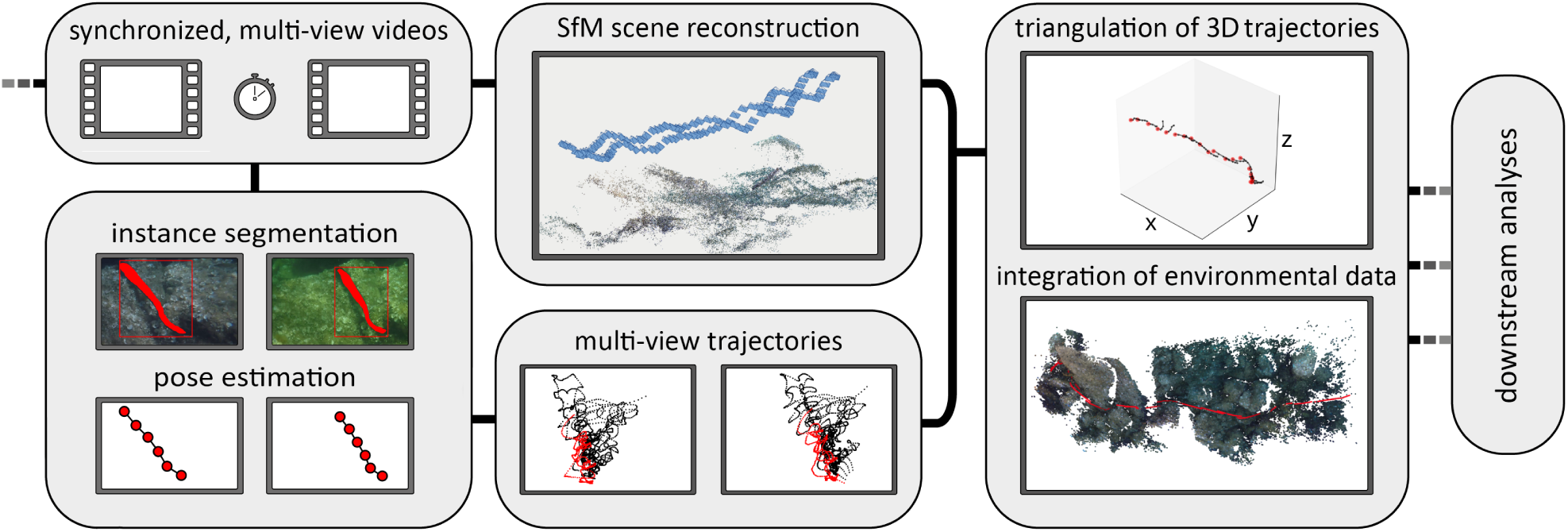
Schematic workflow. Data processing starts with the acquisition of synchronized, multi-view videos, which serve as input to the SfM reconstruction pipeline to recover camera positions and movement. In addition, Mask R-CNN predictions, after training the detection model on a subset of images, result in segmented masks for each video frame, from which animal poses can be estimated. These serve as locations of multi-view animals trajectories in the pixel coordinate system. Subsequently, trajectories can be triangulated using known camera parameters and positions from the SfM pipeline, yielding 3D animal trajectories and poses. Integrating the environmental information from the scene reconstruction, these data can be used for in depth downstream analyses.

## Results

Here we combined Mask-RCNN aided animal detection and tracking with SfM scene reconstruction and triangulation of 3D animal trajectories to obtain high resolution data directly from videos taken while snorkeling or diving in the field.

Given that the proposed method incorporates out-of-domain and novel approaches from computer vision, reliable accuracy measures are required. Therefore, a ground-truth experiment (*‘accuracy’* dataset) was conducted in which two points of fixed distance to each other, the end points of a rigid calibration wand, were filmed underwater over various backgrounds using four cameras. In total, four repeated trials were incorporated for the accuracy estimation. Using our approach, we were able to retrieve both the 3D positions of the tracked calibration wand and the 3D trajectories of the cameras throughout the trials (Fig. 2). The known camera-to-camera distances within the camera array and the known length of the calibration wand allowed the calculation of respective per-frame root mean squared errors (RMSEs), 1.34 ± 0.79 cm (median error -0.14 ± 0.06 cm) for the camera-to-camera distances and 1.09 ± 0.47 cm (median error 0.14 ± 0.33 cm) for the calibration wand length. Further, we projected the triangulated 3D positions back to the original videos and computed the reprojection error as a RMSE of 8.56 ± 5.21 px (median error 3.53 ± 1.96 px).

**Figure 2.**
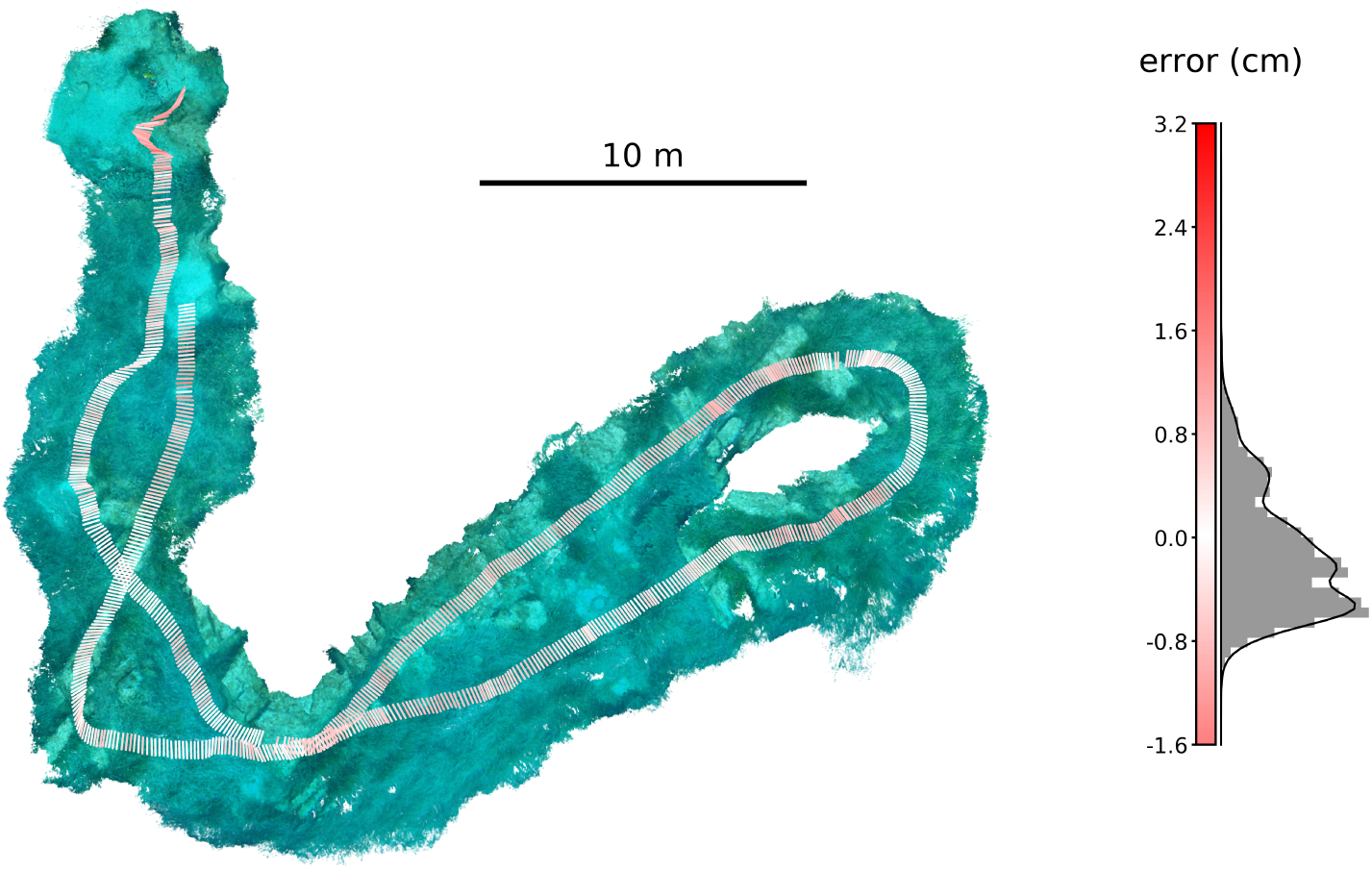
Accuracy validation. Top down view of one of the *‘accuracy’* dataset trials with the COLMAP dense reconstruction in the background (left). A calibration wand of 0.5 m was moved through the environment to create two trajectories with known per-frame distances (visualized as lines at a frequency of 3 Hz, the full temporal resolution of the trajectories is 30 Hz). This allowed the calculation of relative tracking errors as the difference of the triangulated calibration wand end-to-end distance from the its known length of 0.5 m, resulting in the shown error distribution (normalized histogram with probability density function, right). The per-frame tracking error is visualized as line color.

Trajectories were successfully obtained from large groups (*‘school’*), small groups (*‘mixed’*) and single individuals (*‘single’*). Additionally, the camera positions and corresponding environments through which the animals were moving were reconstructed. In case of the *‘single’* and *‘mixed’* datasets, the Mask R-CNN detection results were used to estimate fish body postures in 3D space by inferring spine points from the segmented pixel masks (Fig. 3). We computed the RMSEs of the camera-to-camera distances (1.28 cm *‘single’*, 1.28 cm *‘mixed’* and -0.15 cm *‘school’*) and reprojection errors (20.97 px *‘single’*, 7.77 px *‘mixed’* and 6.79 px *‘school’*) to assess the overall quality of the SfM reconstructions analogously to the calculation of reconstruction errors for the *‘accuracy’* dataset. The results of the accuracy estimations are listed in Tab. 2.

**Figure 3.**
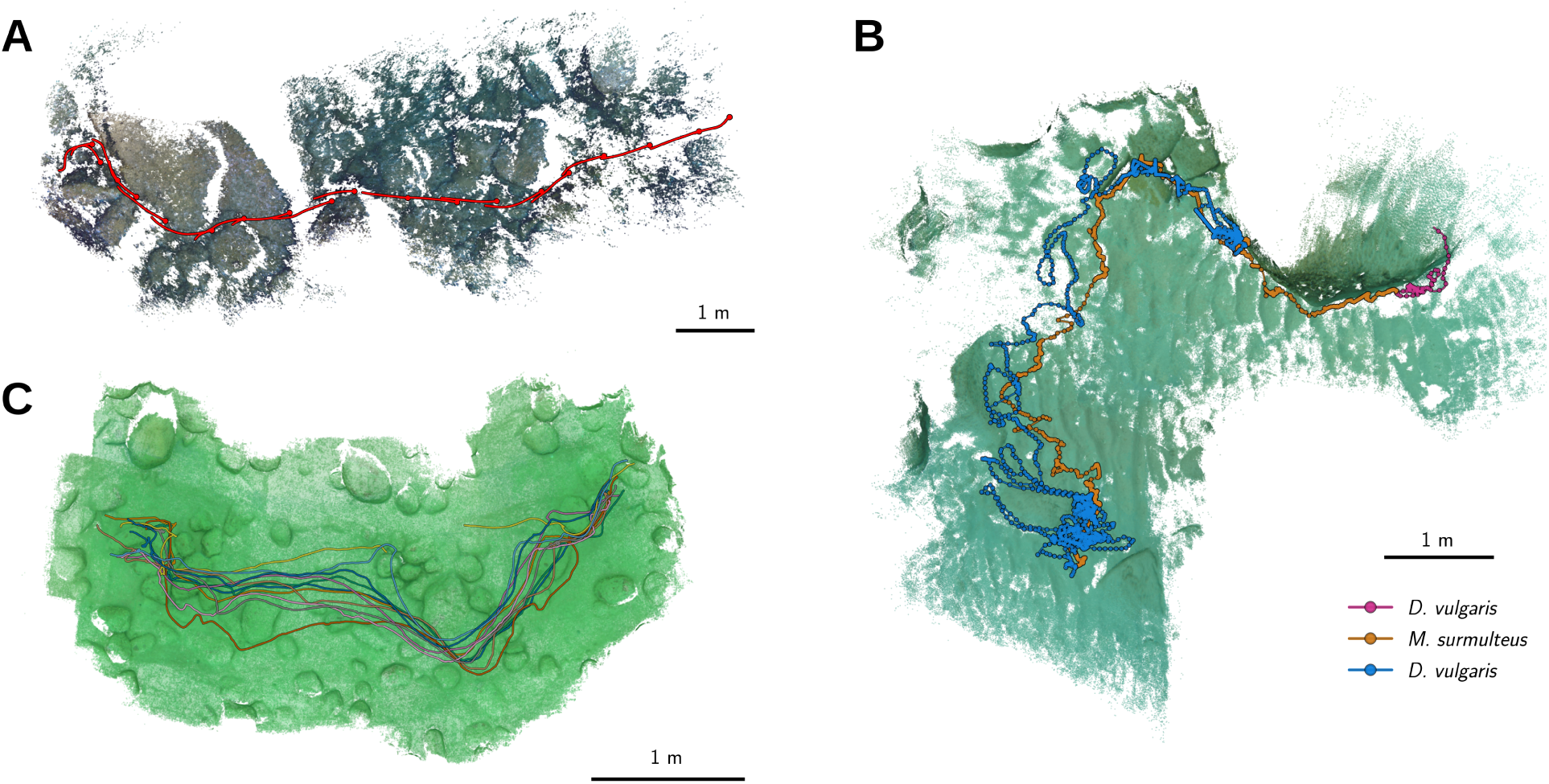
3D environments and animal trajectories. **A** Top down view of the *‘single’* dataset result. Red lines and dots show estimated spine poses and head positions of the tracked European eel (*C. conger*, visualized with one pose per second). The point cloud resulting from the COLMAP reconstruction is shown in the background. **B** Trajectories of *M. surmuletus* (orange) and *D. vulgaris* (purple/blue), and the dense point cloud resulting from the *‘mixed’* dataset. Dots highlight three positions per second, lines visualize the trajectories at full temporal resolution (30 Hz) over a duration of seven minutes. **B** Reconstruction results and trajectories of the *‘school’* dataset, visualizing the trajectories of a small school of *L. callipterus* in Lake Tanganyika. See additional files 1, 2 and 3 for high resolution images.

## Discussion

Here we demonstrate a novel approach to collect highly resolved 3D information of animal motion, including interactions with the physical environment, in aquatic ecosystems. Although being based on relatively advanced computational techniques, the open-source workflow we present requires little domain expertise and can be implemented with low-cost consumer grade cameras. The incorporation of these methods into an accessible framework will allow quantitative analyses of animal behaviour and ecology across systems, scales, and user groups. Our approach allows data collection while swimming, snorkelling, or with the aid of ROVs, making it appropriate for general usage with minimal investment into infrastructure, equipment, or training. Although analyses are computationally demanding, they can be achieved on an average GPU or free cloud-based computing services. The lack of high-end hardware therefore does not interfere with any of the steps required for this method.

Many alternative techniques for tracking of small aquatic animals do exist, however, they often have the considerable drawback of tagging and handling the animals or high infrastructure costs. This is a major barrier to implementation when animals are protected, difficult to catch, or too small to carry tags. In many marine protected areas all three of these factors apply, meaning that many existing approaches are inappropriate. Some of these drawbacks will be alleviated, for instance with improvements in telemetry-based approaches [36] that reduce tag size and increase range. Nevertheless, these techniques cannot simultaneously measure or reconstruct local topography and environmental factors. Although here we do not provide any analyses of environmental structure, this topographical information collected with our approach can be directly used to answer questions on e.g. habitat segmentation and environmental complexity [18, 32].

In highly complex social environments, encounters with numerous con- and heterospecifics can strongly affect behaviour and motion [57]. Using approaches that rely on tagging in these contexts will unavoidably miss or under-sample these interactions because not all individuals can ever be tagged in wild contexts. In contrast, our approach does not require animals to be handled or tagged, nor does specialized equipment need to be deployed in the desired tracking area. Moreover, because the object detection and segmentation approach can take any image input, it is not tied to one particular animal form or visual scene. Our approach can therefore be used even in demanding conditions such as high turbidity or low-light conditions, within certain limits. While it has a lower spatial range than telemetry, underwater filming comes as an unintrusive alternative, with higher spatial resolution possible when small animals are moving over small areas, or when animals are highly site-specific, for example damselfish or cichlids living in close association with coral or rocky reef [16, 53].

While our approach offers many benefits in terms of applicability and data acquisition, it also suffers from some limitations. From the accuracy tests it became apparent, that in cases were the background was composed of moving objects, such as macrophytes or debris, the tracking accuracy severely dropped. The SfM approach relies on the reconstructed components to be static, because environmental key-points are assumed to have the same location over time. Moving particles and objects will result in higher reconstruction errors, rendering our approach problematic e.g. when the filmed animals occupy most the captured images in case of very large fish schools. Complex environments, occlusions of the animals and highly variable lighting conditions are detrimental to the detectability of animals. Observations at greater depths may face similar problems due to the high absorption of red light, although, in this case, detectability could be alleviated through image augmentation approaches such as Sea-Thru [2]. The estimation of 3D animal poses strongly relies on accurate detections and can therefore be compromised by poorly estimated animal shapes during Mask R-CNN segmentation. In these cases, a less detailed approximation of the animals’ positions such as the mask centroids are favorable and can still be reliably employed as showcased in the *‘school’* example. The errors in estimating animal locations and poses can be partially explained by marginal detection errors of Mask R-CNN, but also by inaccuracies derived from trajectory triangulation using the SfM camera positions.

Aware of these error sources, users can actively incorporate accuracy metrics such as reprojection errors or relative camera reconstruction RMSEs into their analytical pathways by using our proposed method. This enables the assessment of the overall reconstruction quality and required fine scale resolution for the specific scientific demands. We were able to demonstrate with the *‘accuracy’* dataset, that the combination of SfM and object detection yields highly accurate trajectories of moving objects over large spatial scales (RMSE tracking error of 1.34 ± 0.79 cm, median error -0.14 ± 0.06 cm, reconstructed areas up to 500 m^2^) without prior manipulation of the underwater environment. Since these accuracy calculations are based on per-frame deviations from known distances, such as the length of a calibration wand or camera-to-camera distances in a stereo-camera setup, they are not suited for the assessment of large-scale SfM accuracy. However, rigorously ground-trouthing SfM is of general interest in the field of computer vision, and various benchmarks showcase the high precision of 3D reconstructions that can be achieved using current SfM pipelines [7, 34].

An additional requirement of our approach is associated with the need to annotate images and train object detection networks. However, this initial time investment is likely to be compensated by the time saved using automated detection and tracking of animals in most cases. For example, this allows the classification of behavioural states by quantifying the behavioural repertoire of the animal using unsupervised machine learning techniques [6, 55]. The incorporation of 3D trajectory data in motion analyses has already improved the understanding of the phenotype and development of animal behaviours [60]. In addition, 3D pose estimation can now be achieved for wild animals, enabling exact reconstruction of the entire animal [61]. There has been a shift in how animal movement is analyzed in light of computational ethological approaches [9, 44, 47], with patterns of motion able to be objectively disentangled, revealing the underlying behavioural syntax to the observer. Automated approaches based on video, or even audio, recordings may also overcome sensory limitations of other systems, allowing a better understanding of the sensory *umwelt* of study species [31] and also facilitate novel experimental designs [4, 47] that can tackle questions of the proximate and ultimate causality of behaviour [9, 44, 61]. These methods are gaining interest and sharply contrast with the traditional approach of trained specialists creating behavioural ethograms, but can usefully be combined and compared to gain further insight into the structure of animal behaviour, potentially generating a more objective and standardized approach to the field of behavioural studies [9].

In order to incorporate these novel techniques into more natural scenarios, we aim to present a complete tracking pipeline, guiding the user through each step after the initial field observation. From video synchronization, object annotation and detection to the final triangulation of animal trajectories, we provide a set of open-source utilities and scripts. Although we heavily rely on other open-source projects (COLMAP for SfM and Mask R-CNN for object segmentation), these specific approaches can be replaced with other implementations by solely adopting respective in- and output data formatting for specific needs. We found COLMAP and Mask R-CNN to be easily employed, as they are well documented, performant and purpose-oriented. However, many alternatives exist for both SfM and object detection, and the general approach of our pipeline is not limited to any particular implementation, thus future-proofing this approach as new and better methods are developed.

## Conclusions

Computational approaches to analyze behaviour, including automated tracking of animal groups, deep-learning, supervised, and unsupervised classification of behaviour, are areas of research that have been extensively developed in laboratory conditions over the past decade. These techniques, in combination with sound evolutionary and ecological theory, will characterize the next generation of breakthroughs in behavioural and movement science, yet are still difficult to achieve in natural contexts, and are unobtainable for many researchers due to implementation and infrastructure costs. Here we present a framework to enable the utilization of these cutting-edge approaches in aquatic ecosystems, at low-cost and for users of different backgrounds. Our proposed tracking method is flexible in both the conditions of use, and the study species being examined, vastly expanding our potential for examining non-model systems and species. In combination with the genomic revolution, allowing sequencing in a matter of days, state-of-the-art behavioural sequencing even under field conditions could revolutionize the field of movement ecology and evolutionary behavioural ecology. The approach we advocate here can further integrate the study of wild animal behaviour with modern techniques, facilitating an integrative understanding of movement in complex natural systems.

## Supporting information

Code_track_utility

Code_training_inference

Code_utils

Practical implementation guide

## List of abbreviations

GPS: Global Positioning System
PSAT: Pop-up satellite archival tag
ROV: Remotely operated underwater vehicle
Mask R-CNN: Mask and Region based Convolution Neural Network
SfM: Structure-from-Motion
RMSE: root-mean-square error
SCUBA: Self-contained underwater breathing apparatus

## Availability of data and material

The datasets used and/or analyzed during the current study are available from the corresponding author on request. All code used for analysis is open-source and either accessible online (*‘multiviewtracks’*, https://doi.org/10.5281/zenodo.3666726) or provided as additional files 4-6 (*‘TrackUtil’*, video synchronization, Mask R-CNN training and inference, trajectory assignment and pose estimation).

## Competing interests

The authors declare that they have no competing interests.

## Funding

This project was funded by the Deutsche Forschungsgemeinschaft (DFG, German research Foundation) under Germany’s Excellence Strategy - EXC 2117 - 422037984.

## Author’s contributions

FAF, PN and AJ collected the raw data in the field. PN and FAF wrote code for data acquisition and performed analyses. PN, FAF, and AJ wrote the manuscript. All authors agree on the standards of authorship put forth by this journal.

## Acknowledgments

We would like to thank the entire Department of Collective Behaviour at the University of Konstanz for their support in making this project possible. We thank Philip Fourmann, Myriam Knöpfle and Jessica Ruff for contributing the *‘single’* and *‘mixed’* species footage. Special thanks also to Hemal Naik, Simon Gingins and Eduardo Sampaio for their suggestions and helpful input. We sincerely thank Etienne Lein for his substantial assistance and support during data acquisition in the field. Further, we thank the COLMAP team for making it open-source and easily accessible.

## Supplementary Information

**Additional file 1 — *Tracking results of ‘single’ dataset* (.png)**

Top down view of *‘single’* results: dense COLMAP 3D reconstruction and trajectories of the tracked animal, *C. conger* (red).

**Additional file 2 — *Tracking results of ‘mixed’ dataset* (.png)**

Top down view of *‘mixed’* results: dense COLMAP 3D reconstruction and trajectories of the tracked animals, *M. surmuletus* (orange) and *D. vulgaris* (purple/blue).

**Additional file 3 — *Tracking results of ‘school’ dataset* (.png)**

Top down view of *‘school’* results: dense COLMAP 3D reconstruction and trajectories of the tracked animals, *L. callipterus*.

**Additional file 4 — *TrackUtil* (.zip)**

Python and Qt5 based GUI for image annotations, trajectory visualization and manual track corrections. This software will be made publicly available on GitHub at publication.

**Additional file 5 — *Mask R-CNN training and inference* (.zip)**

Our training regime for Mask R-CNN and inference on videos. For more information on Mask R-CNN, visit the original repository at https://github.com/matterport/Mask_RCNN.

**Additional file 6 — *Video synchronization, trajectory assignment and pose estimation* (.zip)**

Additional scripts for video synchronization, frame extraction, trajectory assignment and pose estimation. This software will be made publicly available on GitHub at publication.

**Additional file 7 — *A Practical Guide* (.pdf)**

A general guideline and comments regarding the implementation of our method.

## Notes

https://multiviewtracks.readthedocs.io/en/latest/

