## Supplementary material for "High-resolution animal tracking with integration of environmental information in aquatic systems": Code_track_utility: README.md

[![License: MIT](https://img.shields.io/badge/License-MIT-yellow.svg)](LICENSE)

# TrackUtil

**TrackUtil** features polygon-based video annotations for multiple classes.  
**TrackUtil** can be used to visualize, check and correct animal trajectories with optional video playback.  
**TrackUtil** aims to deliver an interactive workflow through a purpose-oriented GUI.

### Installation

The easiest way to get **TrackUtil** up and running across most platforms is using a [*miniconda*](https://docs.conda.io/en/latest/miniconda.html) environment for python3. Once conda is setup, a virtual environment can be created and activated by running the following code in a terminal (*Anaconda Promt* on Windows). Further, [*pip*](https://pypi.org/project/pip/) and [*git*](https://git-scm.com/) should be installed via conda to get **TrackUtil** and its requirements.

```bash
conda create -n trackutil python=3.7
conda activate trackutil
conda install git pip
```

Then, **TrackUtil** can be downloaded with *git*. Use *pip* to install the required packages from [REQUIREMENTS.txt](REQUIREMENTS.txt).

```bash
git clone https://gitlab.mpcdf.mpg.de/pnuehren/trackutil.git
cd trackutil
pip install -r requirements.txt
```

Assuming everything worked well, you can now run **TrackUtil** by executing the following command.

```bash
conda activate trackutil # not necessary if still active
python TrackUtil.py
```
### Basic usage

<kbd>File</kbd> menu - general input/output:
- <kbd>Load video</kbd> - load supported video files through the ffmpeg based OpenCV video reader.
- <kbd>Load tracks</kbd> - load tracks from .pkl file with correct structure.
- <kbd>Load annotations</kbd> - load annotations from previously saved .pkl file.
- <kbd>Save tracks</kbd> - save tracks to supported .pkl file with correct structure.
- <kbd>Save annotations</kbd> - save annotations to supported .pkl file.
- <kbd>Write to dataset</kbd> - write annotations from current video to .h5 dataset file for Mask R-CNN training.
- <kbd>Connect</kbd> - open connected instance of TrackUtil in a new window.

<kbd>Buffer</kbd> - start buffering from current frame, loads frames to memory for faster navigation.  
<kbd>Video OFF</kbd> <kbd>ON</kbd> - toggle video playback, use <kbd>Video OFF</kbd> for faster navigation.  
<kbd>Settings</kbd> - open settings widget. See [Settings](#settings) for details.  
<kbd>Annotator</kbd> - open annotator widget. See [Annotation](#annotation) for details.  
<kbd>Tracks</kbd> menu - User functions for track correction. See [Tracks](#tracks) for details.  
<kbd>Play</kbd> <kbd>Pause</kbd> or <kbd>Space</kbd> - start or pause the player.  
<kbd><<</kbd> <kbd>>></kbd> or <kbd>Left</kbd> <kbd>Right</kbd> - Single click or key: Previous or next frame. Keep mouse pressed for multiple frames.  
<kbd>Extend</kbd> <kbd>Hide</kbd> - Show or hide a finer slider for frame navigation that is scaled in seconds.

### Settings

- The player can visualize either trajectory poses or positions, or annotations. The *draw poses* option will be disabled, if the loaded tracks do not contain pose information. Otherwise, one of the *draw poses*, *positions* or *annotations* can be enabled. Enabling *draw annotations* will set **TrackUtil** into annotation mode, switching the <kbd>right-click</kbd> menu from the <kbd>Tracks</kbd> menu to the <kbd>Annotations</kbd> menu. See [Annotation](#annotation) for details.
- *Position radius* sets the size of the position visualizations in the player.
- *Number of individuals* sets the threshold of individuals used for the navigation function <kbd>More than n individuals</kbd>, see [Tracks](#tracks) for details.
- *Timeline cursor snapping* sets the sensitivity of cursor snapping to frames of interest (i.e. first or last frames of sub-trajectories) in the timeline widget. Snapping is disabled when set to 0 px.
- *Interpolate tracks* toggles the visualization of trajectories or their interpolated versions. Toggling the interpolation may need some time.
- *Synchronize connected* enables synchronization of connected **TrackUtil** instances via the sliders, the timeline widget and <kbd><<</kbd> <kbd>>></kbd> or <kbd>Left</kbd> <kbd>Right</kbd>. Synchronized playback is currently not supported.

### Tracks

- Tracks are visualized in the central player and the timeline widget, which can be opened by dragging the lower edge of the player.
- All displayed individuals (positions or poses in the player and time representations in the timeline) can be selected and deselected by <kbd>left-click</kbd>. To deselect all, press <kbd>Esc</kbd>.
- For exact frame navigation, use <kbd>Ctrl</kbd>+<kbd>left-click</kbd> in the timeline widget.
- The <kbd>Tracks</kbd> menu, also accessible in the status bar, can be opened by <kbd>right-click</kbd> in the player or the timeline.

<kbd>Tracks</kbd> menu - User functions for track correction:
- <kbd>Clear selection</kbd> or <kbd>Esc</kbd> - clear all selected individuals from selection.
- <kbd>Delete selected</kbd> or <kbd>Del</kbd> - delete all selected individual trajectories from tracks.
- <kbd>Split selected</kbd> or <kbd>Y</kbd> - split all selected individual trajectories between previous and current frame and add splitted trajectories to tracks as new individuals.
- <kbd>Switch selected</kbd> or <kbd>X</kbd> - switch the individuals of two selected trajectories, starting at current frame.
- <kbd>Merge selected</kbd> or <kbd>M</kbd> - merge all selected individual trajectories into one trajectory, only possible if selected trajectories do not share common frames.
- <kbd>Remove areas</kbd> - remove all detections that are contained by any annotated area. If all detections of one individual trajectory are removed, the individual trajectory will be completely removed as well. See [Annotation](#annotation) for details.
- <kbd>Navigate</kbd> submenu - Navigation functions:
  - <kbd>New identity</kbd> or <kbd>E</kbd> - go to the first frame of the individual trajectory, that is the next trajectory started after the current frame.
  - <kbd>Lost identity</kbd> or <kbd>Q</kbd> - go to the last frame of the individual trajectory, that is the last trajectory finished before the current frame.
  - <kbd>More than n individuals</kbd> or <kbd>O</kbd> - go to the first frame with more individuals than specified. See [Settings](#settings) on how to set the *Number of individuals* option.

### Annotation

- With the <kbd>Annotator</kbd> widget, **TrackUtil** can be used to annotate polygon-based masks for user-defined classes on multiple videos.
- Add a class by typing the class name in the text field and press <kbd>Return</kbd> or <kbd>Enter</kbd>. <kbd>Remove class</kbd> deletes the active class from the *Select class* menu, however, all annotations of the deleted class remain.
- Start annotating an object instance of the active class (*Select class*) on the specified occlusion level (*Set occlusion level*) by clicking <kbd>Start</kbd>. <kbd>left-click</kbd> in the player will add a point to the current annotation, <kbd>double-click</kbd> finishes the annotation. Starting an annotation will automatically set **TrackUtil** into annotation mode and disables trajectory visualizations. See [Settings](#settings) for details.
- Within one occlusion level, new annotations will occlude other annotations when drawn on top. Otherwise, annotations with higher occlusion levels will occlude annotations with lower occlusion levels.
- All displayed annotations can be selected and deselected by <kbd>left-click</kbd>. To deselect all, press <kbd>Esc</kbd>.
- The <kbd>Annotations</kbd> menu can also opened via <kbd>right-click</kbd> in the player or timeline widget.
- When saving annotations to a .pkl file with <kbd>File</kbd> menu > <kbd>Save annotations</kbd>, the file will also contain all previous annotations from other videos. It is possible to have all annotations within one .pkl file for a project, if the file names are unique for all videos. The .pkl file can be safely overridden with <kbd>Save annotations</kbd>, and allows continued annotation with <kbd>File</kbd> menu > <kbd>Load annotations</kbd>.
- The .pkl file only contains information about the annotated polygons and class names, for training (e.g. with Mask R-CNN), a .h5 dataset file has to be created. When an annotated video is loaded, <kbd>File</kbd> menu > <kbd>Write to dataset</kbd> creates or extends an existing dataset by adding the annotations of the current video. To save the full dataset, each annotated video has to be loaded and written to the same dataset by clicking override in the <kbd>Write to dataset</kbd> file dialog.

<kbd>Annotations</kbd> menu - Additional user functions for annotation:
- <kbd>Clear selection</kbd> or <kbd>Esc</kbd> - clear all selected annotations from selection.
- <kbd>Delete selected</kbd> or <kbd>Del</kbd> - delete all selected annotations.
- <kbd>Navigate</kbd> submenu - Navigation functions:
  - <kbd>Next annotation</kbd> or <kbd>E</kbd> - go to the next frame containing annotations.
  - <kbd>Previous annotation</kbd> or <kbd>Q</kbd> - go to the previous frame containing annotations.  

### License

See the [LICENSE](LICENSE) file for license rights and limitations (MIT).
