## Supplementary material for "High-resolution animal tracking with integration of environmental information in aquatic systems": Code_track_utility: TrackUtil.py

```
#!/usr/bin/env python3

import sys

from TrackUtil import TrackUtil, set_dark_style
from PySide2.QtWidgets import QApplication

def run(app):
    set_dark_style(app)
    tu = TrackUtil()
    app.exec_()
    tu.reader.running = False
    tu.reader.wait()
    return

if __name__=='__main__':
    app = QApplication()
    sys.exit(run(app))
```
