## Supplementary material for "High-resolution animal tracking with integration of environmental information in aquatic systems": Code_track_utility: Annotator.py

```
from PySide2.QtWidgets import QWidget, QVBoxLayout, QHBoxLayout, QCheckBox, QPushButton, QLineEdit, QComboBox, QCompleter, QLabel, QSpinBox
from PySide2.QtGui import QRegExpValidator
from PySide2.QtCore import Signal, QObject
from PySide2 import QtCore

from .QtUtils import NamedSpinBox, BorderlessWindow

class AnnotatorSignal(QObject):
    annotating = Signal(int)
    class_name = Signal(str)
    remove = Signal(str)

class Annotator(BorderlessWindow):

    def __init__(self, menu_annotation):
        super().__init__(main_window=False)
        self.setWindowTitle('Annotator')

        self.signal = AnnotatorSignal()

        self.classes = {}
        for class_name in self.classes:
            self.add_class(class_name, self.classes[class_name])
        max_level = max([len(self.classes),
                        0 if len(self.classes) == 0 else \
                        max([self.classes[class_name] for class_name in self.classes])])

        self._annotating = False

        self.edit_class = QLineEdit()
        self.edit_class.setCompleter(QCompleter([]))
        self.edit_class.setValidator(QRegExpValidator(QtCore.QRegExp("^\S*$")))
        self.edit_class.returnPressed.connect(self.add_class)

        self.occlusion_level = QSpinBox()
        self.occlusion_level.valueChanged.connect(self.change_level)

        self.select_class = QComboBox()
        self.select_class.currentIndexChanged.connect(self.set_level)

        self.button_remove_class = QPushButton('Remove class')
        self.button_remove_class.clicked.connect(self.remove_class)

        self.button_draw = QPushButton('Start')
        self.button_draw.clicked.connect(lambda v: setattr(self, 'annotating', v))

        self.button_annotations = QPushButton('Annotations')
        self.button_annotations.setMenu(menu_annotation)

        layout = self.layout()
        layout.addWidget(self.button_annotations)
        layout.addWidget(QLabel('Add class'))
        layout.addWidget(self.edit_class)
        layout.addWidget(QLabel('Select class'))
        layout.addWidget(self.select_class)
        layout.addWidget(self.button_remove_class)
        layout.addWidget(NamedSpinBox(self.occlusion_level, 'Set occlusion level', 0, 0, max_level, True))
        layout.addWidget(self.occlusion_level)
        layout.addWidget(self.button_draw)
        self.setLayout(layout)

    @property
    def annotating(self):
        return self._annotating

    @annotating.setter
    def annotating(self, value):
        if not self._annotating:
            text = self.select_class.currentText().split(' ')
            if len(text) <= 1:
                return
            class_name = text[0]
            occlusion_level = int(text[1].replace('(', '').replace(')', ''))
            if class_name in self.classes:
                self._annotating = not self._annotating
                self.signal.annotating.emit(occlusion_level)
                self.signal.class_name.emit(class_name)
        else:
            self._annotating = not self._annotating
            self.signal.annotating.emit(-1)
        self.button_draw.setText('Finish' if self.annotating else 'Start')

    def add_class(self, class_name=None, occlusion_level=None, silent=False):
        if class_name is None:
            class_name = self.edit_class.text()
        if class_name == '':
            return
        if class_name not in self.classes:
            if occlusion_level is None:
                occlusion_level = self.occlusion_level.value()
            self.select_class.addItem('{} ({})'.format(class_name, occlusion_level))
            self.classes[class_name] = occlusion_level
            self.occlusion_level.setMaximum(max([len(self.classes), max([self.classes[class_name] for class_name in self.classes])]))
            idx = [idx for idx in range(self.select_class.count()) if self.select_class.itemText(idx).split(' ')[0] == class_name][0]
            self.select_class.setCurrentIndex(idx)
        self.edit_class.setText('')
        if not silent:
            self.signal.class_name.emit(class_name)

    def remove_class(self, class_name=None, silent=False):
        if class_name is None:
            text = self.select_class.currentText().split(' ')
            if len(text) <= 1:
                return
            class_name = text[0]
        if class_name in self.classes:
            idx = [idx for idx in range(self.select_class.count()) if self.select_class.itemText(idx).split(' ')[0] == class_name][0]
            self.select_class.removeItem(idx)
            self.classes.pop(class_name)
        if not silent:
            self.signal.remove.emit(class_name)

    def change_level(self, occlusion_level):
        text = self.select_class.currentText().split(' ')
        if len(text) <= 1:
            return
        class_name = text[0]
        if class_name in self.classes:
            idx = [idx for idx in range(self.select_class.count()) if self.select_class.itemText(idx).split(' ')[0] == class_name][0]
            self.select_class.setItemText(idx, '{} ({})'.format(class_name, occlusion_level))

    def set_level(self):
        text = self.select_class.currentText().split(' ')
        if len(text) > 1:
            class_name = text[0]
            occlusion_level = int(text[1].replace('(', '').replace(')', ''))
            self.occlusion_level.setValue(occlusion_level)
```
