## Supplementary material for "High-resolution animal tracking with integration of environmental information in aquatic systems": Code_track_utility: Drawing.py

```
import numpy as np

from PySide2 import QtCore, QtGui

def draw_pose(scene, spine, rad, identity, color, selected, hovered):
    drawing = []
    for i, (pt, r) in enumerate(zip(spine[::-1], rad[::-1])):
        drawing.append(scene.addEllipse(*(pt - r), r * 2, r * 2,
                                              pen=QtGui.QPen(QtGui.QColor(*(255, 255, 255) if selected else color, 255), 0.5),
                                              brush=QtGui.QBrush(QtGui.QColor(*color, 100 if i == len(spine) - 1 or hovered or selected else 0))))
    pts = [QtCore.QPointF(*pt) for pt in spine]
    polygon = QtGui.QPolygonF(pts)
    line = QtGui.QPainterPath()
    line.addPolygon(polygon)
    drawing.append(scene.addPath(line, pen=QtGui.QPen(QtGui.QColor(*(255, 255, 255) if selected else color), 1.5),
                                       brush=QtGui.QBrush(QtGui.QColor(*color, 0))))
    for i, pt in enumerate(spine):
        r = 2 if hovered else 1
        drawing.append(scene.addEllipse(*(pt - r), r * 2, r * 2,
                                              pen=QtGui.QPen(QtGui.QColor(*(255, 255, 255) if selected else color), 1 if hovered else 0.5),
                                              brush=QtGui.QBrush(QtGui.QColor(*color if selected else (255, 255, 255), 255))))
    for item in drawing:
        item.setToolTip(str(identity))
    return drawing

def draw_position(scene, x, y, size, identity, color, selected, hovered):
    drawing = []
    pt = np.array([x, y])
    drawing.append(scene.addEllipse(*(pt - size), size * 2, size * 2,
                                          pen=QtGui.QPen(QtGui.QColor(*(255, 255, 255) if selected else color, 255 if selected else 0), 0.5),
                                          brush=QtGui.QBrush(QtGui.QColor(*color, 100 if selected else 0))))
    r = size / 2 if hovered else size / 3
    r = 2 if r < 2 else r
    drawing.append(scene.addEllipse(*(pt - r), r * 2, r * 2,
                                          pen=QtGui.QPen(QtGui.QColor(*color if selected else (255, 255, 255)), 1 if hovered else 0.5),
                                          brush=QtGui.QBrush(QtGui.QColor(*(255, 255, 255) if selected else color, 255))))
    for item in drawing:
        item.setToolTip(str(identity))
    return drawing

def draw_polygon(scene, polygon, color, selected=False, hovered=False, class_name=None):
    drawing = []
    line = QtGui.QPainterPath()
    line.addPolygon(polygon)
    if selected:
        drawing.append(scene.addRect(polygon.boundingRect(),
                       pen=QtGui.QPen(QtGui.QColor(255, 255, 255, 255), 1.5),
                       brush=QtGui.QBrush(QtGui.QColor.fromHsv(*color, 50))))
    elif hovered:
        drawing.append(scene.addRect(polygon.boundingRect(),
                       pen=QtGui.QPen(QtGui.QColor.fromHsv(*color, 255), 1.5),
                       brush=QtGui.QBrush(QtGui.QColor(0, 0, 0, 0))))
    drawing.append(scene.addPath(line, pen=QtGui.QPen(QtGui.QColor.fromHsv(*color, 0 if polygon.isClosed() else 150), 1.5),
                                       brush=QtGui.QBrush(QtGui.QColor.fromHsv(*color, 100 if polygon.isClosed() else 50))))
    for pt in polygon:
        drawing.append(scene.addEllipse(*(pt - QtCore.QPointF(0.75, 0.75)).toTuple(), 1.5, 1.5,
                                              pen=QtGui.QPen(QtGui.QColor.fromHsv(*(0, 0, 255) if selected or hovered else color, 255), 0.5),
                                              brush=QtGui.QBrush(QtGui.QColor.fromHsv(*(0, 0, 255) if selected or hovered else color, 255))))
    if class_name is not None:
        text = scene.addText(class_name)
        text.setPos(polygon.boundingRect().topLeft())
        text.setDefaultTextColor(QtGui.QColor.fromHsv(*color, 255))
        drawing.append(text)
    return drawing
```
