## Supplementary material for "High-resolution animal tracking with integration of environmental information in aquatic systems": Code_track_utility: History.py

```
from PySide2.QtWidgets import QScrollArea, QWidget, QVBoxLayout, QPushButton
from PySide2.QtGui import QPalette, QColor
from PySide2 import QtCore

from .QtUtils import named_cb, BorderlessWindow, ClickableLabel

from PySide2.QtCore import Signal, QObject

class HistorySignal(QObject):
    history = Signal(list)

class History(BorderlessWindow):

    def __init__(self):
        super().__init__(main_window=False, height_resizable=True)

        self.setWindowTitle('History')

        self.scroll_area = QScrollArea()
        self.scroll_area.setHorizontalScrollBarPolicy(QtCore.Qt.ScrollBarAlwaysOff)
        self.scroll_area.setWidgetResizable(True)
        palette = QPalette()
        self.scroll_area.setBackgroundRole(palette.Base)

        scroll_bar = self.scroll_area.verticalScrollBar()
        scroll_bar.rangeChanged.connect(lambda x, y: scroll_bar.setValue(y))

        self.container = QWidget()
        self.container_layout = QVBoxLayout()
        self.container_layout.addStretch(-1)
        self.container.setLayout(self.container_layout)
        self.scroll_area.setWidget(self.container)

        self.button_make_backup = QPushButton('Save progress')
        self.button_load_backup = QPushButton('Load progress')

        layout = self.layout()
        layout.addWidget(self.scroll_area)
        layout.addWidget(self.button_load_backup)
        layout.addWidget(self.button_make_backup)
        self.setLayout(layout)

        self.events = []
        self.event_labels = []
        self.checkpoint = 0

        self.signal = HistorySignal()

    def add_event(self, kind, selected, frame):
        self.revert_to_checkpoint()
        event = '[{}] {}'.format(kind, ', '.join([str(i) for i in selected]))
        event_label = ClickableLabel(event, str(len(self.events)))
        self.events.append((kind, selected, frame))
        self.event_labels.append(event_label)
        event_label.signal.activated.connect(self.set_event)
        event_label.signal.deactivated.connect(self.unset_event)
        self.container_layout.insertWidget(self.container_layout.count() - 1, event_label)
        self.checkpoint = len(self.events)

    def set_event(self, active_idx):
        for idx, event_label in enumerate(self.event_labels):
            if idx < int(active_idx):
                event_label.set_active()
            elif idx > int(active_idx):
                event_label.set_inactive()
        self.checkpoint = int(active_idx) + 1
        self.signal.history.emit(self.events[:self.checkpoint])

    def unset_event(self, active_idx):
        for idx, event_label in enumerate(self.event_labels):
            event_label.set_active()
        self.checkpoint = len(self.events)
        self.signal.history.emit(self.events)

    def clear_events(self):
        self.event_labels = []
        for idx in range(len(self.events)):
            event_label = self.container_layout.takeAt(0)
            event_label.widget().deleteLater()
        self.events = []

    def revert_to_checkpoint(self):
        if len(self.event_labels) > 0:
            self.event_labels[self.checkpoint - 1].set_active()
        if len(self.events) > self.checkpoint:
            self.event_labels = self.event_labels[:self.checkpoint]
            for idx in range(len(self.events) - self.checkpoint):
                event_label = self.container_layout.takeAt(self.checkpoint)
                event_label.widget().deleteLater()
            self.events = self.events[:self.checkpoint]
```
