## Supplementary material for "High-resolution animal tracking with integration of environmental information in aquatic systems": Code_track_utility: Player.py

```
from PySide2.QtWidgets import QGraphicsEllipseItem, QGraphicsPathItem, QMenu
from PySide2.QtGui import QPixmap, QImage, QCursor
from PySide2 import QtCore, QtGui, QtWidgets
from PySide2.QtCore import Signal, QObject

import numpy as np
import cv2

from .Utils import diverging_hsv
from .Drawing import *

class PlayerSignal(QObject):
    identity = Signal(int)
    hovered = Signal(int)
    closed = Signal(bool)
    annotations = Signal(dict)
    empty = Signal()

class Player(QtWidgets.QGraphicsView):

    def __init__(self, menu_trajectory, menu_annotation):
        super().__init__()
        self.signal = PlayerSignal()
        self._zoom = 0
        self._empty = True
        self._annotating = False
        self._scene = QtWidgets.QGraphicsScene(self)
        self._photo = QtWidgets.QGraphicsPixmapItem()
        self._scene.addItem(self._photo)
        self.fitted = True
        self.set_annotations(None)
        self.setScene(self._scene)
        self.setTransformationAnchor(QtWidgets.QGraphicsView.AnchorUnderMouse)
        self.setResizeAnchor(QtWidgets.QGraphicsView.AnchorUnderMouse)
        self.setVerticalScrollBarPolicy(QtCore.Qt.ScrollBarAlwaysOff)
        self.setHorizontalScrollBarPolicy(QtCore.Qt.ScrollBarAlwaysOff)
        self.setBackgroundBrush(QtGui.QBrush(QtGui.QColor(15, 15, 15)))
        self.setFrameShape(QtWidgets.QFrame.NoFrame)
        self.setMouseTracking(True)
        self.class_name = None
        self.menus_toggle = True
        self.menu_trajectory = menu_trajectory
        self.menu_annotation = menu_annotation

    @property
    def annotating(self):
        return self._annotating

    @annotating.setter
    def annotating(self, value):
        self._annotating = not self._annotating
        if self._annotating:
            if value >= 0:
                self.occlusion_level = value
                while self.occlusion_level >= len(self.polygons):
                    self.add_level()
            self.signal.hovered.emit(-1)
            self.annotation = []
        self.set_cursor()

    def add_level(self):
        self.polygons.append([])
        self.occluded.append([])
        self.colors.append([])
        self.class_names.append([])
        self.hovered.append([])
        self.selected.append([])

    def set_cursor(self):
        if self.annotating:
            self.viewport().setCursor(QtCore.Qt.CrossCursor)
        else:
            self.viewport().setCursor(QtCore.Qt.ArrowCursor)

    def prepare_annotations(self):
        annotations = {}
        for occlusion_level, (polygons, occluded, class_names, colors) in enumerate(zip(self.polygons, self.occluded, self.class_names, self.colors)):
            annotations[occlusion_level] = {'polygons': [], 'occluded': [], 'class_names': [], 'colors': []}
            annotations[occlusion_level]['polygons'] = [np.array([(pt.x(), pt.y()) for pt in polygon]) for polygon in polygons]
            annotations[occlusion_level]['occluded'] = [np.array([(pt.x(), pt.y()) for pt in polygon]) for polygon in occluded]
            annotations[occlusion_level]['class_names'] = class_names
            annotations[occlusion_level]['colors'] = colors
        return annotations

    def set_annotations(self, annotations):
        self.drawing = None
        self.occlusion_level = 0
        self.annotation = []
        self.hovered = []
        self.selected = []
        self.polygons = []
        self.occluded = []
        self.colors = []
        self.class_names = []
        if annotations is None:
            self.add_level()
        else:
            for occlusion_level in annotations:
                self.add_level()
                for polygon, occluded, class_name, color in zip(*annotations[occlusion_level].values()):
                    self.polygons[occlusion_level].append(QtGui.QPolygonF([QtCore.QPointF(*pt) for pt in polygon]))
                    self.occluded[occlusion_level].append(QtGui.QPolygonF([QtCore.QPointF(*pt) for pt in occluded]))
                    self.class_names[occlusion_level].append(class_name)
                    self.colors[occlusion_level].append(color)
                    self.hovered[occlusion_level].append(False)
                    self.selected[occlusion_level].append(False)
            self.draw_annotations()

    def hover_annotations(self, pos):
        changed = False
        for occlusion_level, polygons in enumerate(self.occluded):
            for idx, p in enumerate(polygons):
                if p.boundingRect().contains(pos) and not self.hovered[occlusion_level][idx]:
                    self.hovered[occlusion_level][idx] = True
                    changed = True
                elif not p.boundingRect().contains(pos) and self.hovered[occlusion_level][idx]:
                    self.hovered[occlusion_level][idx] = False
                    changed = True
        if changed:
            self.draw_annotations()

    def select_annotations(self, pos):
        changed = False
        for occlusion_level, polygons in enumerate(self.occluded):
            for idx, p in enumerate(polygons):
                if p.boundingRect().contains(pos):
                    self.selected[occlusion_level][idx] = not self.selected[occlusion_level][idx]
                    changed = True
        if changed:
            self.draw_annotations()

    def delete_annotations(self):
        changed = False
        for occlusion_level, polygons in enumerate(self.selected):
            changed = True if sum(self.selected[occlusion_level]) > 0 else False
            self.polygons[occlusion_level] = [polygon for polygon, s in zip(self.polygons[occlusion_level], self.selected[occlusion_level]) if not s]
            self.occluded[occlusion_level] = [occluded for occluded, s in zip(self.occluded[occlusion_level], self.selected[occlusion_level]) if not s]
            self.class_names[occlusion_level] = [class_name for class_name, s in zip(self.class_names[occlusion_level], self.selected[occlusion_level]) if not s]
            self.colors[occlusion_level] = [color for color, s in zip(self.colors[occlusion_level], self.selected[occlusion_level]) if not s]
            self.hovered[occlusion_level] = [hovered for hovered, s in zip(self.hovered[occlusion_level], self.selected[occlusion_level]) if not s]
            self.selected[occlusion_level] = [selected for selected, s in zip(self.selected[occlusion_level], self.selected[occlusion_level]) if not s]
        if changed:
            self.draw_annotations()
            self.signal.annotations.emit(self.prepare_annotations())
        else:
            self.signal.empty.emit()

    def draw_annotations(self):
        self.clear_drawings()
        for occlusion_level, polygons in enumerate(self.occluded):
            for idx, p in enumerate(polygons):
                draw_polygon(self._scene, p, self.colors[occlusion_level][idx],
                             class_name=self.class_names[occlusion_level][idx],
                             selected=self.selected[occlusion_level][idx],
                             hovered=self.hovered[occlusion_level][idx])

    def hasPhoto(self):
        return not self._empty

    def fitInView(self, scale=True, upscale=False):
        rect = QtCore.QRectF(self._photo.pixmap().rect())
        if not rect.isNull():
            self.setSceneRect(rect)
            if self.hasPhoto():
                unity = self.transform().mapRect(QtCore.QRectF(0, 0, 1, 1))
                if not upscale:
                    self.scale(1 / unity.width(), 1 / unity.height())
                viewrect = self.viewport().rect()
                scenerect = self.transform().mapRect(rect)
                if 0 in [viewrect.width(), scenerect.width(), viewrect.height(), scenerect.height()]:
                    return
                factor = min(viewrect.width() / scenerect.width(),
                             viewrect.height() / scenerect.height())
                if not upscale or self.fitted:
                    self.scale(factor, factor)
                    self.fitted = True
                elif upscale and factor >= 1:
                    self.scale(factor, factor)
                    self.fitted = True
            self._zoom = 0

    def setPhoto(self, pixmap=None):
        if pixmap and not pixmap.isNull():
            self._empty = False
            self.setDragMode(QtWidgets.QGraphicsView.ScrollHandDrag)
            self._photo.setPixmap(pixmap)
        else:
            self._empty = True
            self.setDragMode(QtWidgets.QGraphicsView.ScrollHandDrag)
            self._photo.setPixmap(QtGui.QPixmap())
        self.fitInView(upscale=True)

    def clear_drawings(self):
        for item in self._scene.items()[:-1]:
            self._scene.removeItem(item)

    def wheelEvent(self, event):
        if event.angleDelta().y() > 0:
            factor = 1.25
            self._zoom += 1
        else:
            factor = 0.8
            self._zoom -= 1
        if self._zoom > 0:
            self.scale(factor, factor)
            self.fitted = False
        else:
            self._zoom = 0
            self.fitInView()

    def mousePressEvent(self, event):
        if event.button() == QtCore.Qt.MouseButton.LeftButton and not self.annotating:
            super().mousePressEvent(event)
            item = self._scene.itemAt(self.mapToScene(event.pos()), QtGui.QTransform())
            if item is not None:
                if item is not self._photo and item.toolTip() != '':
                    self.signal.identity.emit(int(item.toolTip()))
                else:
                    self.select_annotations(self.mapToScene(event.pos()))
        elif event.button() == QtCore.Qt.MouseButton.RightButton and not self.annotating:
            super().mousePressEvent(event)
            if self.menus_toggle:
                self.menu_trajectory.popup(event.globalPos())
            else:
                self.menu_annotation.popup(event.globalPos())
        elif event.button() == QtCore.Qt.MouseButton.LeftButton and self.annotating:
            if len(self.annotation) == 0:
                self.colors[self.occlusion_level].append(diverging_hsv(self.colors))
                self.selected[self.occlusion_level].append(False)
                self.hovered[self.occlusion_level].append(False)
                self.class_names[self.occlusion_level].append(self.class_name)
            self.annotation.append(self.mapToScene(event.pos()))

    def mouseDoubleClickEvent(self, event):
        super().mousePressEvent(event)
        if event.button() == QtCore.Qt.MouseButton.LeftButton and self.annotating:
            pts = self.annotation[:-1] + [self.mapToScene(event.pos()), self.annotation[0]]
            polygon = QtGui.QPolygonF(pts)
            changed = False
            for occlusion_level, polygons in enumerate(self.occluded):
                for idx, p in enumerate(polygons):
                    if p.intersects(polygon):
                        changed = True
                        if occlusion_level > self.occlusion_level:
                            polygon = polygon.subtracted(self.occluded[occlusion_level][idx])
                        else:
                            self.occluded[occlusion_level][idx] = p.subtracted(polygon)
            self.polygons[self.occlusion_level].append(polygon)
            self.occluded[self.occlusion_level].append(polygon)
            self.signal.annotations.emit(self.prepare_annotations())
            self.signal.closed.emit(True)
            self.drawing = None
            self.annotation = []
            self.draw_annotations()

    def mouseMoveEvent(self, event):
        super().mouseMoveEvent(event)
        item = self._scene.itemAt(self.mapToScene(event.pos()), QtGui.QTransform())
        if not self.annotating:
            if item is not None and item is not self._photo and item.toolTip() != '':
                self.signal.hovered.emit(int(item.toolTip()))
            else:
                self.hover_annotations(self.mapToScene(event.pos()))
                self.signal.hovered.emit(-1)
        else:
            if len(self.annotation) > 0:
                pts = self.annotation + [self.mapToScene(event.pos())]
                polygon = QtGui.QPolygonF(pts)
                if self.drawing is not None:
                    for item in self.drawing:
                        self._scene.removeItem(item)
                self.drawing = draw_polygon(self._scene, polygon, self.colors[self.occlusion_level][-1])

    def resizeEvent(self, event):
        self.fitInView(upscale=True)
        super().resizeEvent(event)

    def mouseReleaseEvent(self, event):
        super().mouseReleaseEvent(event)
        self.set_cursor()

    def enterEvent(self, event):
        super().enterEvent(event)
        self.set_cursor()
```
