## Supplementary material for "High-resolution animal tracking with integration of environmental information in aquatic systems": Code_track_utility: QtUtils.py

```
from PySide2.QtWidgets import QPushButton, QCheckBox, QLabel, QHBoxLayout, QVBoxLayout, QSpacerItem, QSizePolicy, QWidget, QSlider, QStyle, QFrame, QButtonGroup, QDesktopWidget, QApplication
from PySide2.QtGui import QPalette, QColor, QCursor
from PySide2 import QtCore, QtGui
from PySide2.QtCore import Signal, QObject

from multiprocessing import Pool, cpu_count
import numpy as np
import sys

def named_cb(label, checked):
    cb = QCheckBox(label)
    if checked:
        cb.setCheckState(QtCore.Qt.Checked)
    return cb

class NamedSpinBox(QWidget):

    def __init__(self, sb, label, value, minimum, maximum, vertical=False):
        super().__init__()
        sb.setMinimum(minimum)
        sb.setMaximum(maximum)
        sb.setValue(value)
        label = QLabel(label)
        if vertical:
            layout = QVBoxLayout()
        else:
            sb.setSizePolicy(QSizePolicy.Fixed, QSizePolicy.Fixed)
            layout = QHBoxLayout()
        layout.addWidget(label)
        layout.addWidget(sb)
        layout.setMargin(0)
        self.setLayout(layout)

class LabelSignal(QObject):
    activated = Signal(str)
    deactivated = Signal(str)

class ClickableLabel(QLabel):

    def __init__(self, text, description=None):
        super().__init__()
        self.setText(text)
        self.setWordWrap(True)
        self.setAutoFillBackground(True)
        self.signal = LabelSignal()
        self.description = '' if description is None else description
        self._is_selected = False

    def mousePressEvent(self, event):
        if QtCore.Qt.MouseButton.LeftButton == event.button():
            self._is_selected = not self._is_selected
            palette = QPalette()
            if self._is_selected:
                self.set_selected()
                self.signal.activated.emit(self.description)
            else:
                self.signal.deactivated.emit(self.description)
                self.set_active()

    def set_selected(self):
        palette = QPalette()
        palette.setColor(QPalette.Background, QColor(15, 15, 15))
        palette.setColor(QPalette.WindowText, QColor(42, 130, 218))
        self.setPalette(palette)
        self.setBackgroundRole(palette.Background)

    def set_active(self):
        self._is_selected = False
        palette = QPalette()
        palette.setColor(QPalette.Background, QColor(15, 15, 15))
        palette.setColor(QPalette.WindowText, QtCore.Qt.white)
        self.setPalette(palette)
        self.setBackgroundRole(palette.Background)

    def set_inactive(self):
        self._is_selected = False
        palette = QPalette()
        palette.setColor(QPalette.Background, QColor(15, 15, 15))
        palette.setColor(QPalette.WindowText, QColor(100, 100, 100))
        self.setPalette(palette)
        self.setBackgroundRole(palette.Background)

class JumpSlider(QSlider):

    def __init__(self, *args, **kwargs):
        super().__init__(*args, **kwargs)

    def mousePressEvent(self, event):
        self.setValue(QStyle.sliderValueFromPosition(self.minimum(), self.maximum(), event.x(), self.width()))

    def mouseMoveEvent(self, event):
        self.setValue(QStyle.sliderValueFromPosition(self.minimum(), self.maximum(), event.x(), self.width()))

class HoldButton(QPushButton):

    def __init__(self, *args, **kwargs):
        super().__init__(*args, **kwargs)
        self.setAutoRepeat(True)
        self.setAutoRepeatDelay(200)
        self.setAutoRepeatInterval(200)
        self.clicked.connect(self.handleClicked)
        self._state = False

    def handleClicked(self):
        if self.isDown():
            if not self._state:
                self._state = True
                self.setAutoRepeatInterval(50)
        elif self._state:
            self._state = False
            self.setAutoRepeatInterval(200)

def slider_set_value(slider, value):
    slider.blockSignals(True)
    slider.setValue(value)
    slider.blockSignals(False)

def point_in_polygons(point):
    in_polygons = np.any([QtGui.QPolygonF([QtCore.QPointF(*pt) for pt in polygon]).containsPoint(QtCore.QPointF(*point),
                                                                                                 QtCore.Qt.OddEvenFill) \
                          for polygon in polygons])
    return in_polygons

def init_polygons(p):
    global polygons
    polygons = p

def points_in_polygons(pts, polygons):
    with Pool(processes=cpu_count(), initializer=init_polygons, initargs=(polygons, )) as pool:
        in_polygons = pool.map(point_in_polygons, pts)
        pool.close()
        pool.join()
    return np.array(in_polygons)

def set_dark_style(app):
    app.setStyle('Fusion')
    app.setStyleSheet('QMenu::separator { height: 1px; background: rgb(53, 53, 53); margin-left: 0px; margin-right: 0px; }')
    palette = QPalette()
    palette.setColor(QPalette.Window, QColor(53, 53, 53))
    palette.setColor(QPalette.WindowText, QtCore.Qt.white)
    palette.setColor(QPalette.Base, QColor(15, 15, 15))
    palette.setColor(QPalette.AlternateBase, QColor(53, 53, 53))
    palette.setColor(QPalette.ToolTipBase, QtCore.Qt.white)
    palette.setColor(QPalette.ToolTipText, QColor(53, 53, 53))
    palette.setColor(QPalette.Text, QtCore.Qt.white)
    palette.setColor(QPalette.Button, QColor(53, 53, 53))
    palette.setColor(QPalette.ButtonText, QtCore.Qt.white)
    palette.setColor(QPalette.BrightText, QtCore.Qt.red)
    palette.setColor(QPalette.Link, QColor(42, 130, 218))
    palette.setColor(QPalette.Highlight, QColor(42, 130, 218))
    palette.setColor(QPalette.HighlightedText, QtCore.Qt.black)
    app.setPalette(palette)

class TransparentWindow(QFrame):

    def __init__(self, main_window=True):
        super().__init__()
        self._is_main_window = main_window
        self.setWindowFlags(QtCore.Qt.Tool |QtCore.Qt.FramelessWindowHint | QtCore.Qt.WindowStaysOnTopHint)
        self.setObjectName('transparentwindow')
        self.setStyleSheet('#transparentwindow { border: 2px solid rgb(42,130,218); }')
        self.setWindowOpacity(0.5)

    def mouseMoveEvent(self, event):
        self.hide()

    def mousePressEvent(self, event):
        self.hide()

class BorderlessWindow(QFrame):

    def __init__(self, main_window=True, height_resizable=False):
        super().__init__()
        self._is_main_window = main_window
        self._height_resizable = True if main_window or height_resizable else False
        self.setWindowFlags(QtCore.Qt.FramelessWindowHint)
        if (sys.platform == 'linux' or sys.platform == 'linux2') and not self._is_main_window:
            self.setWindowFlags(self.windowFlags() ^ QtCore.Qt.WindowStaysOnTopHint)
        self.setObjectName('borderlesswindow')
        self.setStyleSheet('#borderlesswindow { border: 1px solid rgb(70,70,70); }')
        self.setMouseTracking(True)
        self.init_window()
        self._top = False
        self._left = False
        self._bottom = False
        self._right = False
        self._handle = False
        self._drag = False
        self._saved_geometry = None
        self._pos_before_drag = None
        self._pos_starting_drag = None
        self._double_click_interval = QApplication.instance().doubleClickInterval()
        self._click_timer = QtCore.QTimer(self)
        self._click_timer.timeout.connect(self.single_click)
        self._click_timer.setSingleShot(True)
        self._drag_timer = QtCore.QTimer(self)
        self._drag_timer.timeout.connect(self.drag)
        self._drag_timer.setSingleShot(True)
        self._window = TransparentWindow()

    def colored_button(self, color):
        pal = self.palette()
        button = QPushButton()
        pal.setColor(QPalette.Button, color)
        button.setPalette(pal)
        button_size = (self.available_geometry().height()  * 9.5 / 1080)
        button.setFixedWidth(button_size)
        button.setFixedHeight(button_size)
        return button

    def set_on_top(self):
        if (sys.platform == 'linux' or sys.platform == 'linux2') and not self._is_main_window:
            return
        self.setWindowFlags(self.windowFlags() ^ QtCore.Qt.WindowStaysOnTopHint)
        pal = self.palette()
        pal.setColor(QPalette.Button, QtCore.Qt.white if self.windowFlags() & QtCore.Qt.WindowStaysOnTopHint else QColor(53, 53, 53))
        self._stay_on_top.setPalette(pal)
        self.show()

    def init_window(self):
        self._label = QLabel('')
        self._label.setMouseTracking(True)
        self._label.setSizePolicy(QSizePolicy.Preferred, QSizePolicy.Fixed)
        self._minimize = self.colored_button(QtCore.Qt.yellow)
        self._minimize.clicked.connect(self.showMinimized)
        if self._is_main_window:
            self._maximize = self.colored_button(QtCore.Qt.green)
            self._maximize.clicked.connect(self.toggle_maximized)
        else:
            self._stay_on_top = self.colored_button(QtCore.Qt.white if (sys.platform == 'linux' or sys.platform == 'linux2') \
                                                    and not self._is_main_window else QColor(53, 53, 53))
            self._stay_on_top.clicked.connect(self.set_on_top)
        self._close = self.colored_button(QtCore.Qt.red)
        self._close.clicked.connect(self.close)
        self._title_layout = QHBoxLayout()
        if self._is_main_window:
            widgets = [self._label, self._minimize, self._maximize, self._close]
        else:
            widgets = [self._label, self._stay_on_top, self._minimize, self._close]
        for widget in widgets:
            self._title_layout.addWidget(widget)
        line = QFrame()
        line.setFixedHeight(1)
        line.setFrameStyle(QFrame.HLine | QFrame.Plain)
        line.setStyleSheet("border: 1px solid rgb(70,70,70);")
        self._layout = QVBoxLayout()
        self._layout.addLayout(self._title_layout)
        self._layout.addWidget(line)
        self._layout.addStretch(-1)
        self.setLayout(self._layout)

    def layout(self):
        for i in range(self._layout.count()):
            layoutItem = self._layout.itemAt(i)
            if layoutItem.spacerItem():
                self._layout.removeItem(layoutItem)
        return self._layout

    def setWindowTitle(self, title):
        super().setWindowTitle(title)
        self._label.setText(title)

    def mouseMoveEvent(self, event):
        super().mouseMoveEvent(event)
        if not self._handle or self._pos_before_drag is None:
            self._pos_before_drag = event.globalPos()
        before = self.mouse_state()
        after = self.check_mouse(event)
        if before != after or after:
            event = self.check_event(event)
            self.set_drag_cursor()
            self.resize_window(event)
            self.move_window(event)
            self.check_snapping()

    def available_geometry(self):
        desktop = QDesktopWidget()
        return desktop.availableGeometry(desktop.screenNumber(self))

    def is_maximized(self):
        return self.available_geometry() == self.geometry()

    def set_maximized(self):
        desktop = QDesktopWidget()
        self._saved_geometry = self.saveGeometry()
        self.resize(self.available_geometry().size())
        self.move(0, 0)

    def restore(self):
        if self._saved_geometry is not None:
            self.restoreGeometry(self._saved_geometry)

    def toggle_maximized(self, event=None):
        if not self.is_maximized():
            self.set_maximized()
        else:
            self.restore()

    def check_snapping(self):
        if not self._is_main_window or not self._handle or self._right or self._bottom:
            return
        available_geometry = self.available_geometry()
        geometry = self.geometry()
        pos = QCursor.pos()
        top = pos.y() <= available_geometry.top() + 5
        left = pos.x() <= available_geometry.left() + 5
        right = pos.x() >= available_geometry.right() - 5
        bottom = pos.y() >= available_geometry.bottom() - 5
        if not (top or left or right):
            self._window.hide()
            return
        center  = available_geometry.bottomRight() / 2
        if top and left:
            top_left = available_geometry.topLeft()
            bottom_right = center
        elif top and right:
            top_left = available_geometry.topLeft() + QtCore.QPoint(center.x(), 0)
            bottom_right = center
        elif bottom and left:
            top_left = available_geometry.topLeft() + QtCore.QPoint(0, center.y())
            bottom_right = center
        elif bottom and right:
            top_left = center
            bottom_right = center
        elif top:
            top_left = available_geometry.topLeft()
            bottom_right = center * 2
        elif left:
            top_left = available_geometry.topLeft()
            bottom_right = center + QtCore.QPoint(0, center.y())
        elif right:
            top_left = available_geometry.topLeft() + QtCore.QPoint(center.x(), 0)
            bottom_right = center + QtCore.QPoint(0, center.y())
        if self._window.geometry().topLeft() == top_left and self._window.geometry().bottomRight() == bottom_right:
            return
        self._window.move(top_left)
        self._window.resize(bottom_right.x(), bottom_right.y())
        self._window.show()

    def mousePressEvent(self, event):
        super().mousePressEvent(event)
        if QtCore.Qt.MouseButton.LeftButton == event.button():
            self._handle = True
            self._drag = False
            self._pos_starting_drag = self.mapToGlobal(event.pos())
        if not self._drag_timer.isActive():
            self._drag_timer.start(self._double_click_interval)

    def mouseReleaseEvent(self, event):
        super().mouseReleaseEvent(event)
        if QtCore.Qt.MouseButton.LeftButton == event.button():
            self._handle = False
            self._drag = False
        if not self._click_timer.isActive():
            self._click_timer.start(self._double_click_interval)
        else:
            self._click_timer.stop()
            self.double_click()
        if self._drag_timer.isActive():
            self._drag_timer.stop()
        if self._window.isVisible():
            self.setGeometry(self._window.geometry())
            self._window.setVisible(False)

    def single_click(self):
        self._drag = False

    def double_click(self):
        if self._is_main_window and self.cursor() == QtCore.Qt.DragCopyCursor:
            self.toggle_maximized()

    def drag(self):
        self._drag = True

    def check_event(self, event):
        desktop = QDesktopWidget()
        pos = event.globalPos()
        available_geometry = self.available_geometry()
        if pos.x() < available_geometry.left():
            pos.setX(available_geometry.left())
        if pos.x() > available_geometry.right():
            pos.setX(available_geometry.right())
        if pos.y() < available_geometry.top():
            pos.setY(available_geometry.top())
        if pos.y() > available_geometry.bottom():
            pos.setY(available_geometry.bottom())
        QCursor.setPos(pos)
        event.setLocalPos(self.mapFromGlobal(pos))
        return event

    def move_window(self, event):
        if self._handle and self.cursor() == QtCore.Qt.DragCopyCursor:
            delta = event.globalPos() - self._pos_before_drag
            self.move(self.x() + delta.x(), self.y() + delta.y())
            self._pos_before_drag = event.globalPos()

    def resize_window(self, event):
        if not self._is_main_window and not self._height_resizable:
            return
        if (self._handle and self._right and self._is_main_window) or (self._handle and self._bottom and self._height_resizable):
            rect = self.geometry()
            x1 = rect.x()
            y1 = rect.y()
            x2 = event.pos().x() if self._right and self._handle else rect.width()
            y2 = event.pos().y() if self._bottom and self._handle else rect.height()
            rect.setRect(x1, y1, x2, y2)
            self.setGeometry(rect)

    def mouse_state(self):
        return self._top or self._left or self._bottom or self._right or self._handle

    def check_mouse(self, event):
        pos = event.pos()
        if not self._handle:
            self._bottom = pos.y() > self.geometry().height() - 10
            self._right = pos.x() > self.geometry().width() - 10
            self._top = False if self._right else pos.y() < 12 or self._label.underMouse()
            self._left = False if self._bottom else pos.x() < 10
        return self.mouse_state()

    def set_drag_cursor(self):
        if not self._is_main_window and (self._left or self._top or self._right or (self._bottom and not self._height_resizable)):
            self.setCursor(QtCore.Qt.DragCopyCursor)
        elif not self._is_main_window and self._height_resizable and self._bottom:
            self.setCursor(QtCore.Qt.SizeVerCursor)
        elif self._left or self._top:
            self.setCursor(QtCore.Qt.DragCopyCursor)
        elif self._bottom and self._right:
            self.setCursor(QtCore.Qt.SizeFDiagCursor)
        elif self._bottom:
            self.setCursor(QtCore.Qt.SizeVerCursor)
        elif self._right:
            self.setCursor(QtCore.Qt.SizeHorCursor)
        else:
            self.setCursor(QtCore.Qt.ArrowCursor)
```
