## Supplementary material for "High-resolution animal tracking with integration of environmental information in aquatic systems": Code_track_utility: Reader.py

```
from PySide2.QtGui import QPixmap, QImage
from PySide2.QtCore import QThread, Signal, QObject, QMutex

import numpy as np
import cv2
import psutil
import time

mutex = QMutex()

class BufferSignal(QObject):
    buffered = Signal(int)

class Buffer(QThread):

    def __init__(self, file_name, height, width):
        super().__init__()
        self.signal = BufferSignal()
        self.cap = cv2.VideoCapture(file_name)
        self.frame_count = int(self.cap.get(cv2.CAP_PROP_FRAME_COUNT))
        self.height = height
        self.width = width
        self.running = False
        self.init_buffer()

    def init_buffer(self):
        self.buffer_size = min(500, np.floor(0.5 * psutil.virtual_memory().available / (self.height * self.width * 3)).astype(np.int))
        self.frame = 0
        self.frames = np.arange(self.buffer_size)
        self.buffered = np.zeros(self.buffer_size, dtype=np.bool)
        self.imgs = np.zeros((self.buffer_size, self.height, self.width, 3), dtype=np.uint8)

    def buffered_frame(self):
        buffered_frames = self.frames[self.buffered]
        if buffered_frames.size > 0:
            return buffered_frames[-1]
        return 0

    def buffered_percent(self):
        return int(100 * self.buffered.sum() / self.buffered[self.frames < self.frame_count].size)

    def run(self):
        self.running = True
        self.buffered = np.zeros(self.buffer_size, dtype=np.bool)
        self.frames = np.arange(self.buffer_size) + self.frame
        if self.frame != self.cap.get(cv2.CAP_PROP_POS_FRAMES):
            self.cap.set(cv2.CAP_PROP_POS_FRAMES, self.frame)
        while self.frame <= self.frames[-1] and self.running and self.frame < self.frame_count:
            ret, img = self.cap.read()
            if ret:
                self.imgs[self.frames == self.frame] = img
                self.buffered[self.frames == self.frame] = True
            self.frame += 1
            self.signal.buffered.emit(self.buffered_percent())
        self.running = False

class ReaderSignal(QObject):
    image = Signal(QPixmap)
    frame = Signal(int)
    toggle = Signal()

class Reader(QThread):

    def __init__(self, file_name):
        super().__init__()
        self._video = False
        self.signal = ReaderSignal()
        self.running = False
        self.default = None
        self.frame_count = 1
        self.fps = 30
        self.frame = 0
        self.height = 1520
        self.width = 2704
        self.cap = cv2.VideoCapture(file_name) if file_name is not None else None
        if self.cap is not None and self.cap.isOpened():
            self.init_video(file_name)
        self.default = self.prepare_img(np.zeros((self.height, self.width, 3), dtype=np.uint8))

    @property
    def video(self):
        return self._video

    @video.setter
    def video(self, value):
        if self.cap is not None:
            self._video = value
        if not self.running:
            self.read()

    def init_video(self, file_name):
        self.width = int(self.cap.get(cv2.CAP_PROP_FRAME_WIDTH))
        self.height = int(self.cap.get(cv2.CAP_PROP_FRAME_HEIGHT))
        self.frame_count = int(self.cap.get(cv2.CAP_PROP_FRAME_COUNT))
        self.fps = int(self.cap.get(cv2.CAP_PROP_FPS))
        self.buffer = Buffer(file_name, self.height, self.width)
        self.video = True

    def prepare_img(self, img):
        img = cv2.cvtColor(img, cv2.COLOR_BGR2RGB).copy()
        return QPixmap(QImage(img.data, img.shape[1], img.shape[0], img.strides[0], QImage.Format_RGB888))

    def start_buffer(self):
        if not self.video:
            return
        mutex.lock()
        self.buffer.running = False
        self.buffer.wait()
        self.buffer.frame = self.frame
        self.buffer.start()
        mutex.unlock()

    def read(self, from_video=False):
        start = time.time()
        if self.video:
            mutex.lock()
            if np.isin(self.frame, self.buffer.frames) and self.buffer.buffered[self.buffer.frames == self.frame] and not from_video:
                ret, img = True, self.buffer.imgs[self.buffer.frames == self.frame].reshape(self.height, self.width, 3)
                mutex.unlock()
            else:
                mutex.unlock()
                if self.frame != self.cap.get(cv2.CAP_PROP_POS_FRAMES):
                    self.cap.set(cv2.CAP_PROP_POS_FRAMES, self.frame)
                ret, img = self.cap.read()
            if ret:
                image = self.prepare_img(img)
                self.signal.image.emit(image)
                self.signal.frame.emit(self.frame)
                if from_video:
                    return img[:, :, ::-1]
            else:
                self.signal.image.emit(self.default)
                self.signal.frame.emit(self.frame)
        else:
            self.signal.image.emit(self.default)
            self.signal.frame.emit(self.frame)
        stop = time.time()
        msec = (stop - start) * 1000
        return msec

    def run(self):
        self.running = True
        self.signal.toggle.emit()
        while self.running and self.frame < self.frame_count - 1:
            self.frame += 1
            msec = self.read()
            msec_wait = 1000 / self.fps - msec
            if msec_wait > 0:
                self.msleep(msec_wait)
        self.running = False
        self.signal.toggle.emit()
```
