## Supplementary material for "High-resolution animal tracking with integration of environmental information in aquatic systems": Code_track_utility: Settings.py

```
from PySide2.QtWidgets import QWidget, QVBoxLayout, QHBoxLayout, QCheckBox, QSpinBox, QLabel, QButtonGroup
from PySide2 import QtCore

from .QtUtils import named_cb, NamedSpinBox, BorderlessWindow

class Settings(BorderlessWindow):

    def __init__(self, options):
        super().__init__(main_window=False)

        self.setWindowTitle('Settings')

        self.cb_pose = named_cb('Draw poses', options.pose)
        self.cb_position = named_cb('Draw positions', options.position)
        self.cb_annotation = named_cb('Draw annotations', options.annotation)
        self.cb_interpolate = named_cb('Interpolate tracks', options.interpolated)
        self.cb_connect = named_cb('Synchronize connected in set frame', options.connected)
        self.cb_fix = named_cb('Fix identities in merge connected', options.fix)

        self.sb_position = QSpinBox()
        self.sb_n_inds = QSpinBox()
        self.sb_timeline_snapping = QSpinBox()

        self.sb_position.setEnabled(self.cb_position.checkState())
        self.cb_position.stateChanged.connect(self.sb_position.setEnabled)

        option_group = QButtonGroup(self)
        layout_option_group = QHBoxLayout()
        for cb in [self.cb_pose, self.cb_position, self.cb_annotation]:
            option_group.addButton(cb)
            layout_option_group.addWidget(cb)

        layout = self.layout()
        layout.addLayout(layout_option_group)
        layout.addWidget(NamedSpinBox(self.sb_position, 'Position radius (px)', options.position_size, 1, 100))
        layout.addWidget(NamedSpinBox(self.sb_n_inds, 'Number of individuals', options.n_inds, 1, 100))
        layout.addWidget(NamedSpinBox(self.sb_timeline_snapping, 'Timeline cursor snapping (px)', options.timeline_snapping, 0, 50))
        layout.addWidget(self.cb_interpolate)
        layout.addWidget(self.cb_connect)
        layout.addWidget(self.cb_fix)

        self.setLayout(layout)
```
