## Supplementary material for "High-resolution animal tracking with integration of environmental information in aquatic systems": Code_track_utility: Timeline.py

```
from PySide2.QtWidgets import QGraphicsItem, QGraphicsRectItem, QApplication, QGraphicsLineItem
from PySide2 import QtCore, QtGui, QtWidgets
from PySide2.QtCore import Signal, QObject, QRectF

import numpy as np

import time

class TimelineSignal(QObject):
    identity = Signal(int)
    hovered = Signal(int)
    diff = Signal(int)

class Timeseries(QGraphicsItem):

    def __init__(self, series, identity, color, selected, hovered, row, scale):
        super().__init__()
        self.setToolTip(str(identity))
        self.series = series
        self.color = color
        self.row = row
        self.scale = scale
        self.selected = selected
        self.hovered = hovered
        self.width = 2 if self.hovered and not self.selected or self.hovered else 1
        self.pen = QtGui.QPen(QtGui.QColor(255, 255, 255, 255), self.width)
        self.brush = QtGui.QBrush(QtGui.QColor(*self.color, 255))
        self.parts = []
        if self.selected:
            part = QGraphicsRectItem(QRectF(self.scale * self.series[0].min() - 2.5,
                                            self.row * 15 - 2.5,
                                            self.scale * (self.series[-1].max() - self.series[0].min()) + 5,
                                            7.5 + 5))
            part.setBrush(QtGui.QBrush(QtGui.QColor(*self.color, 100)))
            part.setPen(QtGui.QPen(QtGui.QColor(*self.color, 255), 1))
            self.parts.append(part)
        for part in series:
            part = QGraphicsRectItem(QRectF(self.scale * part.min(),
                                            self.row * 15,
                                            self.scale * (part.max() - part.min()),
                                            7.5))
            part.setBrush(self.brush)
            part.setPen(self.pen)
            self.parts.append(part)

    def boundingRect(self):
        if self.selected:
            rect = QRectF(self.scale * self.series[0].min() - 2.5 - 0.5,
                          self.row * 15 - 2.5 - 0.5,
                          self.scale * (self.series[-1].max() - self.series[0].min()) + 5 + 1,
                          7.5 + 5 + 1)
        else:
            rect = QRectF(self.scale * self.series[0].min() - self.width / 2,
                          self.row * 15 - self.width / 2,
                          self.scale * (self.series[-1].max() - self.series[0].min()) + self.width + 1,
                          7.5 + self.width)
        return rect

    def paint(self, painter, option, widget):
        for part in self.parts:
            part.paint(painter, option, widget)

class Timeline(QtWidgets.QGraphicsView):

    def __init__(self, menu_trajectory, menu_annotation, options):
        super().__init__()
        self.setFrameStyle(0)
        self.signal = TimelineSignal()
        self._scene = QtWidgets.QGraphicsScene(self)
        self.setScene(self._scene)
        self.setAlignment((QtCore.Qt.AlignLeft | QtCore.Qt.AlignTop))
        self.setHorizontalScrollBarPolicy(QtCore.Qt.ScrollBarAlwaysOff)
        self.setBackgroundBrush(QtGui.QBrush(QtGui.QColor(15, 15, 15)))
        self.setFrameShape(QtWidgets.QFrame.NoFrame)
        self.setMouseTracking(True)
        self.annotations = ({}, None)
        self.timeseries = []
        self.timepoints = []
        self.identities = []
        self.colors = []
        self.selected = []
        self.hovered = []
        self.timeline_snapping = options.timeline_snapping
        self.time_window = 600
        self.position = 0
        self.frame = 0
        self.fps = 30
        self.menus_toggle = True
        self.menu_trajectory = menu_trajectory
        self.menu_annotation = menu_annotation
        self.width = self.get_width()
        self.height = self.get_height()
        self.scale = self.width / self.time_window
        self.position_line = None
        self.draw_time()

    def draw_position(self, event):
        pos = self.mapFromGlobal(QtGui.QCursor.pos())
        modifiers = QApplication.keyboardModifiers()
        if event.type() == QtCore.QEvent.Leave or \
           (event.type() == QtCore.QEvent.KeyRelease and event.key() == QtCore.Qt.Key_Control) or \
           (event.type() == QtCore.QEvent.KeyRelease and modifiers != QtCore.Qt.ControlModifier) or \
           (event.type() == QtCore.QEvent.KeyPress and event.key() != QtCore.Qt.Key_Control and modifiers != QtCore.Qt.ControlModifier) or \
           (event.type() == QtCore.QEvent.MouseMove and modifiers != QtCore.Qt.ControlModifier):
            if self.position_line is not None:
                self._scene.removeItem(self.position_line)
                self.position_line = None
            return
        x = pos.x()
        if len(self.timepoints) > 0 and self.timeline_snapping > 0:
            distances = np.absolute(x - self.timepoints)
            if np.min(distances) < self.timeline_snapping:
                x = self.timepoints[np.argmin(distances)]
        pen = QtGui.QPen(QtGui.QColor(255, 255, 255, 200), 2)
        pen.setStyle(QtCore.Qt.DashLine)
        if self.position_line is not None:
            self._scene.removeItem(self.position_line)
        self.position_line = self._scene.addLine(QtCore.QLineF(x, 0, x, self.height), pen=pen)
        self._scene.update()

    def draw_annotation_histogram(self):
        if self.annotations[1] is None:
            return
        self.scale = self.width / self.time_window
        timepoints = np.arange(0, self.time_window + 1)
        offset = self.frame - self.annotations[1]
        max_count = 100
        pen = QtGui.QPen(QtGui.QColor(255, 255, 255, 200), self.scale - 1)
        pen.setCapStyle(QtCore.Qt.FlatCap)
        for t in timepoints:
            frame = t - offset
            if not frame in self.annotations[0]:
                continue
            n_annotations = sum([len(self.annotations[0][frame][occlusion_level]['polygons']) for occlusion_level in self.annotations[0][frame]])
            self._scene.addLine(QtCore.QLineF(self.scale * t, self.height * (1 - n_annotations / max_count),
                                              self.scale * t, self.height),
                                pen=pen)

    def draw_timeseries(self):
        if self.width == 0:
            return
        scale = self.width / self.time_window
        if len(self.timeseries) > 0:
            self.timepoints = np.concatenate([np.array([(part.min(), part.max()) for part in series]).ravel() \
                                              for series in self.timeseries]) * scale
        for idx in range(len(self.timeseries)):
            self._scene.addItem(Timeseries(self.timeseries[idx],
                                           self.identities[idx],
                                           self.colors[idx],
                                           self.selected[idx],
                                           self.hovered[idx],
                                           idx + 1, scale))

    def draw_time(self):
        self.scale = self.width / self.time_window
        timepoints = np.append(np.arange(0, self.time_window + 1, self.fps), self.frame)
        for t in timepoints:
            current = t == self.frame
            pen = QtGui.QPen(QtGui.QColor(255, 255, 255, 200 if current else 100), 2 if current else 1)
            pen.setCapStyle(QtCore.Qt.FlatCap)
            if current:
                self.position = self.scale * t
            self._scene.addLine(QtCore.QLineF(self.scale * t, 0, self.scale * t, self.height),
                                pen=pen)

    def clear(self):
        for item in self._scene.items():
            if item != self.position_line:
                self._scene.removeItem(item)

    def fitInView(self):
       self.height = self.get_height()
       self.clear()
       self.draw_timeseries()
       self.draw_time()
       self.draw_annotation_histogram()
       self.adjust_scene()

    def resizeEvent(self, event):
        self.width = self.get_width()
        self.height = self.get_height()
        self.fitInView()
        super().resizeEvent(event)

    def get_width(self):
        return self.mapToScene(self.viewport().geometry()).boundingRect().width()

    def get_height(self):
        return self.mapToScene(self.viewport().geometry()).boundingRect().height() # max(15 * (len(self.timeseries)), self.mapToScene(self.viewport().geometry()).boundingRect().height() - 2)

    def adjust_scene(self):
        r = self._scene.itemsBoundingRect()
        view_rect = self.mapToScene(self.viewport().rect()).boundingRect()
        w = max(r.size().width(), view_rect.size().width())
        h = max(r.size().height(), view_rect.size().height()) # - 1
        self._scene.setSceneRect(QtCore.QRectF(QtCore.QPointF(), QtCore.QSizeF(w, h)))

    def mouseMoveEvent(self, event):
        self.draw_position(event)
        modifiers = QApplication.keyboardModifiers()
        item = self._scene.itemAt(self.mapToScene(event.pos()), QtGui.QTransform())
        if item is not None and item.toolTip() != '' and modifiers != QtCore.Qt.ControlModifier:
            self.signal.hovered.emit(int(item.toolTip()))
        else:
            self.signal.hovered.emit(-1)

    def mousePressEvent(self, event):
        modifiers = QApplication.keyboardModifiers()
        if event.button() == QtCore.Qt.MouseButton.LeftButton and modifiers != QtCore.Qt.ControlModifier:
            self.viewport().setCursor(QtCore.Qt.ClosedHandCursor)
            item = self._scene.itemAt(self.mapToScene(event.pos()), QtGui.QTransform())
            if item is not None and item.toolTip() != '':
                self.signal.identity.emit(int(item.toolTip()))
        elif event.button() == QtCore.Qt.MouseButton.RightButton and modifiers != QtCore.Qt.ControlModifier:
            if self.menus_toggle:
                self.menu_trajectory.popup(event.globalPos())
            else:
                self.menu_annotation.popup(event.globalPos())
        elif modifiers == QtCore.Qt.ControlModifier:
            diff = (self.position_line.line().p1().x() - self.position) / self.scale
            self.signal.diff.emit(int(np.round(diff)))

    def mouseReleaseEvent(self, event):
        self.viewport().setCursor(QtCore.Qt.ArrowCursor)

    def keyPressEvent(self, event):
        super().keyPressEvent(event)
        self.draw_position(event)

    def keyReleaseEvent(self, event):
        super().keyReleaseEvent(event)
        self.draw_position(event)

    def enterEvent(self, event):
        super().enterEvent(event)
        self.setFocus(QtCore.Qt.FocusReason.MouseFocusReason)

    def leaveEvent(self, event):
        super().leaveEvent(event)
        self.clearFocus()
        self.draw_position(event)
```
