## Supplementary material for "High-resolution animal tracking with integration of environmental information in aquatic systems": Code_track_utility: Tracks.py

```
import numpy as np

from copy import deepcopy
from multiprocessing import Pool, cpu_count

import time

def filter_trajectory(trajectory, condition):
    trajectory_filtered = {}
    for key in trajectory:
        trajectory_filtered[key] = trajectory[key][condition]
    return trajectory_filtered

def split_trajectory(trajectory, frame):
    trajectory_pre = filter_trajectory(trajectory, frame > trajectory['FRAME_IDX'])
    trajectory_post = filter_trajectory(trajectory, frame <= trajectory['FRAME_IDX'])
    return trajectory_pre, trajectory_post

def append_trajectory(trajectory_pre, trajectory_post):
    if np.isin(trajectory_pre['FRAME_IDX'], trajectory_post['FRAME_IDX']).any():
        return trajectory_pre
    appended = {}
    sorted_idx = np.argsort(np.append(trajectory_pre['FRAME_IDX'], trajectory_post['FRAME_IDX']))
    for key in trajectory_pre:
        appended[key] = np.append(trajectory_pre[key], trajectory_post[key], axis=0)[sorted_idx]
    return appended

def trajectory_lookup(i):
    return np.isin(frames, tracks[str(i)]['FRAME_IDX'])

def init_lookup(f, t):
    global frames, tracks
    frames = f
    tracks = t

def identity_lookup(tracks):
    frames = np.array([])
    if tracks['FRAME_IDX'].size > 0:
        frames = np.arange(0, tracks['FRAME_IDX'].max() + 1)
    # with Pool(processes=cpu_count(), initializer=init_lookup, initargs=(frames, tracks, )) as pool:
    #     lookup = pool.map(trajectory_lookup, tracks['IDENTITIES'])
    #     pool.close()
    #     pool.join()
    # lookup = np.transpose(lookup)
    lookup = np.transpose([np.isin(frames, tracks[str(i)]['FRAME_IDX']) \
                           for i in tracks['IDENTITIES']])
    return lookup

def update_frames(tracks):
    tracks['FRAME_IDX'] = np.unique(np.concatenate([tracks[str(i)]['FRAME_IDX'] for i in tracks['IDENTITIES']])).astype(np.int)
    return tracks

def pop_trajectory(tracks, i, lookup):
    if lookup is not None:
        lookup = lookup[:, tracks['IDENTITIES'] != i]
    tracks['IDENTITIES'] = tracks['IDENTITIES'][tracks['IDENTITIES'] != i]
    return tracks.pop(str(i)), lookup

def delete_trajectories(tracks, identities, lookup):
    for i in identities:
        _, lookup = pop_trajectory(tracks, i, lookup)
    tracks = update_frames(tracks)
    return tracks, lookup

def add_trajectories(tracks, trajectories, lookup):
    new_identities = np.arange(tracks['IDENTITIES'].max() + 1, tracks['IDENTITIES'].max() + 1 + len(trajectories))
    tracks['IDENTITIES'] = np.append(tracks['IDENTITIES'], new_identities)
    if lookup is not None:
        lookup = np.append(lookup, np.zeros((lookup.shape[0], new_identities.size), dtype=np.bool), axis=1)
        frames = np.arange(lookup.shape[0])
    for i, trajectory in zip(new_identities, trajectories):
        tracks[str(i)] = trajectory
        if lookup is not None:
            lookup[:, tracks['IDENTITIES'] == i] = np.isin(frames, tracks[str(i)]['FRAME_IDX']).reshape(-1, 1)
    tracks = update_frames(tracks)
    return tracks, lookup

def split_trajectories(tracks, identities, frame, lookup):
    trajectories_pre = []
    trajectories_post = []
    for i in identities:
        trajectory_pre, trajectory_post = split_trajectory(tracks[str(i)], frame)
        trajectories_pre.append(trajectory_pre)
        trajectories_post.append(trajectory_post)
    for i, trajectory_pre in zip(identities, trajectories_pre):
        if trajectory_pre['FRAME_IDX'].size > 0:
            tracks[str(i)] = trajectory_pre
            if lookup is not None:
                lookup[frame:, tracks['IDENTITIES'] == i] = False
        else:
            tracks, lookup = delete_trajectories(tracks, [i], lookup)
    tracks, lookup = add_trajectories(tracks, [trajectory for trajectory in trajectories_post if trajectory['FRAME_IDX'].size > 0], lookup)
    return tracks, lookup

def switch_trajectories(tracks, identities, frame, lookup):
    if len(identities) != 2:
        return tracks, lookup
    if lookup is not None:
        frames = np.arange(lookup.shape[0])
    trajectories_pre = []
    trajectories_post = []
    for i in identities:
        trajectory_pre, trajectory_post = split_trajectory(tracks[str(i)], frame)
        trajectories_pre.append(trajectory_pre)
        trajectories_post.append(trajectory_post)
    for i, trajectory_pre, trajectory_post in zip(identities, trajectories_pre, trajectories_post[::-1]):
        tracks[str(i)] = append_trajectory(trajectory_pre, trajectory_post)
        if lookup is not None:
            lookup[:, tracks['IDENTITIES'] == i] = np.isin(frames, tracks[str(i)]['FRAME_IDX']).reshape(-1, 1)
    return tracks, lookup

def merge_trajectories(tracks, identities, lookup):
    identity = identities[0]
    merge_identities = identities[1:]
    for i in merge_identities:
        if np.isin(tracks[str(identity)]['FRAME_IDX'], tracks[str(i)]['FRAME_IDX']).any():
            return tracks, lookup
    for i in merge_identities:
        trajectory, lookup = pop_trajectory(tracks, i, lookup)
        tracks[str(identity)] = append_trajectory(tracks[str(identity)], trajectory)
    lookup[:, tracks['IDENTITIES'] == identity] = np.isin(np.arange(lookup.shape[0]), tracks[str(identity)]['FRAME_IDX']).reshape(-1, 1)
    return tracks, lookup

def delete_short_trajectories(tracks, min_length):
    short = identity_lookup(tracks).sum(axis=0) < min_length
    tracks, _ = delete_trajectories(tracks, tracks['IDENTITIES'][short], None)
    return tracks

def interpolate_tracks(tracks, smooth_window=0, n_iter=3, degree=1):
    tracks_copy = deepcopy(tracks)
    tracks['FRAME_IDX'] = np.array([])
    for i in tracks['IDENTITIES']:
        frame_idx = np.arange(tracks[str(i)]['FRAME_IDX'][0], tracks[str(i)]['FRAME_IDX'][-1] + 1)
        for key in list(tracks[str(i)]):
            if key in ['X', 'Y', 'Z']:
                tracks[str(i)][key] = np.interp(frame_idx, tracks[str(i)]['FRAME_IDX'], tracks[str(i)][key])
            elif key == 'SPINE':
                spine = np.zeros((frame_idx.size, *tracks[str(i)][key].shape[1:]))
                for u in range(spine.shape[1]):
                    for v in range(spine.shape[2]):
                        spine[:, u, v] = np.interp(frame_idx, tracks[str(i)]['FRAME_IDX'], tracks[str(i)][key][:, u, v])
                tracks[str(i)][key] = spine
            elif key == 'RADII':
                radii = np.zeros((frame_idx.size, tracks[str(i)][key].shape[1]))
                for u in range(radii.shape[1]):
                    radii[:, u] = np.interp(frame_idx, tracks[str(i)]['FRAME_IDX'], tracks[str(i)][key][:, u])
                tracks[str(i)][key] = radii
            elif key != 'FRAME_IDX':
                tracks[str(i)].pop(key)
        tracks[str(i)]['FRAME_IDX'] = frame_idx
        tracks['FRAME_IDX'] = np.append(tracks['FRAME_IDX'], tracks[str(i)]['FRAME_IDX'])
    tracks['FRAME_IDX'] = np.unique(tracks['FRAME_IDX']).astype(np.int)
    tracks['IDENTITIES'] = tracks['IDENTITIES'].astype(np.int)
    return tracks, tracks_copy
```
