## Supplementary material for "High-resolution animal tracking with integration of environmental information in aquatic systems": Code_track_utility: TrackUtil.py

```
import os

from PySide2 import QtCore
from PySide2.QtGui import QCursor
from PySide2.QtWidgets import QSplitter, QPushButton, QWidget, QVBoxLayout, QHBoxLayout, QSlider, QLabel, QFileDialog, QMenu, QToolTip, QStatusBar, QProgressBar, QSpacerItem, QRubberBand
from PySide2.QtCore import Qt

import numpy as np

from .UI import *
from .QtUtils import *
from .Utils import *
from .Tracks import *
from .Drawing import *

class Options:

    def __init__(self):
        self.n_inds = 1
        self.pose = True
        self.position = False
        self.annotation = False
        self.position_size = 25
        self.timeline_snapping = 5
        self.class_names = []
        self.connected = False
        self.fix = False
        self.interpolated = False

    def add_class(self, class_name):
        if class_name not in self.class_names:
            self.class_names.append(class_name)

    def remove_class(self, class_name):
        if class_name in self.class_names:
            self.class_names.remove(class_name)

class TrackUtil(BorderlessWindow):

    def __init__(self, parent=None):
        super().__init__()

        self.parent = parent
        self.children = []

        self.options = Options()
        self._extended = False
        self._playing = False
        self._hovered = -1
        self._selected = set()

        self.setWindowTitle('TrackUtil' + (' --- (connected)' if self.parent is not None else ''))

        self.button_play = QPushButton('Play')
        self.button_play.clicked.connect(lambda v: setattr(self, 'playing', v))

        self.button_extended = QPushButton('Extend')
        self.button_extended.clicked.connect(lambda v: setattr(self, 'extended', v))

        self.button_video = QPushButton('Video ON')
        self.button_video.clicked.connect(lambda v: setattr(self, 'video', v))

        self.button_previous = HoldButton('<<')
        self.button_previous.clicked.connect(self.previous_frame)

        self.button_next = HoldButton('>>')
        self.button_next.clicked.connect(self.next_frame)

        file_menu = QMenu(self)
        file_menu.addAction('Load video', lambda: load_video(self))
        file_menu.addAction('Load tracks', lambda: load_tracks(self))
        file_menu.addAction('Load annotations', lambda: load_annotations(self))
        file_menu.addSeparator()
        file_menu.addAction('Save tracks', lambda: save_tracks(self))
        file_menu.addAction('Save annotations', lambda: save_annotations(self))
        file_menu.addSeparator()
        file_menu.addAction('Save progress', lambda: make_backup(self))
        file_menu.addAction('Load progress', lambda: load_backup(self))
        file_menu.addSeparator()
        file_menu.addAction('Write to dataset', lambda: write_dataset(self))
        file_menu.addSeparator()
        file_menu.addAction('Connect', lambda: open_child(self))

        button_file = QPushButton('File')
        button_file.setMenu(file_menu)

        self.button_buffer = QPushButton('Buffer')
        self.button_settings = QPushButton('Settings')
        self.button_annotator = QPushButton('Annotator')
        self.button_history = QPushButton('History')

        self.slider_fine = JumpSlider(QtCore.Qt.Horizontal)
        self.slider_fine.setRange(0, 0)
        self.slider_fine.setTickPosition(QSlider.TickPosition.TicksBelow)

        self.slider = JumpSlider(QtCore.Qt.Horizontal)
        self.slider.setRange(0, 0)
        self.slider.setTickPosition(QSlider.TickPosition.TicksBelow)

        play_control = QHBoxLayout()
        play_control.setMargin(0)
        play_control.addWidget(self.button_previous)
        play_control.addWidget(self.button_play)
        play_control.addWidget(self.button_next)
        play_control.addWidget(self.slider)
        play_control.addWidget(self.button_extended)

        self.play_control = QWidget()
        self.play_control.setLayout(play_control)

        extended_control = QHBoxLayout()
        extended_control.setMargin(0)
        extended_control.addWidget(self.slider_fine)

        self.extended_control = QWidget()
        self.extended_control.setLayout(extended_control)
        self.extended_control.setVisible(self.extended)

        self.buffered = QProgressBar()
        self.buffered.setRange(0, 100)
        self.buffered.setTextVisible(False)
        self.buffered.setMaximumWidth(100)

        self.status_bar = QStatusBar()
        self.status_bar.setStyleSheet('QStatusBar::item {border: None;}')
        self.status_bar.addPermanentWidget(self.button_buffer)
        self.status_bar.addPermanentWidget(self.buffered)
        self.status_bar.addPermanentWidget(self.button_video)
        self.status_bar.addPermanentWidget(self.button_history)
        self.status_bar.addPermanentWidget(self.button_settings)
        self.status_bar.addPermanentWidget(self.button_annotator)

        init_menus(self)

        init_player(self)
        init_timeline(self)

        splitter = QSplitter()
        splitter.setOrientation(Qt.Vertical)
        splitter.addWidget(self.player)
        splitter.addWidget(self.timeline)

        button_tracks = QPushButton('Tracks')
        button_tracks.setMenu(self.player.menu_trajectory)

        self.status_bar.addPermanentWidget(button_tracks)
        self.status_bar.addPermanentWidget(button_file)
        self.status_bar.setSizeGripEnabled(False)

        layout = self.layout()
        layout.addWidget(self.status_bar)
        layout.addWidget(splitter)
        layout.addWidget(self.extended_control)
        layout.addWidget(self.play_control)

        self.setLayout(layout)

        init_history(self)
        init_video(self, select=False)
        init_tracks(self, select=False)
        init_annotations(self, select=False)
        init_settings(self)
        init_annotator(self)

        splitter.setSizes([np.sum(splitter.sizes()), 0])

        self.video=False
        init_shortcuts(self)

        self.resize(1280, 720)
        self.show()


    @property
    def extended(self):
        return self._extended

    @property
    def playing(self):
        return self._playing

    @property
    def video(self):
        return self.reader.video

    @property
    def hovered(self):
        return self._hovered

    @property
    def selected(self):
        return self._selected

    @extended.setter
    def extended(self, value):
        self._extended = not self._extended
        self.button_extended.setText('Hide' if self.extended else 'Extend')
        self.extended_control.setVisible(self.extended)

    @playing.setter
    def playing(self, value=None):
        self.reader.running = not self.reader.running
        if self.reader.running:
            self.reader.start()

    @video.setter
    def video(self, value):
        self.reader.video = not self.reader.video
        self.update_buttons()

    @hovered.setter
    def hovered(self, identity):
        if self._hovered != -1 and identity == -1:
            self._hovered = identity
            self.update_image()
            self.update_timeline()
        elif self._hovered != identity:
            self._hovered = identity
            self.update_image()
            self.update_timeline()
        else:
            self._hovered = identity

    @selected.setter
    def selected(self, identity):
        if identity in self._selected:
            self._selected.remove(identity)
        else:
            self._selected.add(identity)
        self.update_image()
        self.update_timeline()

    def update_buttons(self):
        self.button_video.setText('Video OFF' if self.reader.video else 'Video ON')
        self.button_play.setText('Pause' if self.reader.running else 'Play')

    def previous_frame(self):
        self.set_frame(self.reader.frame - 1)

    def next_frame(self):
        self.set_frame(self.reader.frame + 1)

    def set_frame(self, frame, from_video=False, connect=True):
        if frame < 0 or frame >= self.reader.frame_count:
            return
        if self.parent is None or from_video or not connect or not self.options.connected:
            if frame >= self.reader.frame_count or self.reader.running:
                self.playing = not self.playing
            self.reader.frame = frame
            img = self.reader.read(from_video)
            self.update_ui(frame)
            if from_video:
                return img
        if self.parent is not None and self.options.connected and connect:
            self.parent.set_frame(frame)
        elif self.parent is None and self.options.connected:
            for child in self.children:
                child.set_frame(frame, connect=False)

    def update_ui(self, frame):
        if not in_range(frame, self.slider_fine.minimum(), self.slider_fine.maximum()):
            self.slider_fine.blockSignals(True)
            self.slider_fine.setRange(*get_range(frame, self.reader.fps * 60, self.reader.frame_count - 1))
            self.slider_fine.blockSignals(False)
            slider_set_value(self.slider, frame)
        if frame != self.slider.value() and frame != self.slider_fine.value():
            slider_set_value(self.slider, frame)
            slider_set_value(self.slider_fine, frame)
        elif frame != self.slider.value() and not self.reader.running:
            slider_set_value(self.slider, frame)
        elif frame != self.slider_fine.value() and not self.reader.running:
            slider_set_value(self.slider_fine, frame)
        elif self.reader.running and not (frame == self.slider.value() and frame == self.slider_fine.value()):
            self.playing = not self.playing
        self.status_bar.showMessage('{} / {}'.format(frame, self.reader.frame_count - 1), 500)
        self.update_image()
        self.update_timeline()

    def closeEvent(self, event):
        if self.parent is not None:
            self.parent.children.remove(self)
        if len(self.children) > 0:
            self.status_bar.showMessage('Close all connected windows first.', 2000)
            event.ignore()
            return
        self.reader.running = False
        self.reader.wait()
        if hasattr(self.reader, 'buffer'):
            self.reader.buffer.running = False
            self.reader.buffer.wait()
        self.settings.close()
        self.annotator.close()
        self.history.close()
        super().closeEvent(event)

    def add_annotations(self, annotations, file_name=None, frame=None):
        if file_name is None and frame is None:
            connected = ([self.parent] if self.parent is not None else []) + self.children
            for c in connected:
                c.add_annotations(annotations, self.file_name, self.reader.frame)
                if not c.playing and c.reader.frame == self.reader.frame and c.file_name == self.file_name:
                    c.update_image()
                    c.update_timeline()
        file_name = self.file_name if file_name is None else file_name
        frame = self.reader.frame if frame is None else frame
        if file_name not in self.annotations:
            self.annotations[file_name] = {}
        self.annotations[file_name][frame] = annotations
        self.update_timeline()

    def get_poses(self, frame):
        poses = []
        color = []
        selected = []
        hovered = []
        if self.lookup.size == 0 or frame >= self.lookup.shape[0]:
            return poses, color, selected, hovered
        visible = self.lookup[frame, :]
        identities = self.tracks['IDENTITIES'][visible]
        colors = self.colors[visible]
        for i, c in zip(identities, colors):
            idx = np.argwhere(self.tracks[str(i)]['FRAME_IDX'] == frame).ravel()
            for x in idx:
                poses.append([self.tracks[str(i)]['SPINE'][x, :, ::-1],
                              self.tracks[str(i)]['RADII'][x],
                              i])
                color.append(c)
                selected.append(True if i in self.selected else False)
                hovered.append(True if i == self.hovered else False)
        return poses, color, selected, hovered

    def get_positions(self, frame):
        positions = []
        color = []
        selected = []
        hovered = []
        if self.lookup.size == 0 or frame >= self.lookup.shape[0]:
            return positions, color, selected, hovered
        visible = self.lookup[frame, :]
        identities = self.tracks['IDENTITIES'][visible]
        colors = self.colors[visible]
        for i, c in zip(identities, colors):
            idx = np.argwhere(self.tracks[str(i)]['FRAME_IDX'] == frame).ravel()
            for x in idx:
                positions.append([self.tracks[str(i)]['X'][x],
                                  self.tracks[str(i)]['Y'][x],
                                  self.options.position_size,
                                  i])
                color.append(c)
                selected.append(True if i in self.selected else False)
                hovered.append(True if i == self.hovered else False)
        return positions, color, selected, hovered

    def get_timeseries(self, frame):
        timeseries = []
        identities = []
        color = []
        selected = []
        hovered = []
        left = frame - self.reader.fps * 10
        right = frame + self.reader.fps * 10
        set_time = 0
        if left < 0:
            left = 0
            set_time = -1
        if right > self.reader.frame_count:
            right = self.reader.frame_count
            set_time = 1
        self.timeline.time_window = 1 + (right - left)
        if set_time == 0:
            self.timeline.frame = self.timeline.time_window // 2
        elif set_time < 0:
            self.timeline.frame = frame
        else:
            self.timeline.frame = frame - left
        if self.lookup.size == 0 or frame >= self.lookup.shape[0]:
            return timeseries, identities, color, selected, hovered
        visible = self.lookup[left:(right + 1), :].any(axis=0)
        identities = self.tracks['IDENTITIES'][visible].tolist()
        colors = self.colors[visible]
        for i, c in zip(identities, colors):
            frame_idx = self.tracks[str(i)]['FRAME_IDX']
            visible = self.tracks[str(i)]['FRAME_IDX'][(frame_idx >= left) & (frame_idx <= right)] - left
            series = np.split(visible, np.argwhere(np.diff(visible) > 1).ravel() + 1)
            timeseries.append(series)
            color.append(c)
            selected.append(True if i in self.selected else False)
            hovered.append(True if i == self.hovered else False)
        return timeseries, identities, color, selected, hovered

    def draw_poses(self):
        for pose, c, s, h in zip(*self.get_poses(self.reader.frame)):
            draw_pose(self.player._scene, *pose, c, s, h)

    def draw_positions(self):
        for position, c, s, h in zip(*self.get_positions(self.reader.frame)):
            draw_position(self.player._scene, *position, c, s, h)

    def update_image(self):
        self.trajectory_more.setText('More than {} individual{}'.format(self.options.n_inds, 's' if self.options.n_inds > 1 else ''))
        self.player.clear_drawings()
        if self.options.pose and not self.player.annotating:
            self.draw_poses()
        elif self.options.position and not self.player.annotating:
            self.draw_positions()
        if self.options.annotation and self.file_name in self.annotations and self.reader.frame in self.annotations[self.file_name]:
            self.player.set_annotations(self.annotations[self.file_name][self.reader.frame])
        else:
            self.player.set_annotations(None)

    def update_timeline(self):
        if self.file_name is None and self.tracks is None:
            return
        self.timeline.clear()
        self.timeline.timeseries, self.timeline.identities, self.timeline.colors, self.timeline.selected, self.timeline.hovered = self.get_timeseries(self.reader.frame)
        if self.options.annotation:
            self.timeline.annotations = (self.annotations[self.file_name], self.reader.frame)
        else:
            self.timeline.annotations = ({}, None)
        self.timeline.fitInView()

    def delete_trajectories(self):
        if self.options.interpolated:
            self.status_bar.showMessage('Operation not supported with interpolated tracks.', 2000)
            return
        if len(self.selected) == 0:
            self.status_bar.showMessage('Select at least one individual to delete identities.', 2000)
            return
        identities = sorted(self.selected)
        self.history.add_event('delete', identities, self.reader.frame)
        self.tracks, self.lookup = delete_trajectories(self.tracks, identities, self.lookup)
        self._selected = set()
        self.colors = generate_colors(self.lookup)
        self.update_image()
        self.update_timeline()

    def split_trajectories(self):
        if self.options.interpolated:
            self.status_bar.showMessage('Operation not supported with interpolated tracks.', 2000)
            return
        if len(self.selected) == 0:
            self.status_bar.showMessage('Select at least one individual to split identities.', 2000)
            return
        identities = sorted(self.selected)
        self.history.add_event('split', identities, self.reader.frame)
        self.tracks, self.lookup = split_trajectories(self.tracks, identities, self.reader.frame, self.lookup)
        self._selected = set()
        self.colors = generate_colors(self.lookup)
        self.update_image()
        self.update_timeline()

    def switch_trajectories(self):
        if self.options.interpolated:
            self.status_bar.showMessage('Operation not supported with interpolated tracks.', 2000)
            return
        if len(self.selected) != 2:
            self.status_bar.showMessage('Select two individuals to switch identities.', 2000)
            return
        identities = sorted(self.selected)
        self.history.add_event('switch', identities, self.reader.frame)
        self.tracks, self.lookup = switch_trajectories(self.tracks, identities, self.reader.frame, self.lookup)
        self._selected = set()
        self.colors = generate_colors(self.lookup)
        self.update_image()
        self.update_timeline()

    def merge_trajectories(self):
        if self.options.interpolated:
            self.status_bar.showMessage('Operation not supported with interpolated tracks.', 2000)
            return
        if merge_connected(self):
            return
        if len(self.selected) < 2:
            self.status_bar.showMessage('Select at least two individuals to merge identities.', 2000)
            return
        identities = sorted(self.selected)
        identity = identities[0]
        merge_identities = identities[1:]
        for i in merge_identities:
            if np.isin(self.tracks[str(identity)]['FRAME_IDX'], self.tracks[str(i)]['FRAME_IDX']).any():
                self.status_bar.showMessage('Could not merge all identities due to shared frames.', 2000)
                return
        self.history.add_event('merge', identities, self.reader.frame)
        self.tracks, self.lookup = merge_trajectories(self.tracks, identities, self.lookup)
        self._selected = set()
        self.colors = generate_colors(self.lookup)
        self.update_image()
        self.update_timeline()

    def toggle_interpolated(self, value=None):
        if hasattr(self, 'tracks_copy'):
            self.tracks = self.tracks_copy
            delattr(self, 'tracks_copy')
        else:
            self.tracks, self.tracks_copy = interpolate_tracks(self.tracks)
        if hasattr(self, 'lookup'):
            delattr(self, 'lookup')
        self.lookup = identity_lookup(self.tracks)
        self.colors = generate_colors(self.lookup)
        self.update_image()
        self.update_timeline()
```
