## Supplementary material for "High-resolution animal tracking with integration of environmental information in aquatic systems": Code_track_utility: UI.py

```
import os
import posixpath
import numpy as np
import h5py

import time

from PySide2.QtWidgets import QFileDialog, QShortcut, QMenu, QTimeEdit, QLabel
from PySide2.QtGui import QKeySequence
from PySide2 import QtCore

from copy import deepcopy

from .Reader import Reader
from .Player import Player
from .Settings import Settings
from .Annotator import Annotator
from .Timeline import Timeline
from .History import History
from .QtUtils import points_in_polygons, BorderlessWindow
from .Utils import *
from .Tracks import *

import TrackUtil

def init_shortcuts(tu):
    tu.shortcut_play = QShortcut(QKeySequence(QtCore.Qt.Key_Space), tu, lambda: setattr(tu, 'playing', None))
    tu.shortcut_previous_frame = QShortcut(QKeySequence(QtCore.Qt.Key_Left), tu, tu.previous_frame)
    tu.shortcut_previous_frame.setAutoRepeat(False)
    tu.shortcut_next_frame = QShortcut(QKeySequence(QtCore.Qt.Key_Right), tu, tu.next_frame)
    tu.shortcut_next_frame.setAutoRepeat(False)
    tu.shortcut_draw = QShortcut(QKeySequence(QtCore.Qt.Key_A), tu, lambda: setattr(tu.annotator, 'annotating', None))
    tu.shortcut_draw.setAutoRepeat(False)
    tu.shortcut_cb_pose = QShortcut(QKeySequence(QtCore.Qt.Key_F1), tu, lambda: tu.settings.cb_pose.nextCheckState() \
                                    if tu.settings.cb_pose.isEnabled() else None)
    tu.shortcut_cb_position = QShortcut(QKeySequence(QtCore.Qt.Key_F2), tu, lambda: tu.settings.cb_position.nextCheckState() \
                                    if tu.settings.cb_position.isEnabled() else None)
    tu.shortcut_cb_annotation = QShortcut(QKeySequence(QtCore.Qt.Key_F3), tu, lambda: tu.settings.cb_annotation.nextCheckState() \
                                    if tu.settings.cb_annotation.isEnabled() else None)
    tu.trajectory_clear.setShortcut(QKeySequence(QtCore.Qt.Key_Escape))
    tu.trajectory_clear.setShortcutVisibleInContextMenu(True)
    tu.trajectory_invert.setShortcut(QKeySequence(QtCore.Qt.Key_I))
    tu.trajectory_invert.setShortcutVisibleInContextMenu(True)
    tu.trajectory_delete.setShortcut(QKeySequence(QtCore.Qt.Key_Delete))
    tu.trajectory_delete.setShortcutVisibleInContextMenu(True)
    tu.trajectory_split.setShortcut(QKeySequence(QtCore.Qt.Key_Y))
    tu.trajectory_split.setShortcutVisibleInContextMenu(True)
    tu.trajectory_switch.setShortcut(QKeySequence(QtCore.Qt.Key_X))
    tu.trajectory_switch.setShortcutVisibleInContextMenu(True)
    tu.trajectory_merge.setShortcut(QKeySequence(QtCore.Qt.Key_M))
    tu.trajectory_merge.setShortcutVisibleInContextMenu(True)
    tu.trajectory_new.setShortcut(QKeySequence(QtCore.Qt.Key_E))
    tu.trajectory_new.setShortcutVisibleInContextMenu(True)
    tu.trajectory_lost.setShortcut(QKeySequence(QtCore.Qt.Key_Q))
    tu.trajectory_lost.setShortcutVisibleInContextMenu(True)
    tu.trajectory_more.setShortcut(QKeySequence(QtCore.Qt.Key_O))
    tu.trajectory_more.setShortcutVisibleInContextMenu(True)
    tu.annotation_clear.setShortcut(QKeySequence(QtCore.Qt.Key_Escape))
    tu.annotation_clear.setShortcutVisibleInContextMenu(True)
    tu.annotation_delete.setShortcut(QKeySequence(QtCore.Qt.Key_Delete))
    tu.annotation_delete.setShortcutVisibleInContextMenu(True)
    tu.annotation_next.setShortcut(QKeySequence(QtCore.Qt.Key_E))
    tu.annotation_next.setShortcutVisibleInContextMenu(True)
    tu.annotation_previous.setShortcut(QKeySequence(QtCore.Qt.Key_Q))
    tu.annotation_previous.setShortcutVisibleInContextMenu(True)

def init_menus(tu):
    tu.menu_trajectory = QMenu()
    tu.trajectory_clear = tu.menu_trajectory.addAction('Clear selection')
    tu.trajectory_clear.triggered.connect(lambda: clear_selection(tu))
    tu.trajectory_invert = tu.menu_trajectory.addAction('Invert selection')
    tu.trajectory_invert.triggered.connect(lambda: invert_selection(tu))
    tu.trajectory_delete = tu.menu_trajectory.addAction('Delete selected')
    tu.trajectory_delete.triggered.connect(lambda: delete_selection(tu))
    tu.trajectory_split = tu.menu_trajectory.addAction('Split selected')
    tu.trajectory_split.triggered.connect(tu.split_trajectories)
    tu.trajectory_switch = tu.menu_trajectory.addAction('Switch selected')
    tu.trajectory_switch.triggered.connect(tu.switch_trajectories)
    tu.trajectory_merge = tu.menu_trajectory.addAction('Merge selected')
    tu.trajectory_merge.triggered.connect(tu.merge_trajectories)
    tu.menu_trajectory.addSeparator()
    tu.trajectories_filter = tu.menu_trajectory.addAction('Remove areas')
    tu.menu_trajectory.addSeparator()
    tu.trajectories_filter.triggered.connect(lambda: delete_detections(tu))
    submenu_trajectory = tu.menu_trajectory.addMenu('Navigate')
    tu.trajectory_time = submenu_trajectory.addAction('Insert time')
    tu.trajectory_time.triggered.connect(lambda: jump_time(tu))
    tu.trajectory_new = submenu_trajectory.addAction('New identity')
    tu.trajectory_new.triggered.connect(lambda: jump_next(tu))
    tu.trajectory_lost = submenu_trajectory.addAction('Lost identity')
    tu.trajectory_lost.triggered.connect(lambda: jump_previous(tu))
    tu.trajectory_more = submenu_trajectory.addAction('More than n individuals')
    tu.trajectory_more.triggered.connect(lambda: jump_more_trajectory(tu))
    tu.menu_annotation = QMenu()
    tu.annotation_clear = tu.menu_annotation.addAction('Clear selection')
    tu.annotation_clear.triggered.connect(lambda: clear_selection(tu))
    tu.annotation_delete = tu.menu_annotation.addAction('Delete selected')
    tu.annotation_delete.triggered.connect(lambda: delete_selection(tu))
    tu.menu_annotation.addSeparator()
    submenu_annotation = tu.menu_annotation.addMenu('Navigate')
    tu.annotation_time = submenu_annotation.addAction('Insert time')
    tu.annotation_time.triggered.connect(lambda: jump_time(tu))
    tu.annotation_next = submenu_annotation.addAction('Next annotation')
    tu.annotation_next.triggered.connect(lambda: jump_next(tu))
    tu.annotation_previous = submenu_annotation.addAction('Previous annotation')
    tu.annotation_previous.triggered.connect(lambda: jump_previous(tu))

def init_player(tu):
    tu.player = Player(tu.menu_trajectory, tu.menu_annotation)
    tu.player.signal.identity.connect(lambda i: setattr(tu, 'selected', i))
    tu.player.signal.hovered.connect(lambda v: setattr(tu, 'hovered', v))
    tu.player.signal.annotations.connect(tu.add_annotations)
    tu.player.signal.empty.connect(lambda: tu.status_bar.showMessage('Select at least one annotation to delete.', 2000))
    tu.player.set_annotations(None)

def init_timeline(tu):
    tu.timeline = Timeline(tu.menu_trajectory, tu.menu_annotation, tu.options)
    tu.timeline.signal.identity.connect(lambda i: setattr(tu, 'selected', i))
    tu.timeline.signal.hovered.connect(lambda v: setattr(tu, 'hovered', v))
    tu.timeline.signal.diff.connect(lambda diff: tu.set_frame(tu.reader.frame + diff))

def init_video(tu, select=True):
    if not hasattr(tu, 'reader'):
        tu.reader = Reader(None)
        tu.file_name = None
    if select:
        tu.status_bar.showMessage('Select video.')
        file_name = QFileDialog.getOpenFileName(tu, 'Select video', '', 'All files (*.*)')[0]
        ext = os.path.splitext(file_name)[1]
        img_ext = ['.png', '.PNG', '.jpg', '.JPG']
        if ext in img_ext:
            file_name = os.path.join(os.path.split(file_name)[0], '%*' + ext)
        if os.path.exists(file_name) or ext in img_ext:
            tu.status_bar.showMessage('Initializing video.')
            tu.reader = Reader(file_name)
            tu.timeline.fps = tu.reader.fps
            tu.reader.frame = tu.slider.value()
            tu.file_name = os.path.split(file_name)[1]
            tu.setWindowTitle('TrackUtil --- {}'.format(tu.file_name) + (' --- (connected)' if tu.parent is not None else ''))
            if tu.file_name not in tu.annotations:
                tu.annotations[tu.file_name] = {}
    tu.reader.signal.image.connect(tu.player.setPhoto)
    tu.reader.signal.frame.connect(tu.update_ui)
    tu.reader.signal.toggle.connect(tu.update_buttons)
    if hasattr(tu.reader, 'buffer'):
        tu.reader.buffer.signal.buffered.connect(tu.buffered.setValue)
        tu.button_buffer.clicked.connect(tu.reader.start_buffer)
    tu.status_bar.showMessage('Ready.', 2000)

def init_tracks(tu, select=True):
    if not hasattr(tu, 'tracks'):
        tu.tracks_file = ''
        tu.tracks = {'FRAME_IDX': np.array([], dtype=np.int),
                     'IDENTITIES': np.array([], dtype=np.int)}
    if select:
        tu.status_bar.showMessage('Select tracks.')
        tu.tracks_file = QFileDialog.getOpenFileName(tu, 'Select tracks', '', 'All files (*.pkl)')[0]
    file_name = tu.tracks_file
    if os.path.exists(file_name):
        tu.tracks = load(file_name)
        tu.status_bar.showMessage('Initialized tracks.', 2000)
        tu.reader.frame_count = 0 if tu.tracks['FRAME_IDX'].size == 0 \
                                else max(tu.tracks['FRAME_IDX'].max() + 1, tu.reader.frame_count)
        if 'SPINE' not in tu.tracks[str(tu.tracks['IDENTITIES'][0])]:
            tu.status_bar.showMessage('Initialized tracks, pose rendering disabled.', 2000)
            tu.settings.cb_position.setCheckState(QtCore.Qt.Checked)
            tu.settings.cb_pose.setEnabled(False)
    if hasattr(tu, 'lookup'):
        delattr(tu, 'lookup')
    tu.lookup = identity_lookup(tu.tracks)
    tu.colors = generate_colors(tu.lookup)
    make_backup(tu)

def init_sliders(tu):
    tu.slider_fine.setRange(*get_range(tu.slider.value(), tu.reader.fps * 60, tu.reader.frame_count - 1))
    tu.slider_fine.setTickInterval(tu.reader.fps)
    tu.slider_fine.setValue(tu.slider.value())
    tu.slider_fine.valueChanged.connect(tu.set_frame)
    tu.slider.setRange(0, tu.reader.frame_count - 1)
    tu.slider.setTickInterval(tu.reader.fps * 60)
    tu.slider.setValue(tu.slider_fine.value())
    tu.slider.valueChanged.connect(tu.set_frame)

def init_settings(tu):
    tu.settings = Settings(tu.options)
    tu.settings.cb_pose.stateChanged.connect(lambda v: setattr(tu.options, 'pose', v == 2))
    tu.settings.cb_pose.stateChanged.connect(tu.update_image)
    tu.settings.cb_position.stateChanged.connect(lambda v: setattr(tu.options, 'position', v == 2))
    tu.settings.cb_position.stateChanged.connect(tu.update_image)
    tu.settings.cb_annotation.stateChanged.connect(lambda v: setattr(tu.options, 'annotation', v == 2))
    tu.settings.cb_annotation.stateChanged.connect(tu.update_image)
    tu.settings.cb_annotation.stateChanged.connect(tu.update_timeline)
    tu.settings.cb_annotation.stateChanged.connect(lambda v: setattr(tu.player, 'menus_toggle', v == 0))
    tu.settings.cb_annotation.stateChanged.connect(lambda v: setattr(tu.timeline, 'menus_toggle', v == 0))
    tu.settings.sb_position.valueChanged.connect(lambda v: setattr(tu.options, 'position_size', v))
    tu.settings.sb_position.valueChanged.connect(tu.update_image)
    tu.settings.sb_n_inds.valueChanged.connect(lambda v: setattr(tu.options, 'n_inds', v))
    tu.settings.sb_n_inds.valueChanged.connect(tu.update_image)
    tu.settings.sb_timeline_snapping.valueChanged.connect(lambda v: setattr(tu.options, 'timeline_snapping', v))
    tu.settings.sb_timeline_snapping.valueChanged.connect(lambda v: setattr(tu.timeline, 'timeline_snapping', v))
    tu.settings.cb_interpolate.stateChanged.connect(lambda v: setattr(tu.options, 'interpolated', v == 2))
    tu.settings.cb_interpolate.stateChanged.connect(tu.toggle_interpolated)
    tu.settings.cb_connect.stateChanged.connect(lambda v: setattr(tu.options, 'connected', v == 2))
    tu.settings.cb_connect.stateChanged.connect(lambda: update_connected(tu))
    tu.settings.cb_fix.stateChanged.connect(lambda v: setattr(tu.options, 'fix', v == 2))
    tu.button_settings.clicked.connect(lambda: tu.settings.setVisible(True))
    tu.button_settings.clicked.connect(lambda: tu.settings.activateWindow())
    tu.button_settings.clicked.connect(lambda: tu.settings.showNormal())

def init_history(tu):
    tu.history = History()
    tu.button_history.clicked.connect(lambda: tu.history.setVisible(True))
    tu.button_history.clicked.connect(lambda: tu.history.activateWindow())
    tu.button_history.clicked.connect(lambda: tu.history.showNormal())
    tu.history.signal.history.connect(lambda v: set_history(tu, v))
    tu.history.button_make_backup.clicked.connect(lambda: make_backup(tu))
    tu.history.button_load_backup.clicked.connect(lambda: load_backup(tu))

def init_annotations(tu, select=True):
    if not hasattr(tu, 'annotations'):
        tu.annotations = {'class_names': []}
    if select:
        tu.status_bar.showMessage('Select annotations.')
        file_name = QFileDialog.getOpenFileName(tu, 'Select annotations', '', 'All files (*.pkl)')[0]
        if os.path.exists(file_name):
            annotations = load(file_name)
            if 'class_names' in annotations:
                tu.annotations = annotations
                for class_name in annotations['class_names']:
                    tu.annotator.add_class(class_name=class_name)
                if tu.file_name is not None and tu.file_name not in tu.annotations:
                    tu.annotations[tu.file_name] = {}
    tu.status_bar.showMessage('Ready.', 2000)

def init_annotator(tu):
    tu.annotator = Annotator(tu.menu_annotation)
    for class_name in tu.options.class_names:
        tu.annotator.add_class(class_name=class_name)
    tu.annotator.signal.annotating.connect(lambda v: setattr(tu.player, 'annotating', v))
    tu.annotator.signal.class_name.connect(lambda v: setattr(tu.player, 'class_name', v))
    tu.annotator.signal.class_name.connect(tu.options.add_class)
    tu.annotator.signal.class_name.connect(lambda: update_connected(tu))
    tu.annotator.signal.remove.connect(tu.options.remove_class)
    tu.annotator.signal.remove.connect(lambda: update_connected(tu))
    tu.player.signal.closed.connect(lambda v: setattr(tu.annotator, 'annotating', v))
    tu.button_annotator.clicked.connect(lambda: tu.annotator.setVisible(True))
    tu.button_annotator.clicked.connect(lambda: tu.annotator.activateWindow())
    tu.button_annotator.clicked.connect(lambda: tu.annotator.showNormal())
    tu.annotator.signal.annotating.connect(lambda: tu.settings.cb_annotation.setCheckState(QtCore.Qt.Checked))

def load_video(tu):
    resume = False
    if tu.reader.running:
        resume = True
        tu.playing = not tu.playing
    init_video(tu)
    init_sliders(tu)
    tu.update_buttons()
    tu.set_frame(tu.reader.frame)
    tu.player.fitInView(upscale=True)
    if resume:
        tu.playing = not tu.playing

def load_tracks(tu):
    resume = False
    if tu.reader.running:
        resume = True
        tu.playing = not tu.playing
    init_tracks(tu)
    init_sliders(tu)
    tu.set_frame(tu.reader.frame)
    if resume:
        tu.playing = not tu.playing

def load_annotations(tu):
    resume = False
    if tu.reader.running:
        resume = True
        tu.playing = not tu.playing
    init_annotations(tu)
    init_sliders(tu)
    tu.set_frame(tu.reader.frame)
    if resume:
        tu.playing = not tu.playing

def save_tracks(tu):
    file_name = QFileDialog.getSaveFileName(tu, 'Save file', '', 'All files (*.pkl)')[0]
    if os.path.exists(os.path.split(file_name)[0]):
        save(dump=tu.tracks, file_name=file_name)
        tu.status_bar.showMessage('Saved tracks to {}.'.format(file_name), 2000)
    else:
        tu.status_bar.showMessage('Did not save tracks.'.format(file_name), 2000)

def save_annotations(tu):
    file_name = QFileDialog.getSaveFileName(tu, 'Save file', '', 'All files (*.pkl)')[0]
    if os.path.exists(os.path.split(file_name)[0]):
        tu.annotations['class_names'] = tu.options.class_names
        save(dump=tu.annotations, file_name=file_name)
        tu.status_bar.showMessage('Saved annotations to {}.'.format(file_name), 2000)
    else:
        tu.status_bar.showMessage('Did not save annotations.', 2000)

def write_dataset(tu):
    if tu.file_name is None:
        tu.status_bar.showMessage('Requires video.', 2000)
        return
    elif len(tu.annotations[tu.file_name]) == 0:
        tu.status_bar.showMessage('No annotations for current video.', 2000)
        return
    tu.reader.video = True
    if tu.reader.running:
        tu.playing = not tu.playing
    file_name = QFileDialog.getSaveFileName(tu, 'Write dataset', '', 'All files (*.h5)')[0]
    if os.path.exists(os.path.split(file_name)[0]):
        tu.settings.cb_annotation.setCheckState(QtCore.Qt.Checked)
        tu.job = (file_name, list(sorted(tu.annotations[tu.file_name].keys())), QtCore.QTimer(tu))
        tu.job[2].timeout.connect(lambda: write_annotation(tu))
        tu.job[2].start()

def write_annotation(tu):
    tu.buffered.setValue(100 - 100 * len(tu.job[1]) // len(tu.annotations[tu.file_name]))
    if len(tu.job[1]) == 0:
        tu.job[2].stop()
        with h5py.File(tu.job[0], 'a') as h5_file:
            annotations = []
            for file_name in h5_file:
                if file_name != 'annotations':
                    annotations.extend([posixpath.join(file_name, str(frame)) for frame in sorted(h5_file[file_name])])
            annotations = np.array(annotations).astype(np.bytes_)
            if 'annotations' not in h5_file:
                h5_file.create_dataset('annotations', data=annotations, dtype='|S400', maxshape=(None,), compression='gzip', compression_opts=9)
            else:
                h5_file['annotations'].resize(annotations.shape)
                h5_file['annotations'][:] = annotations
        tu.status_bar.showMessage('Ready.', 2000)
        return
    with h5py.File(tu.job[0], 'a') as h5_file:
        if tu.file_name not in h5_file:
            dataset = h5_file.create_group(tu.file_name)
        else:
            dataset = h5_file[tu.file_name]
        frame = tu.job[1].pop(0)
        img = tu.set_frame(frame, from_video=True)
        mask, class_names = write_mask(tu.annotations[tu.file_name][frame], *img.shape[:2])
        tu.status_bar.showMessage('Writing dataset {}.'.format(tu.job[0]), 2000)
        if str(frame) in dataset:
            dataset[str(frame)]['image'][:] = img
            dataset[str(frame)]['mask'].resize(mask.shape)
            dataset[str(frame)]['mask'][:] = mask
            dataset[str(frame)]['class_names'].resize(class_names.shape)
            dataset[str(frame)]['class_names'][:] = class_names
        else:
            annotation = dataset.create_group(str(frame))
            annotation.create_dataset('image', data=img, dtype=np.uint8, compression='gzip', compression_opts=9)
            annotation.create_dataset('mask', data=mask, dtype=np.bool, maxshape=(*img.shape[:2], None,), compression='gzip', compression_opts=9)
            annotation.create_dataset('class_names', data=class_names, dtype='|S200', maxshape=(None,), compression='gzip', compression_opts=9)

def jump_previous(tu):
    previous = np.array([])
    if tu.player.menus_toggle and tu.lookup.shape[0] > 0:
        frames = np.arange(tu.lookup.shape[0])
        last = tu.lookup.shape[0] - np.argmax(np.flip(tu.lookup, axis=0), axis=0) - 1
        previous = last[frames[last] < tu.reader.frame]
    elif not tu.player.menus_toggle and tu.file_name in tu.annotations:
        annotated_frames = np.array(list(tu.annotations[tu.file_name].keys()))
        previous = annotated_frames[annotated_frames < tu.reader.frame]
    if previous.size == 0:
        tu.status_bar.showMessage('No {} found before frame {}.'.format('discontinued trajectory' if tu.player.menus_toggle else 'annotation',
                                                                        tu.reader.frame), 2000)
    else:
        tu.set_frame(previous.max())

def jump_next(tu):
    next = np.array([])
    if tu.player.menus_toggle and tu.lookup.shape[0] > 0:
        frames = np.arange(tu.lookup.shape[0])
        first = np.argmax(tu.lookup, axis=0)
        next = first[frames[first] > tu.reader.frame]
    elif not tu.player.menus_toggle and tu.file_name in tu.annotations:
        annotated_frames = np.array(list(tu.annotations[tu.file_name].keys()))
        next = annotated_frames[annotated_frames > tu.reader.frame]
    if next.size == 0:
        tu.status_bar.showMessage('No {} found after frame {}.'.format('new trajectory' if tu.player.menus_toggle else 'annotation',
                                                                       tu.reader.frame), 2000)
    else:
        tu.set_frame(next.min())

def jump_more_trajectory(tu):
    frames = np.arange(tu.lookup.shape[0])
    more = np.array([])
    if tu.lookup.size > 0:
        more = np.sum(tu.lookup, axis=1) > tu.options.n_inds
        more = np.argwhere(more).ravel()
    if more.size == 0:
        tu.status_bar.showMessage('Never more than {} individual{}.'.format(tu.options.n_inds, 's' if tu.options.n_inds > 1 else ''), 2000)
    else:
        next = more[frames[more] > tu.reader.frame]
        if next.size == 0:
            frame = frames[more[0]]
        else:
            frame = frames[next[0]]
        tu.set_frame(frame)

def time_to_frame(tu, time):
    msecs = time.msecsSinceStartOfDay()
    return int(np.round(msecs * tu.reader.fps / 1000))

def jump_time(tu):
    if not hasattr(tu, 'time_widget'):
        time_widget = BorderlessWindow(main_window=False)
        time_widget.setWindowTitle('Insert Time')
        layout = time_widget.layout()
        time_widget.time = QTimeEdit()
        time_widget.time.setDisplayFormat('hh:mm:ss.zzz')
        layout.addWidget(time_widget.time)
        tu.time_widget = time_widget
        tu.time_widget.time.timeChanged.connect(lambda: tu.set_frame(time_to_frame(tu, tu.time_widget.time.time())))
        tu._close.clicked.connect(tu.time_widget.close)
    tu.time_widget.show()

def make_backup(tu):
    save([tu.tracks, tu.lookup], '.backup')
    tu.history.clear_events()

def load_backup(tu):
    tu.tracks, tu.lookup = load('.backup')
    tu.colors = generate_colors(tu.lookup)
    tu.update_image()
    tu.update_timeline()
    tu.history.clear_events()

def set_history(tu, history):
    tu.tracks, tu.lookup = load('.backup')
    if len(history) == 0:
        return
    for kind, selected, frame in history:
        if kind == 'delete':
            tu.tracks, tu.lookup = delete_trajectories(tu.tracks, selected, tu.lookup)
        elif kind == 'split':
            tu.tracks, tu.lookup = split_trajectories(tu.tracks, selected, frame, tu.lookup)
        elif kind == 'merge':
            tu.tracks, tu.lookup = merge_trajectories(tu.tracks, selected, tu.lookup)
        elif kind == 'switch':
            tu.tracks, tu.lookup = switch_trajectories(tu.tracks, selected, frame, tu.lookup)
    tu.colors = generate_colors(tu.lookup)
    tu.set_frame(frame)

def clear_selection(tu):
    if tu.player.menus_toggle:
        tu._selected = set()
        tu.update_image()
        tu.update_timeline()
    else:
        for occlusion_level, polygons in enumerate(tu.player.occluded):
            for idx, p in enumerate(polygons):
                tu.player.selected[occlusion_level][idx] = False
        tu.player.draw_annotations()

def invert_selection(tu):
    if tu.player.menus_toggle:
        tu._selected = set(tu.tracks['IDENTITIES']) - tu.selected
        tu.update_image()
        tu.update_timeline()
    else:
        pass # this would invert annotations per frame?

def delete_selection(tu):
    if tu.player.menus_toggle:
        tu.delete_trajectories()
    else:
        tu.player.delete_annotations()

def open_child(tu):
    parent = tu if tu.parent is None else tu.parent
    child = TrackUtil.TrackUtil(parent=parent)
    for class_name in parent.options.class_names:
        child.annotator.add_class(class_name)
    child.annotations = parent.annotations
    child.setVisible(True)
    parent.children.append(child)

def merge_connected(source):
    if source.parent is None:
        connected = [source] + [child for child in source.children]
    else:
        connected = [source.parent] + [child for child in source.parent.children]
    if len(connected) == 1:
        return False
    check_selected = np.array([len(tu.selected) for tu in connected])
    if np.sum(check_selected > 0) == 1:
        return False
    if np.any(check_selected > 1):
        for tu in connected:
            tu.status_bar.showMessage('Select exactly one individual to merge connected.', 2000)
        return False
    selected = [tu.selected.pop() if tu.selected else None for tu in connected]
    connected = [tu for s, tu in zip(selected, connected) if s is not None]
    selected = [s for s in selected if s is not None]
    fixed = np.array([tu.options.fix for tu in connected])
    if np.sum(fixed) > 1:
        for tu in connected:
            tu.update_image()
            tu.update_timeline()
            tu.status_bar.showMessage('More than one fixed identity, could not merge connected.', 2000)
        return False
    if np.sum(fixed) == 1:
        identity = selected[np.argwhere(fixed).ravel()[0]]
        if identity in np.concatenate([tu.tracks['IDENTITIES'] for fix, tu in zip(fixed, connected) if not fix]):
            for tu in connected:
                tu.update_image()
                tu.update_timeline()
                tu.status_bar.showMessage('Could not assign fixed identity to same identity.', 2000)
            return False
    else:
        identity = np.random.randint(10000, 100000)
        while identity in np.concatenate([tu.tracks['IDENTITIES'] for tu in connected]):
            identity = np.random.randint(10000, 100000)
    for i, tu in zip(selected, connected):
        if i is not None:
            tu.tracks[str(identity)] = tu.tracks.pop(str(i))
            tu.tracks['IDENTITIES'][tu.tracks['IDENTITIES'] == i] = identity
            tu.update_image()
            tu.update_timeline()
    return True

def update_connected(tu, source=None):
    if tu.parent is None:
        if source is not None:
            update_classes(tu, set(source.annotator.classes))
            update_settings(tu, source.options)
        for child in tu.children:
            if child is source:
                continue
            update_classes(child, set(tu.annotator.classes))
            update_settings(child, tu.options)
    else:
        update_connected(tu.parent, source=tu)

def update_classes(tu, class_names):
    add_classes = class_names - set(tu.options.class_names)
    remove_classes = set(tu.options.class_names) - class_names
    for class_name in add_classes:
            tu.annotator.add_class(class_name, silent=True)
            tu.options.add_class(class_name)
    for class_name in remove_classes:
            tu.annotator.remove_class(class_name, silent=True)
            tu.options.remove_class(class_name)

def update_settings(tu, options):
    tu.options.connected = options.connected
    state = QtCore.Qt.Checked if options.connected else QtCore.Qt.Unchecked
    tu.settings.cb_connect.blockSignals(True)
    tu.settings.cb_connect.setCheckState(state)
    tu.settings.cb_connect.blockSignals(False)

def delete_detections(tu):
    polygons = []
    if tu.file_name in tu.annotations:
        for frame in tu.annotations[tu.file_name]:
            annotation = tu.annotations[tu.file_name][frame]
            for occlusion_level in annotation:
                polygons.extend([polygon for polygon in annotation[occlusion_level]['occluded']])
    if len(polygons) == 0:
        return
    pts = np.concatenate([np.transpose([tu.tracks[str(i)]['X'], tu.tracks[str(i)]['Y']]) \
                          for i in tu.tracks['IDENTITIES']])
    if pts.size == 0:
        return
    in_polygons = points_in_polygons(pts, polygons)
    for i in range(tu.lookup.shape[1]):
        identity = tu.tracks['IDENTITIES'][i]
        num_detections = tu.lookup[:, i].sum()
        in_polygons_trajectory = in_polygons[:num_detections]
        # complicated condition necessary because double detection in faulty trajectories
        condition = np.isin(tu.tracks[str(identity)]['FRAME_IDX'], np.unique(tu.tracks[str(identity)]['FRAME_IDX'])[np.invert(in_polygons_trajectory)])
        tu.tracks[str(identity)] = filter_trajectory(tu.tracks[str(identity)], condition) # np.invert(in_polygons_trajectory)
        in_polygons = in_polygons[num_detections:]
    tu.tracks = update_frames(tu.tracks)
    if hasattr(tu, 'lookup'):
        delattr(tu, 'lookup')
    tu.lookup = identity_lookup(tu.tracks)
    tu.tracks, tu.lookup = delete_trajectories(tu.tracks, tu.tracks['IDENTITIES'][tu.lookup.sum(axis=0) == 0], tu.lookup)
    tu.colors = generate_colors(tu.lookup)
    tu.update_image()
    tu.update_timeline()
```
