## Supplementary material for "High-resolution animal tracking with integration of environmental information in aquatic systems": Code_track_utility: Utils.py

```
import _pickle
import cv2
import numpy as np
from seaborn import color_palette
from scipy.signal import savgol_filter

import time

def save(dump, file_name):
    with open(file_name, 'wb') as fid:
        _pickle.dump(dump, fid)
    return True

def load(file_name):
    with open(file_name, 'rb') as fid:
        dump = _pickle.load(fid)
    return dump

def in_range(value, minimum, maximum):
    if value >= minimum and value <= maximum:
        return True
    return False

def get_range(value, duration, total):
    minimum = value - (value % duration)
    maximum = minimum + duration - 1
    if maximum > total:
        maximum = total
    return minimum, maximum

def generate_colors(lookup):
    if lookup.size == 0:
        return []
    colors = np.array(color_palette('pastel')) * 255
    color_idx = np.arange(lookup.shape[1]) % len(colors)
    fixed_colors = np.zeros((lookup.shape[1], 3))
    for idx in range(len(colors)):
        fixed_colors[color_idx == idx] = colors[idx]
    return fixed_colors

def diverging_hsv(colors):
    if np.array(colors).size == 0:
        return (np.random.randint(0, 359), 255, 255)
    hues = np.array(sum(colors, []))[:, 0]
    if hues.size == 1:
        return (359 - hues[0], 255, 255)
    else:
        hues = np.sort(hues)
        diff = np.append(np.diff(hues), 360 - hues[-1] + hues[0])
        idx = np.argmax(diff)
        hue = hues[idx] + diff[idx] // 2
        hue = hue - 360 if hue > 359 else hue
        return (hue, 255, 255)

def write_mask(annotation, height, width):
    layers = []
    class_names = []
    occlusion = np.zeros((height, width), dtype=np.bool)
    for occlusion_level in sorted(annotation):
        for p, c in zip(annotation[occlusion_level]['polygons'][::-1],
                        annotation[occlusion_level]['class_names'][::-1]):
            layer = np.zeros((height, width, 3), dtype=np.uint8)
            cv2.fillPoly(layer, [p.reshape(-1, 1, 2).astype(np.int)], (255, 255, 255), cv2.LINE_AA)
            layer = np.sum(layer, axis=2) > 0
            layers.append(layer & np.invert(occlusion))
            class_names.append(c)
            occlusion = occlusion | layer
    mask = np.stack(layers, axis=2)
    non_empty = mask.sum(axis=(0, 1)) > 0
    class_names = np.array(class_names).astype(np.bytes_)[non_empty]
    mask = mask[:, :, non_empty]
    return mask, class_names
```
