## Supplementary material for "High-resolution animal tracking with integration of environmental information in aquatic systems": Code_training_inference: readme_training-inference.txt

install a conda or miniconda environment with jupyter notebook to open the interactive python notebook (.ipynb)
follow installation instructions of the original mask rcnn repository (https://github.com/matterport/Mask_RCNN)
