## Supplementary material for "High-resolution animal tracking with integration of environmental information in aquatic systems": Code_training_inference: training-inference.ipynb

```
# %% [markdown]
The following notebook was used for training and inference in case of the *single* dataset (tracking the european eal *Conger conger*)

# %% [code]
# import everything and set paths to coco weights, log dir

import os
import sys
import random
import math
import re
import time
import numpy as np
import cv2
import matplotlib
import matplotlib.pyplot as plt
import h5py
from imgaug import augmenters as iaa
from imgaug import parameters as iap
from glob import glob

# Root directory of the project
ROOT_DIR = os.path.abspath('/home/jordanlab/Mask_RCNN')

# Import Mask RCNN
sys.path.append(ROOT_DIR)  # To find local version of the library
from mrcnn.config import Config
from mrcnn import utils
import mrcnn.model as modellib
from mrcnn import visualize
from mrcnn.model import log

# Directory to save logs and trained model
MODEL_DIR = os.path.join(ROOT_DIR, "logs")

# Local path to trained weights file
COCO_MODEL_PATH = os.path.join(ROOT_DIR, "mask_rcnn_coco.h5")
# Download COCO trained weights from Releases if needed
if not os.path.exists(COCO_MODEL_PATH):
    utils.download_trained_weights(COCO_MODEL_PATH)
    
# Following is needed for cudnn when continuing with different resolution
import tensorflow as tf
config = tf.ConfigProto()
config.gpu_options.allow_growth = True
session = tf.Session(config=config)

# for visualization
def get_ax(rows=1, cols=1, size=8):
    """Return a Matplotlib Axes array to be used in
    all visualizations in the notebook. Provide a
    central point to control graph sizes.
    
    Change the default size attribute to control the size
    of rendered images
    """
    _, ax = plt.subplots(rows, cols, figsize=(size*cols, size*rows))
    return ax

# %% [code]
# replicate the classes used in TrackUtil annotations and create a config instance used for training

classes = {1: 'eel'}

class Config(Config):
    NAME = 'eel'

    GPU_COUNT = 1
    IMAGES_PER_GPU = 1

    NUM_CLASSES = len(classes) + 1  # background + number of classes

    IMAGE_MIN_DIM = 1024 # set resolution for training (image downsampling)
    IMAGE_MAX_DIM = 1024
    
    IMAGE_RESIZE_MODE = 'square'

    # Use smaller anchors because our image and objects are small
    RPN_ANCHOR_SCALES = (16, 32, 128, 256, 512)

    # Reduce training ROIs per image because the images are small and have
    # few objects. Aim to allow ROI sampling to pick 33% positive ROIs.
    TRAIN_ROIS_PER_IMAGE = 128

    # Use a small epoch since the data is simple
    STEPS_PER_EPOCH = 200

    # use small validation steps since the epoch is small
    VALIDATION_STEPS = 50
    
config = Config()
config.display()

# %% [code]
# define our custom Dataset class that loads the datasets exported with TrackUtil

from multiprocessing import Lock, Value, Pool, cpu_count
import tqdm

def load_images(i, images, annotation_file, temp):
    path = os.path.join(images[i], 'image')
    img_path = os.path.join(temp, os.path.split(path)[0].replace('/', '-').replace(':', '') + '.png')
    if os.path.exists(img_path):
        height, width = cv2.imread(img_path).shape[:2]
    else:
        with h5py.File(annotation_file, 'r') as h5_file:
            height, width = h5_file[path][:].shape[:2]
            image = h5_file[path][:]
            cv2.imwrite(img_path, image[:, :, ::-1])        
    lock.acquire()
    count.value += 1
    print('Reading images: {:.2f} %'.format(100 * count.value / len(images)),
          sep=' ', end='\r' if count.value < len(images) else '\n', flush=True)
    lock.release()
    return (i, path, width, height)

def init_pool(l, c):
    global lock, count
    lock = l
    count = c

class Dataset(utils.Dataset):

    def load_images(self, annotation_file, classes, source, mode='full'):
        self.classes = classes
        self.annotation_file = annotation_file
        self.temp = os.path.join(os.path.join(os.path.dirname(annotation_file), os.path.split(annotation_file)[-1] + 'temp'))
        if not os.path.exists(self.temp):
            os.mkdir(self.temp)
        with h5py.File(self.annotation_file, 'r') as h5_file:
            for c in self.classes:
                self.add_class(source, c, self.classes[c])
            self.images = h5_file['annotations'][:].astype(np.str)
        np.random.seed(0)
        image_idx = np.arange(self.images.size)
        val_idx = np.random.choice(image_idx, image_idx.size // 6)
        if mode == 'train':
            self.images = self.images[np.invert(np.isin(image_idx, val_idx))]
        elif mode == 'val':
            self.images = self.images[np.isin(image_idx, val_idx)]
        else:
            print('Warning: set mode to "train" or "val", otherwise using full dataset')
        
        l = Lock()
        c = Value('d', 0)
        with Pool(processes=cpu_count(), initializer=init_pool, initargs=(l, c)) as pool:
            to_add = pool.starmap(load_images, list(zip(np.arange(len(self.images)),
                                                        [self.images] * len(self.images),
                                                        [self.annotation_file] * len(self.images),
                                                        [self.temp] * len(self.images))))
        for (i, path, width, height) in to_add:
            self.add_image(source, image_id=i, path=path, width=width, height=height)

    def load_image(self, image_id):
        info = self.image_info[image_id]
        path = info['path']
        img_path = os.path.join(self.temp, os.path.split(path)[0].replace('/', '-').replace(':', '') + '.png')
        image = cv2.imread(img_path)[:, :, ::-1]
        return image

    def load_mask(self, image_id):
        info = self.image_info[image_id]
        img_path = info['path']
        path = os.path.split(img_path)[0]
        with h5py.File(self.annotation_file, 'r') as h5_file:
            mask = h5_file[os.path.join(path, 'mask')][:]
            classes = h5_file[os.path.join(path, 'class_names')][:].astype(np.str)
        use = np.array([idx for idx, name in enumerate(classes) for c in self.classes if name in self.classes[c].split(',')], dtype=np.int32)
        class_ids = np.array([c for name in classes for c in self.classes if name in self.classes[c].split(',')], dtype=np.int32)
        mask = mask[:, :, use]
        non_empty = mask.sum(axis=(0, 1)) > 10
        return mask[:, :, non_empty], class_ids[non_empty]
    
    def prepare(self):
        super().prepare()
        print('{} images, classes: '.format(len(self.image_ids)), *['[{}: {}]'.format(idx, self.classes[idx]) for idx in self.classes])

# %% [code]
# use image augmentations via the imgaug package to avoid overfitting

augmentation = iaa.Sequential([   
    iaa.Fliplr(0.5),
    iaa.Flipud(0.5),
    iaa.Affine(rotate=iap.Choice([0,90,180,270])),
    iaa.Affine(scale={'x': (0.8, 1.2), 'y': (0.8, 1.2)},
                      translate_percent={'x': (-0.2, 0.2), 'y': (-0.2, 0.2)},
                      rotate=(-45, 45))
    ], random_order=True)

# %% [code]
# prepare training and validation datasets, set paths accordingly

# Training dataset
dataset_train = Dataset()
dataset_train.load_images(annotation_file='/home/jordanlab/Pictures/annotation/paul/eel_refined_2.h5',
                          classes=classes,
                          source='eel',
                          mode = 'train')
dataset_train.prepare()

# Validation dataset
dataset_val = Dataset()
dataset_val.load_images(annotation_file='/home/jordanlab/Pictures/annotation/paul/eel_refined_2.h5',
                        classes=classes,
                        source='eel',
                        mode = 'val') 
dataset_val.prepare()

# %% [code]
# visualize some annotations

image_ids = dataset_val.image_ids[:5]
for image_id in image_ids:
    image = dataset_val.load_image(image_id)
    mask, class_ids = dataset_val.load_mask(image_id)
    visualize.display_top_masks(image, mask, class_ids, dataset_val.class_names, limit=1)

# %% [code]
# create a mask rcnn model instance, load coco weights

model = modellib.MaskRCNN(mode='training', config=config, model_dir=MODEL_DIR)

# Which weights to start with?
init_with = 'coco'  # coco, or last

if init_with == 'coco':
    model.load_weights(COCO_MODEL_PATH, by_name=True,
                       exclude=['mrcnn_class_logits', 'mrcnn_bbox_fc', 
                                'mrcnn_bbox', 'mrcnn_mask'])
elif init_with == 'last':
    model.load_weights(model.find_last(), by_name=True) # model.find_last()

# %% [code]
# first train head layers

model.train(dataset_train, dataset_val,
            learning_rate=config.LEARNING_RATE, 
            epochs=60, 
            layers='heads',
            augmentation=augmentation)

# %% [code]
# and then the full model

model.train(dataset_train, dataset_val, 
            learning_rate=config.LEARNING_RATE / 10,
            epochs=140,
            layers='all',
            augmentation=augmentation)

# %% [code]
# create a config instance used for inference

class InferenceConfig(Config):
    GPU_COUNT = 1
    IMAGES_PER_GPU = 1
    
    IMAGE_MAX_DIM = 1024
    IMAGE_MIN_DIM = 1024
    
    IMAGE_RESIZE_MODE = 'square'

inference_config = InferenceConfig()
batch_size = inference_config.BATCH_SIZE

# %% [code]
# load the last training weight for inference

model = modellib.MaskRCNN(mode="inference", 
                          config=inference_config,
                          model_dir=MODEL_DIR)

model.load_weights(model.find_last(), by_name=True) # 72

# %% [code]
# visualize inference on some images of the validation set

threshold = 0.0

image_ids = np.random.choice(dataset_val.image_ids, 20)
for image_id in image_ids:
    original_image, image_meta, gt_class_id, gt_bbox, gt_mask =\
        modellib.load_image_gt(dataset_val, inference_config, 
                               image_id, use_mini_mask=False)

    fig, ax = plt.subplots(1, 2, figsize=(32, 32))

    visualize.display_instances(original_image, gt_bbox, gt_mask, gt_class_id, 
                                dataset_train.class_names, figsize=(16, 16), ax=ax[0])

    results = model.detect([original_image], verbose=0)

    r = results[0]
    visualize.display_instances(original_image, r['rois'][r['scores'] > threshold], r['masks'][:, :, r['scores'] > threshold], r['class_ids'][r['scores'] > threshold], 
                                dataset_val.class_names, r['scores'][r['scores'] > threshold], ax=ax[1])

    plt.show()

# %% [code]
# define some functions for inference and visualization

def generate_tiles(img, size=512, overlap=100, shifts=1, as_list=False):
    height, width = img.shape[:2]
    origins = np.mgrid[0:height:(size // shifts - overlap),0:width:(size // shifts - overlap)].T.reshape(-1,2)
    imgs = []
    for tl in origins:
        tile = img[tl[0]:(tl[0] + size), tl[1]:(tl[1] + size)]
        if tile.shape[0] < size and tile.shape[1] == size:
            tile = img[(height - size):height, tl[1]:(tl[1] + size)]
        elif tile.shape[0] == size and tile.shape[1] < size:
            tile = img[tl[0]:(tl[0] + size), (width - size):width]
        elif tile.shape[0] < size and tile.shape[1] < size:
            tile = img[(height - size):height, (width - size):width]
        imgs.append(tile)
    if as_list:
        return imgs
    return np.stack(imgs, axis=2)

def stitch_tiles(tiles, target_shape, size=512, overlap=100, shifts=1, flatten=False):
    height, width = target_shape[:2]
    origins = np.mgrid[0:height:(size // shifts - overlap),0:width:(size // shifts - overlap)].T.reshape(-1,2)
    img = np.zeros((height, width, np.sum([tile.shape[2] for tile in tiles])), dtype=np.uint8)
    idx = 0
    for tile, tl in zip(tiles, origins):
        if tl[0] + size > height:
            tl[0] = height - size
        if tl[1] + size > width:
            tl[1] = width - size
        img[tl[0]:(tl[0] + size), tl[1]:(tl[1] + size), idx:(idx + tile.shape[2])] = tile
        idx += tile.shape[2]
    if flatten:
        img = img.sum(axis=(2)) > 0
    return img

def stitch_rois(rois, target_shape, size=512, overlap=100, shifts=1):
    height, width = target_shape[:2]
    origins = np.mgrid[0:height:(size // shifts - overlap),0:width:(size // shifts - overlap)].T.reshape(-1,2)
    for idx, (roi, tl) in enumerate(zip(rois, origins)):
        if tl[0] + size > height:
            tl[0] = height - size
        if tl[1] + size > width:
            tl[1] = width - size
        rois[idx][:, ::2] += tl[0]
        rois[idx][:, 1::2] += tl[1]
    rois = np.concatenate(rois)
    return rois

def stitch_predictions(results, target_shape, size=512, overlap=100, shifts=1):
    stitched_result = {}
    stitched_result['masks'] = stitch_tiles([result['masks'] for result in results], target_shape, size, overlap, shifts)
    stitched_result['rois'] = stitch_rois([result['rois'] for result in results], target_shape, size, overlap, shifts)
    stitched_result['scores'] = np.concatenate([result['scores'] for result in results])
    stitched_result['class_ids'] = np.concatenate([result['class_ids'] for result in results])
    return stitched_result

import colorsys
from glob import glob
import h5py
import skvideo.io

def split_image(image, overlap=1/6):
    shape = image.shape[:2]
    overlap = round(min(shape) * overlap / 2)
    ul = np.concatenate([np.array([0, 0]), np.array(shape) / 2 + overlap]).astype(np.int)
    ur = np.array([0, shape[1] / 2 - overlap, shape[0] / 2 + overlap, shape[1]]).astype(np.int)
    bl = np.array([shape[0] / 2 - overlap, 0, shape[0], shape[1] / 2 + overlap]).astype(np.int)
    br = np.concatenate([np.array(shape) / 2 - overlap, shape]).astype(np.int)
    ul = image[ul[0]:ul[2], ul[1]:ul[3]]
    ur = image[ur[0]:ur[2], ur[1]:ur[3]]
    bl = image[bl[0]:bl[2], bl[1]:bl[3]]
    br = image[br[0]:br[2], br[1]:br[3]]
    return [ul, ur, bl, br]

def random_colors(N, bright=True):
    """
    Generate random colors.
    To get visually distinct colors, generate them in HSV space then
    convert to RGB.
    """
    brightness = 1.0 if bright else 0.7
    hsv = [(i / N, 1, brightness) for i in range(N)]
    colors = list(map(lambda c: colorsys.hsv_to_rgb(*c), hsv))
    random.shuffle(colors)
    return colors


def apply_mask(image, mask, color, alpha=0.25):
    """Apply the given mask to the image.
    """
    for c in range(3):
        image[:, :, c] = np.where(mask == 1,
                                  image[:, :, c] *
                                  (1 - alpha) + alpha * color[c] * 255,
                                  image[:, :, c])
        image = image.astype(np.uint8)
    return image


def draw_instances(image, boxes, masks, class_ids, class_names, scores=None):
    if len(image.shape) > 2 and image.shape[2] == 1:
        image = np.stack([image.reshape(*image.shape[:2])] * 3, axis=2)
    N = boxes.shape[0]
    if not N:
        return image
    else:
        colors = random_colors(N)
        height, width = image.shape[:2]
        masked_image = image.astype(np.uint32).copy()
        for i in range(N):
            color = colors[i]
            if not np.any(boxes[i]):
                continue
            y1, x1, y2, x2 = boxes[i]
            class_id = class_ids[i]
            score = scores[i] if scores is not None else None
            label = class_names[class_id]
            caption = "{} {:.3f}".format(label, score) if score else label
            mask = masks[:, :, i]
            masked_image = apply_mask(masked_image, mask, color)
            color = [int(c) * 255 for c in color]
#             cv2.rectangle(masked_image, (x1, y1), (x2, y2), color, 1, cv2.LINE_AA)
            cv2.putText(masked_image, caption, (x1, y1 - 8), cv2.FONT_HERSHEY_SIMPLEX, 0.5, color, 1, cv2.LINE_AA)
    return masked_image

# %% [code]
# run inference on videos after setting paths accordingly

import skvideo.io

threshold = 0.0

video_path = '/home/jordanlab/Pictures/annotation/paul/'
output_path = '/home/jordanlab/Pictures/annotation/paul/'

if not os.path.exists(output_path):
    os.makedirs(output_path)

video_files = sorted(glob(os.path.join(video_path, '*.mp4')))

for video_file in video_files:
    
    queue = []
    images = []
    results = []
    frame_idx = 0

    video_name = os.path.splitext(os.path.basename(video_file))[0]
    with h5py.File(os.path.join(output_path, 'predictions_' + video_name + '.h5'), 'w') as h5_file:

        predictions = h5_file.create_dataset('predictions', shape=(0, ), maxshape=(None, ), dtype='|S200')

        cap = skvideo.io.vreader(video_file) # inputdict={'-ss': '02:05:00'}
        
        os.makedirs(os.path.join(output_path, 'predictions_' + video_name), exist_ok=True)

        for idx, frame in enumerate(cap):
            
            images.append(frame)
            tiles = [frame] # use next line if images are too big to load for inference
#             tiles = generate_tiles(frame, size=1024, as_list=True)
            queue.extend(tiles)
            
            while len(queue) >= batch_size:
                results.extend(model.detect(queue[:batch_size]))
                queue = queue[batch_size:]

            while len(results) >= len(tiles):
                    
                r = results[0] # use next line if images are too big to load for inference
#                 r = stitch_predictions(results[:len(tiles)], images[0].shape, size=1024)
                
                # comment following block to prevent image output
                masked_img = draw_instances(images[0], r['rois'][r['scores'] > threshold], r['masks'][:, :, r['scores'] > threshold], r['class_ids'][r['scores'] > threshold], 
                            dataset_val.class_names, r['scores'][r['scores'] > threshold])
                cv2.imwrite(os.path.join(output_path, 'predictions_' + video_name, 'pred_{:>05}.jpg').format(frame_idx), cv2.cvtColor(masked_img, cv2.COLOR_RGB2BGR))
                
                group = h5_file.create_group(str(frame_idx))
                group.create_dataset('masks', data=r['masks'], compression="gzip", compression_opts=9)
                group.create_dataset('scores', data=r['scores'])
                group.create_dataset('classes', data=np.array([classes[c] for c in r['class_ids']], dtype=np.bytes_))
                
                predictions.resize((predictions.shape[0] + 1, ))
                predictions[-1] = np.bytes_(str(frame_idx))
                
                frame_idx += 1
                images = images[1:]
                results = results[len(tiles):]
                
                print('{}'.format(frame_idx), sep=' ', end='\r', flush=True)
```
