## Supplementary material for "High-resolution animal tracking with integration of environmental information in aquatic systems": Code_utils: extract_frames.ipynb

```
# %% [code]
import os
import argparse
import cv2
import numpy as np
from glob import glob
from multiprocessing import Pool

# %% [code]
def extract_frames(video_file, sampling_rate, output_directory, start_frame=0):
    base_name = os.path.splitext(os.path.basename(video_file))[0]
    output_directory = os.path.join(output_directory, base_name)
    cap = cv2.VideoCapture(video_file)
    n_frames = int(cap.get(cv2.CAP_PROP_FRAME_COUNT))
    end_frame = n_frames
    ret = True
    frame_idx = 0
    if not os.path.exists(output_directory) and start_frame == 0:
        os.mkdir(output_directory)
    while ret:
        ret, img = cap.read()
        if frame_idx % sampling_rate == start_frame and frame_idx >= start_frame and frame_idx <= end_frame:
            cv2.imwrite(os.path.join(output_directory, '{}_{:05d}.jpg'.format(base_name, frame_idx)), img)
        frame_idx += 1
        if frame_idx > end_frame:
            break
    return True

class FrameSampler:
    
    def __init__(self, video_file, sampling_rate, output_directory, n_jobs=-1):
        self.n_jobs = os.cpu_count() if n_jobs <= 0 or not isinstance(j_jobs, int) else n_jobs
        self.video_file = video_file
        self.sampling_rate = sampling_rate
        self.output_directory = output_directory
        
    def run(self):
        if self.n_jobs == 1:
            extract_frames(self.video_file, self.sampling_rate, self.output_directory, start_frame=0)
        else:
            process_list = [[self.video_file] * self.n_jobs,
                            [self.sampling_rate * (self.n_jobs + 1)] * self.n_jobs,
                            [self.output_directory] * self.n_jobs,
                            np.arange(0, self.sampling_rate * self.n_jobs, self.sampling_rate)]
            with Pool(processes=self.n_jobs) as pool:
                pool.starmap(extract_frames, list(zip(*process_list)))
                pool.close()
                pool.join()
        return True

# %% [code]
# set paths and sampling rate accordingly
# note sampling_rate refers to every nth frame that should be extracted
# so if you have a 60fps video, sampling_rate=20 will extract 3 frames per second

sampler = FrameSampler(video_file='/path/to/video/video.mp4',
                       sampling_rate=20,
                       output_directory='/path/to/output/images')

# %% [code]
# run the frame sampler

sampler.run()
```
