## Supplementary material for "High-resolution animal tracking with integration of environmental information in aquatic systems": Code_utils: masks_to_tracks.ipynb

```
# %% [code]
from scipy.interpolate import splprep, splev
from skimage.morphology import skeletonize
from scipy.optimize import linear_sum_assignment
from sklearn.metrics.pairwise import euclidean_distances
import numpy as np
import cv2
from sklearn.decomposition import PCA
from scipy.interpolate import interp1d
from scipy.signal import savgol_filter
from scipy.signal import find_peaks_cwt
from sklearn.metrics.pairwise import euclidean_distances
from sklearn.cluster import KMeans
from copy import deepcopy
import matplotlib.pyplot as plt
from scipy import interpolate
from sklearn.neighbors import KDTree
import networkx as nx
from sklearn.neighbors import NearestNeighbors
import os
import h5py
from multiprocessing import Pool, cpu_count, Lock, Value
import pandas as pd
import _pickle

def merge_tiles(tiles, classes, width=2704, height=2028):
    layers = np.sum([tile.shape[2] for tile in tiles])
    classes = np.concatenate(classes)
    shape = tiles[0].shape[:2]
    mask = np.zeros((height, width, layers), dtype=np.bool)
    l = 0
    mask[0:shape[0], 0:shape[1], l:(l + tiles[0].shape[2])] = tiles[0]
    l += tiles[0].shape[2]
    mask[0:shape[0], (width - shape[1]):width, l:(l + tiles[1].shape[2])] = tiles[1]
    l += tiles[1].shape[2]
    mask[(height - shape[0]):height, 0:shape[1], l:(l + tiles[2].shape[2])] = tiles[2]
    l += tiles[2].shape[2]
    mask[(height - shape[0]):height, (width - shape[1]):width, l:(l + tiles[3].shape[2])] = tiles[3]
    return mask, classes

def merge_masks(mask, classes, min_overlap=0.75): # check this out
    for l1, c1 in zip(range(mask.shape[2]), classes):
        for l2, c2 in zip(range(mask.shape[2]), classes):
            if l1 == l2 or c1 != c2:
                continue
            overlap = (mask[:, :, l1] & mask[:, :, l2]).sum()
            if overlap == 0:
                continue
            smaller = np.min([mask[:, :, l1].sum(), mask[:, :, l2].sum()])
            if overlap / smaller > min_overlap:
                merged = mask[:, :, l1] | mask[:, :, l2]
                mask[:, :, l1] = merged
                mask[:, :, l2] = 0
    layers = mask.sum(axis=(0, 1)) > 0
    mask = mask[:, :, layers]
    classes = classes[layers]
    return mask, classes

def process_prediction(prediction, file_name, pose=False, flatten=False):
    with h5py.File(file_name, 'r') as h5_file:
        classes = h5_file[prediction]['classes'][:].astype(np.str)
        masks = h5_file[prediction]['masks'][:]
        scores = h5_file[prediction]['scores'][:]
        count_total = h5_file['predictions'][:].size
    if flatten:
        masks, classes = merge_masks(masks, classes, min_overlap=0.75)
    result = []
    if masks.shape[2] > 0:
        if pose:
            result = generate_poses(masks, n_pts=15)
        else:
            result = generate_positions(masks)
    lock.acquire()
    count.value += 1
    print('{:.2f} %'.format(100 * count.value / count_total), sep=' ', end='\r', flush=True)
    lock.release()
    return result

def approve_assignment(cost_matrix, assignment, max_cost):
    if np.any(cost_matrix[assignment[0], assignment[1]] >= max_cost):
        return False
    return True

def draw_contour(contour, width, height):
    contour = cv2.drawContours(np.zeros((height.astype(np.int) + 1, width.astype(np.int) + 1, 3), dtype=np.uint8), [contour.astype(np.int)], 0, (255, 255, 255), -1, cv2.LINE_AA)
    contour = contour.sum(axis=2) > 0
    return contour

def contour_overlap(contours_current, contours_previous):
    max_cost = np.inf
    cost_matrix = np.zeros((len(contours_current), len(contours_previous))) + max_cost
    for i, current in enumerate(contours_current):
        for j, previous in enumerate(contours_previous):
            intersection = np.sum(current & previous)
            union = np.sum(current | previous)
            cost_matrix[i, j] = union / intersection if intersection > 0 else max_cost
    if np.isfinite(cost_matrix).sum() > 0:
        max_cost = cost_matrix[np.isfinite(cost_matrix)].max() * 10
        cost_matrix[np.isinf(cost_matrix)] = max_cost
    else:
        max_cost = 0
        cost_matrix[:, :] = max_cost
    return cost_matrix, max_cost

def centroid_distance(contours_current, contours_previous):
    max_cost = np.inf
    cost_matrix = np.zeros((len(contours_current), len(contours_previous))) + max_cost
    for i, current in enumerate(contours_current):
        for j, previous in enumerate(contours_previous):
            cost_matrix[i, j] = np.sqrt(np.square(np.argwhere(current).mean(axis=0) - np.argwhere(previous).mean(axis=0)).sum())
    if np.isfinite(cost_matrix).sum() > 0:
        max_cost = cost_matrix[np.isfinite(cost_matrix)].max() * 10
        cost_matrix[np.isinf(cost_matrix)] = max_cost
    else:
        max_cost = 0
        cost_matrix[:, :] = max_cost
    return cost_matrix, max_cost

def assign_identities(positions, frame_idx, contours=None, max_merge_dist=30):
    identities = np.arange(positions.shape[0])
    unique_frame_idx = np.arange(frame_idx.min(), frame_idx.max() + 1)
    detection_count = np.bincount(frame_idx)[frame_idx.min():]
    if contours is None:
        max_cost = max_merge_dist
    for idx, (frame, count) in enumerate(zip(unique_frame_idx, detection_count)):
        print('{:.2f} %'.format(100 * idx / unique_frame_idx.size), sep=' ', end='\r', flush=True)
        if frame == frame_idx.min():
            continue
        else:
            if contours is None:
                positions_current = positions[frame_idx == frame]
                positions_previous = positions[frame_idx == frame - 1]
                if len(positions_current) == 0 or len(positions_previous) == 0:
                    continue
                cost_matrix = euclidean_distances(positions_current, positions_previous)
            else:
                contours_current = contours[frame_idx == frame]
                contours_previous = contours[frame_idx == frame - 1]
                if len(contours_current) == 0 or len(contours_previous) == 0:
                    continue
                contours_concatenated = np.concatenate([np.concatenate([c for c in contours_current], axis=0),
                                                        np.concatenate([c for c in contours_previous], axis=0)])
                xmin, ymin = contours_concatenated.min(axis=0)
                xmax, ymax = contours_concatenated.max(axis=0)
                contours_current = [draw_contour(c - np.array([xmin, ymin]), width=xmax - xmin, height=ymax - ymin) \
                                    for c in contours_current]
                contours_previous = [draw_contour(c - np.array([xmin, ymin]), width=xmax - xmin, height=ymax - ymin) \
                                    for c in contours_previous]
                cost_matrix, max_cost = contour_overlap(contours_current, contours_previous)
            assignment = linear_sum_assignment(cost_matrix)
            if not approve_assignment(cost_matrix, assignment, max_cost=max_cost):
                continue
            if count == 1 and detection_count[idx - 1] == 1:
                identities[frame_idx == frame] = int(identities[frame_idx == frame - 1])
            elif count <= detection_count[idx - 1]: # and count > 0
                identities[frame_idx == frame] = identities[frame_idx == frame - 1][assignment[1]]
            elif count > detection_count[idx - 1] and count > 1: # and detection_count[idx - 1] > 0
                current_identities = identities[frame_idx == frame]
                current_identities[assignment[0]] = identities[frame_idx == frame - 1][assignment[1]]
                identities[frame_idx == frame] = current_identities
    for idx, identity in enumerate(np.unique(identities)):
        identities[identities == identity] = idx
    print('{:.2f} %'.format(100))
    return identities

def generate_poses(mask, n_pts):    
    mask = mask.reshape(mask.shape[0], mask.shape[1], -1).astype(np.uint8) * 255
    poses = []
    for layer in range(mask.shape[2]): # mask_copy.shape[2]):
        img = mask[:, :, layer].reshape(mask.shape[0], mask.shape[1])
        poses.append(get_pose(img, n_pts))
    return poses

def generate_positions(mask):
    mask = mask.reshape(mask.shape[0], mask.shape[1], -1).astype(np.uint8) * 255
    positions = []
    for layer in range(mask.shape[2]): # mask_copy.shape[2]):
        img = mask[:, :, layer].reshape(mask.shape[0], mask.shape[1])
        if img.sum() > 0:
            positions.append(get_position(img))
    return positions

def get_position(img):
    return np.mean(np.argwhere(img), axis=0)[::-1]

def get_pose(img, n_pts):
    contours = cv2.findContours(img.copy(), cv2.RETR_TREE, cv2.CHAIN_APPROX_SIMPLE)[-2] # img?
    contours = [cnt.reshape(-1, 2) for cnt in contours if cnt.size > 10]
    contours = [np.append(cnt, cnt[0].reshape(1, 2), axis=0) for cnt in contours]
    if len(contours) == 0:
        return [[]] * 3
    contours = interpol_objects(contours)
    contour = np.concatenate(contours)
    if contour.shape[0] < 5: # and maybe ratio
        return [[]] * 3
    img_copy = img.copy()
    img, xmin, ymin, scale = crop_scale_mask(img, max_len=100)
    skeleton = skeletonize(img > 0)
    pts = np.argwhere(skeleton).reshape(-1, 2)
    mean_distances = euclidean_distances(pts, pts).mean(axis=1)
    end_idx = np.argsort(mean_distances)[-(pts.shape[0] // 4):].reshape(-1)
    end_pts = pts[end_idx]
    if end_pts.shape[0] < 2:
        return [[]] * 3
    kmeans = KMeans(n_clusters=2)
    kmeans.fit(end_pts)
    end_pts = np.round(kmeans.cluster_centers_).astype(np.int)
    if False in [(pt == pts).all(axis=1).any() for pt in end_pts]:
        pts = np.unique(np.concatenate([pts, end_pts]), axis=0)
    end_idx = [np.argwhere((pts == pt).all(axis=1))[0][0] for pt in end_pts]
    nn = 2
    path = []
    while True and nn < pts.shape[0]:
        try:
            clf = NearestNeighbors(nn).fit(pts)
            G = clf.kneighbors_graph()
            T = nx.from_scipy_sparse_matrix(G)

            path = nx.algorithms.shortest_path(T, source=end_idx[0], target=end_idx[1])
            break
        except:
            nn = nn * 2
    if len(path) == 0:
        return [[]] * 3
    spine = pts[path]
    spine = (spine / scale) + np.array([ymin, xmin])
    window_length = spine.shape[0] // 16
    if window_length < 3:
        window_length = 3
    elif window_length % 2 == 0:
        window_length += 1
    if window_length <= spine.shape[0]:
        spine[:, 0] = savgol_filter(spine[:, 0], window_length=window_length, polyorder=1)
        spine[:, 1] = savgol_filter(spine[:, 1], window_length=window_length, polyorder=1)
    dists = np.cumsum(np.sqrt(np.square(np.diff(spine, axis=0)).sum(axis=1)))
    dists = np.concatenate([[0], dists / dists[-1]])
    fx, fy = interp1d(dists, spine[:, 1]), interp1d(dists, spine[:, 0])
    regular = np.linspace(0, 1, n_pts)
    x_regular, y_regular = fx(regular), fy(regular)
    spine = np.transpose([y_regular, x_regular])
    rad = []
    for pt in spine:
        rad.append(np.sort(np.sqrt(np.square(contour - pt[::-1]).sum(axis=1)))[0])
    rad = np.array(rad)
    head_radii = rad[2]
    tail_radii = rad[-2]
    if head_radii < tail_radii:
        spine = spine[::-1]
        rad = rad[::-1]
    return [spine, rad, contour]

def edge_length(edge):
    return np.sum(np.sqrt(np.square(np.diff(edge[:, 0])) + np.square(np.diff(edge[:, 1]))))

def interpol_objects(objects, precision = 5):
    interpolated = []
    for obj in objects:
        length = edge_length(obj)
        tckp, u = splprep([obj[:, 0], obj[:, 1]], s = 0, k = 1)
        obj_x, obj_y = splev(np.linspace(0, 1, 100), tckp)
        obj = np.transpose((obj_x, obj_y))
        interpolated.append(obj)
    return interpolated

def wrap(angles):
    angles = (angles + np.pi) % (2 * np.pi) - np.pi
    return angles

def crop_scale_mask(mask, max_len=100):
    ymin, xmin = np.min(np.argwhere(mask), axis=0)
    ymax, xmax = np.max(np.argwhere(mask), axis=0)
    cropped = mask[ymin:(ymax + 1), xmin:(xmax + 1)]
    scale = max_len / np.max(cropped.shape)
    scaled = cv2.resize(cropped, (0, 0), fx=scale, fy=scale, interpolation=cv2.INTER_AREA)
    return scaled, xmin, ymin, scale

def i_lookup(i):
    return np.isin(frames, tracks[str(i)]['FRAME_IDX'])

def init_lookup(f, t):
    global frames, tracks
    frames = f
    tracks = t

def identity_lookup(tracks):
    frames = np.array([])
    if tracks['FRAME_IDX'].size > 0:
        frames = np.arange(tracks['FRAME_IDX'].min(), tracks['FRAME_IDX'].max() + 1)
    with Pool(processes=cpu_count(), initializer=init_lookup, initargs=(frames, tracks, )) as pool:
        lookup = pool.map(i_lookup, tracks['IDENTITIES'])
        pool.close()
        pool.join()
    lookup = np.transpose(lookup)
    return lookup

def append_trajectory(trajectory_pre, trajectory_post):
    if np.isin(trajectory_pre['FRAME_IDX'], trajectory_post['FRAME_IDX']).any():
        return trajectory_pre
    appended = {}
    sorted_idx = np.argsort(np.append(trajectory_pre['FRAME_IDX'], trajectory_post['FRAME_IDX']))
    for key in trajectory_pre:
        appended[key] = np.append(trajectory_pre[key], trajectory_post[key], axis=0)[sorted_idx]
    return appended

def get_positions(identities, tracks, idx):
    positions = []
    for i in identities:
        positions.append([tracks[str(i)]['X'][idx], tracks[str(i)]['Y'][idx]])
    return np.array(positions)

def merge_identities(tracks, lookup, max_gap=10, max_distance=20, validate=False, min_size=2):
    frame_idx = np.arange(lookup.shape[0])
    for idx, i in enumerate(range(lookup.shape[1])):
        while True:
            i_lu = np.argwhere(lookup[:, i])
            if i_lu.size == 0:
                break
            i_end = i_lu.max()
            i_position = get_positions([tracks['IDENTITIES'][i]], tracks, -1)
            next_filter = (frame_idx > i_end) & (frame_idx <= i_end + max_gap)
            next_identities = np.argwhere(lookup[next_filter, :].sum(axis=0) > 0).ravel()
            next_start = np.argmax(lookup[:, next_identities], axis=0)
            next_identities = next_identities[next_start > i_end]
            next_start = next_start[next_start > i_end]
            if next_identities.size == 0:
                break
            if validate:
                previous_filter = (frame_idx < np.min(next_start)) & (frame_idx >= i_end - max_gap)
                previous_identities = np.argwhere(lookup[previous_filter, :].sum(axis=0) > 0).ravel()
                previous_identities = previous_identities[previous_identities != i]
                previous_end = lookup.shape[0] - np.argmax(np.flip(lookup[:, previous_identities], axis=0), axis=0) - 1
                previous_identities = previous_identities[previous_end < np.min(next_start)]
                previous_end = previous_end[previous_end < np.min(next_start)]
                if previous_identities.size == 0:
                    break
                previous_positions = get_positions(tracks['IDENTITIES'][previous_identities], tracks, -1)
                validation_cost = euclidean_distances(i_position, previous_positions).ravel()
                if np.any(validation_cost < max_distance):
                    break
            next_positions = get_positions(tracks['IDENTITIES'][next_identities], tracks, 0)
            cost = euclidean_distances(i_position, next_positions).ravel()
            next_identities = next_identities[cost < max_distance]
            next_start = next_start[cost < max_distance]
            if next_identities.size == 0:
                break
            next_identity = next_identities[np.argmin(next_start)]
            if np.isin(tracks[str(tracks['IDENTITIES'][i])]['FRAME_IDX'],
                          tracks[str(tracks['IDENTITIES'][next_identity])]['FRAME_IDX']).any():
                break
            lookup[:, next_identity] = 0
            tracks[str(tracks['IDENTITIES'][i])] = append_trajectory(tracks[str(tracks['IDENTITIES'][i])],
                                                 tracks[str(tracks['IDENTITIES'][next_identity])])
            lookup[:, i] = np.isin(frame_idx, tracks[str(tracks['IDENTITIES'][i])]['FRAME_IDX'])
        print('{:.2f} %'.format(100 * idx / lookup.shape[1]), sep=' ', end='\r', flush=True)
    identity_filter = lookup.sum(axis=0) > min_size
    remove_identities = tracks['IDENTITIES'][np.invert(identity_filter)]
    tracks['IDENTITIES'] = tracks['IDENTITIES'][identity_filter]
    lookup = lookup[:, identity_filter]
    for i in remove_identities:
        tracks.pop(str(i))
    print('{:.2f} %'.format(100))
    return tracks, lookup

def init(l, c):
    global lock, count
    lock = l
    count = c

def masks_to_tracks(file_name, use_contours=False, flatten_masks=False, max_merge_dist=30):
    with h5py.File(file_name, 'r') as h5_file:
        predictions = h5_file['predictions'][:].astype(np.str)
        
    sort_idx = np.argsort([int(prediction.split('_')[0]) for prediction in predictions])
    predictions = predictions[sort_idx][:10000]
    
    l = Lock()
    c = Value('d', 0)

    print('Generating {}:'.format('poses' if use_contours else 'positions'))
    with Pool(processes=cpu_count(), initializer=init, initargs=(l, c)) as pool:

        output_list = pool.starmap(process_prediction, list(zip(predictions,
                                                                [file_name] * len(predictions),
                                                                [use_contours] * len(predictions),
                                                                [flatten_masks] * len(predictions))))
        pool.close()
        pool.join()
        print('{:.2f} %'.format(100))
        
    frame_idx = np.array([int(prediction.split('_')[0]) for prediction in predictions])
    
    if use_contours:
        positions = np.array([pose[0].mean(axis=0) for poses in output_list for pose in poses if len(pose[0]) > 0])
        frame_idx = np.array([frame for frame, poses in zip(frame_idx, output_list) for pose in poses if len(pose[0]) > 0])
        spines = np.array([pose[0] for poses in output_list for pose in poses if len(pose[0]) > 0])
        radii = np.array([pose[1] for poses in output_list for pose in poses if len(pose[0]) > 0])
        contours = np.array([pose[2] for poses in output_list for pose in poses if len(pose[0]) > 0])
    else:
        positions = np.array([position for positions in output_list for position in positions if len(positions) > 0])
        frame_idx = np.array([frame for frame, positions in zip(frame_idx, output_list) for position in positions if len(positions) > 0])
        contours = None
        spines = None

    print('Assigning identities:')
    identities = assign_identities(positions, frame_idx, contours=contours, max_merge_dist=max_merge_dist)
    
    n_inds = np.unique(identities).size
    print('Number of individuals: {}'.format(n_inds))
    
    print('Generating tracks:')
    
    tracks = {}
    tracks['FRAME_IDX'] = []
    for idx, i in enumerate(np.unique(identities)):
        tracks[str(i)] = {}
        tracks[str(i)]['FRAME_IDX'] = frame_idx[identities == i]
        tracks['FRAME_IDX'].append(tracks[str(i)]['FRAME_IDX'])
        if spines is not None:
            tracks[str(i)]['X'] = spines[:, 1, 1][identities == i]
            tracks[str(i)]['Y'] = spines[:, 1, 0][identities == i]
            tracks[str(i)]['SPINE'] = spines[identities == i]
            tracks[str(i)]['RADII'] = radii[identities == i]
            tracks[str(i)]['CONTOUR'] = contours[identities == i]
            heading = np.arctan2(spines[:, 0, 0][identities == i] - spines[:, 1, 0][identities == i],
                                 spines[:, 0, 1][identities == i] - spines[:, 1, 1][identities == i])
            tracks[str(i)]['HEADING'] = heading
        else:
            tracks[str(i)]['X'] = positions[:, 0][identities == i]
            tracks[str(i)]['Y'] = positions[:, 1][identities == i]
        print('{:.2f} %'.format(100 * idx / n_inds), sep=' ', end='\r', flush=True)
    tracks['IDENTITIES'] = np.unique(identities).astype(np.int)
    tracks['FRAME_IDX'] = np.unique(np.concatenate(tracks['FRAME_IDX'])).astype(np.int)
    print('{:.2f} %'.format(100))
    
    def print_iteration(iteration, lookup):
        print('Iteration {}: {} individuals'.format(iteration, lookup.shape[1]))
        iteration += 1
    
    print('Autostitching tracks')
    lookup = identity_lookup(tracks)
    n_inds = lookup.shape[1] + 1
    iteration = 0
    while lookup.shape[1] < n_inds:
        n_inds = lookup.shape[1]
        tracks, lookup = merge_identities(tracks, lookup, validate=True, min_size=1)
        print_iteration(iteration, lookup)
        tracks, lookup = merge_identities(tracks, lookup, max_gap=30, validate=True, min_size=1)
        print_iteration(iteration, lookup)
        tracks, lookup = merge_identities(tracks, lookup, max_gap=60, max_distance=100, validate=True, min_size=2)
        print_iteration(iteration, lookup)
        tracks, lookup = merge_identities(tracks, lookup, max_gap=60, max_distance=100, validate=True, min_size=5)
        print_iteration(iteration, lookup)
    return tracks

def plot_tracks(tracks, figsize=(20, 20), resolution=None, legend=False):
    fig = plt.figure(figsize=figsize)
    ax = fig.add_axes([0, 0, 1, 1])
    for i in tracks['IDENTITIES']:
        if 'SPINE' in tracks[str(i)]:
            if tracks[str(i)]['X'].size > 1:

                ax.plot(tracks[str(i)]['SPINE'][:, :, 1].mean(axis=1), tracks[str(i)]['SPINE'][:, :, 0].mean(axis=1), label=i)
            else:
                ax.plot(tracks[str(i)]['SPINE'][:, :, 1].mean(axis=1), tracks[str(i)]['SPINE'][:, :, 0].mean(axis=1) ,'.', label=i)
        else:
            if tracks[str(i)]['X'].size > 1:
                ax.plot(tracks[str(i)]['X'], tracks[str(i)]['Y'], label=i)
            else:
                ax.plot(tracks[str(i)]['X'], tracks[str(i)]['Y'], '.', label=i)
    ax.set_aspect('equal')
    if resolution is not None:
        ax.set_ylim((resolution[1], 0))
        ax.set_xlim((0, resolution[0]))
    if legend:
        plt.legend()
    ax.set_xlim()

def save(dump, file_name):
    with open(file_name, 'wb') as fid:
        _pickle.dump(dump, fid)
    return True

def load(file_name):
    with open(file_name, 'rb') as fid:
        dump = _pickle.load(fid)
    return dump

def tracks_to_pool(tracks):
    x = np.array([])
    y = np.array([])
    frame_idx = np.array([], dtype = np.int)
    identity = np.array([], dtype = np.int)
    for i in tracks['IDENTITIES']:
        x = np.append(x, tracks[str(i)]['X'])
        y = np.append(y, tracks[str(i)]['Y'])
        frame_idx = np.append(frame_idx, tracks[str(i)]['FRAME_IDX'])
        identity = np.append(identity, np.repeat(i, tracks[str(i)]['FRAME_IDX'].size))
    tracks = {}
    tracks['X'] = x
    tracks['Y'] = y
    tracks['FRAME_IDX'] = frame_idx
    tracks['IDENTITY'] = identity
    return tracks

def write_tracks_pooled(tracks, file_name):
    if 'IDENTITIES' in tracks:
        tracks = tracks_to_pool(tracks)
    track_data = pd.DataFrame.from_dict(tracks)
    track_data.to_csv(file_name, index=False)
    return True

# %% [code]
# set path to prediction file from mask rcnn inference
# decide whether to estimate fish spine poses or to just use the centroid of the mask
# set flatten_masks=True if you want to merge overlapping masks per frame (use this if you used the tiling for mask rcnn inference)
# set maximum merge distance (pixel) between subsequent frames

tracks = masks_to_tracks('path/to/prediction_file.h5',
                         use_contours=True,
                         flatten_masks=True,
                         max_merge_dist=300)
save(tracks, file_name=i.replace('predictions', 'tracks').replace('h5', 'pkl'))
```
