## Supplementary material for "High-resolution animal tracking with integration of environmental information in aquatic systems": Code_utils: readme_utils.txt

install a conda or miniconda environment with jupyter notebook to open the interactive python notebook (.ipynb)
use synchronize.ipynb to synchronize multi-view videos of one trial
use extract_frames.ipynb to extract frames for COLMAP input
use mask_to_tracks.ipynb to estimate positions/poses based on mask rcnn predictions, to automatically assign identities and to save tracks for correction in TrackUtil
