## Supplementary material for "High-resolution animal tracking with integration of environmental information in aquatic systems": Code_utils: synchronize.ipynb

```
# %% [code]
from scipy.io import wavfile
from scipy.signal import fftconvolve

import json
import os

import numpy as np
import subprocess

def parse_metadata(video_file):
    temp_file = os.path.splitext(video_file)[0] + '.json'
    subprocess.run('ffprobe -v quiet -print_format json -show_format -show_streams {} > {}'.format(video_file,
                                                                                                   temp_file),
                   shell=True)
    metadata = json.load(open(temp_file, 'r'))
    os.remove(temp_file)
    fps = np.array(metadata['streams'][0]['r_frame_rate'].split('/')).astype(np.int)
    channels = int(metadata['streams'][1]['channels'])
    sample_rate = int(metadata['streams'][1]['sample_rate'])
    fps = fps[0] / fps[1]
    return {'fps': fps,
            'sample_rate': sample_rate,
            'channels': channels}

def extract_audio(video_file, start=0, duration=30):
    metadata = parse_metadata(video_file)
    
    temp_file = os.path.splitext(video_file)[0] + '.wav'
    subprocess.run('ffmpeg -y -v quiet -i {} -vn -c:a pcm_s16le -ss {} -t {} {}'.format(video_file,
                                                                                     start,
                                                                                     duration,
                                                                                     temp_file),
                   shell=True)
    sample_rate, signal = wavfile.read(temp_file)
    os.remove(temp_file)
    assert (sample_rate == metadata['sample_rate']) & (signal.shape[1] == metadata['channels']), \
        'audio stream did not match video metadata'
    if metadata['channels'] > 1:
        signal = signal.mean(axis=1)
    return signal

def compute_offset(audio_1, audio_2, sample_rate):
    corr = fftconvolve(audio_1, audio_2[::-1], mode='full')
    offset = np.argmax(corr)
    offset_seconds = ((2 * audio_2.size - 1) // 2 - offset) / sample_rate
    return offset_seconds

def check_input(args, video_files):
    checked_args = []
    for arg in args:
        if isinstance(arg, (float, int)):
            arg = [arg] * len(video_files)
        elif arg is None:
            arg = [30] * len(video_files)
        assert isinstance(arg, list) and (len(arg) == len(video_files)), \
            'start and duration must be either None, or of type int, float or list with length of video files'
        checked_args.append(arg)
    return checked_args

def cut_video(video_file, offset):
    cut_file = os.path.splitext(video_file)[0] + '_cut' + os.path.splitext(video_file)[1]
    subprocess.run('ffmpeg -v quiet -i {} -ss {} -c:v libx264 -an -crf 18 -preset ultrafast {}'.format(video_file,
                                                                                                       offset,
                                                                                                       cut_file),
                   shell=True)
    return True

def copy_video(video_file):
    cut_file = os.path.splitext(video_file)[0] + '_cut' + os.path.splitext(video_file)[1]
    subprocess.run('cp {} {}'.format(video_file, cut_file), shell=True)
    return True

def synchronize_videos(video_files, start=None, duration=None, dry_run=False):
    metadata = [parse_metadata(video_file) for video_file in video_files]
    assert len(set([metadata[idx]['fps'] for idx in range(len(metadata))])) == 1, \
        'videos must have the same frame rate'
    assert len(set([metadata[idx]['fps'] for idx in range(len(metadata))])) == 1, \
        'audio streams must have the same sample rate'
    start, duration = check_input([start, duration], video_files)
    signals = [extract_audio(video_file, start=s, duration=d) \
               for video_file, s, d in zip(video_files, start, duration)]
    offsets = [0]
    ref = signals[0]
    signals = signals[1:]
    for sig in signals:
        offsets.append(compute_offset(ref, sig, metadata[0]['sample_rate']))
    offsets = np.array(offsets) - np.min(offsets)
    for idx, video_file in enumerate(video_files):
        print(video_file, offsets[idx])
        if dry_run:
            continue
        if offsets[idx] * metadata[idx]['fps'] < 1:
            copy_video(video_file)
        else:
            cut_video(video_file, offsets[idx])
    return True

# %% [code]
# add the video files to following list, set duration of audio signal used for synchronization
# dry_run=True can be used to just print the video offsets without cutting
# ffmpeg needs to be installed (this is only tested on linux systems)

video_files = ['/path/to/video1/video1.mp4',
               '/path/to/video2/video2.mp4',
               '/path/to/video3/video3.mp4']
synchronize_videos(video_files, duration=180, dry_run=False)
```
