## Supplementary material for "High-resolution animal tracking with integration of environmental information in aquatic systems": Practical implementation guide

A Practical Guide

### 1 Data collection

Our approach requires little preparation in the field, however, some things need to be considered. In order to successfully triangulate 3D animal trajectories, we need at least two detections per time point (i.e. video frame) of each animal throughout the videos. Hence, we need at least two cameras in our multi-view camera array. For better coverage, higher overlap and easier filming, four cameras are recommended (arranged in a square). The size of the observed animals and the distance from which they are filmed determine the camera-to-camera distances in the array (and correspondingly the video overlap). We found a distance of 0.6 m between the cameras appropriate when filming from one to three meters above the animals. See Figure 1 for example setups.

Since the Structure-from-Motion (SfM) approach relies on moving cameras, the camera array should not remain stationary throughout a full recording. This is not problematic when swimming/diving above mobile animals (as for our *'single'* and *'school'* datasets), but when filming site-specific animals for a longer duration ($> 5$ min) it may become tedious for the observer and increase later computational load for the SfM algorithm. Here, we suggest using a semi-stationary camera array mounted on a tripod (as for the *'mixed'* dataset), that can be moved when the animals are moving, and left stationary when the animals stay in one spot for some time.

Finally, the observer should make a noise after all cameras were turned on (e.g. knocking on the camera array), so that the videos can be easily synchronized.

### 2 Video synchronization

Generally, video cutting and synchronization should be the first step after filming animals in the field. This will reduce later data management and organization. We provide a Python script, that extracts audio signals from the videos, convolves the Fourier-transformed signals to calculate offsets and cuts the videos so that they are synchronized, using ffmpeg and common Python packages (e.g. scipy).

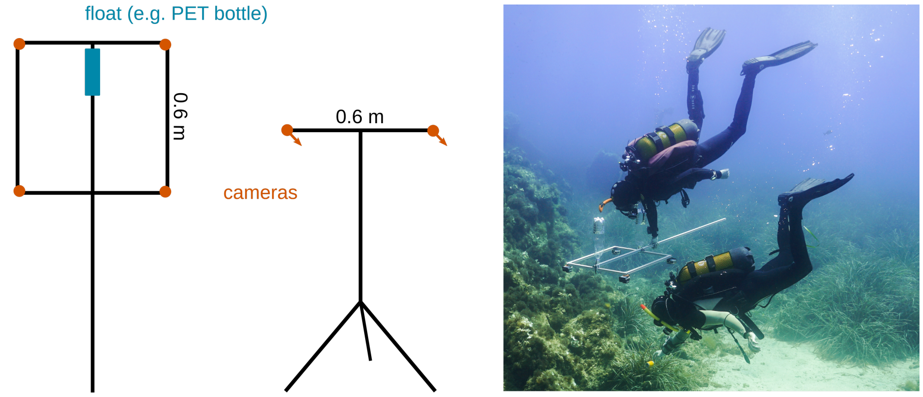


Figure 1: Example setups for the multi-view camera arrays. When filming mobile animals, we recommend using four cameras (left). If the observed animals are site-specific, a semi-stationary setup may be preferable, that is only moved when the animals change their locations (middle). The setup we used for recording the videos of the *'accuracy'* dataset (right).

### 3 Mask R-CNN training and inference

The next step is training a Mask R-CNN model for object detection and instance segmentation to generate pixel locations of the observed animals from the synchronized videos. Mask R-CNN is a complex convolutional neural network that would take a long time to train from scratch, but we can use transfer-learning (starting from a pre-trained snapshot using the COCO dataset) to speed up the process.

First, a custom dataset containing image annotations has to be created. We developed a purpose-oriented GUI based on Python and Qt5 within the lab (*'TrackUtil'*), that allows to mask objects of interest during video playback. It supports interactive annotation of multiple classes (for example when multiple species were observed), as well as simultaneous playback of multiple videos, and outputs the dataset in a format that can be directly imported for Mask R-CNN training. *'TrackUtil'* is available in the additional files section, but will be released on GitHub once it is fully documented.

The resulting dataset containing the images and respective pixel masks is then used as input to the training process, which is well-documented in the original GitHub repository. We customized the code for input, inference (i.e. using the model to predict animal locations on the videos), and output (Additional file 5). Training and inference both require GPU-accelerated systems, and can be performed either on local hardware, on computing clusters (for example available through respective research institute or university) or free computing services such as Google Colaboratory.

### 4 Structure-from-Motion

We use SfM to reconstruct the trajectories of the cameras from the video recordings, and to generate 3D point clouds of the filmed environment. Further, the camera calibration parameters are obtained through SfM. Our approach implements input and output for one implementation of SfM that we found well-documented and performant, COLMAP. However, free alternative implementations exist (e.g. Theia and Meshroom) and can be used by changing input and output data formatting.

First, images have to be extracted. For this purpose, we provide a Python script (Additional file 6). We recommend a sampling rate of 3 Hz, so that fast camera motion will be well represented in the resulting reconstructions without too much computational load. For all subsequent SfM steps, using the default COLMAP options will result in good reconstruction results in most cases. If a semi-stationary setup is used, images that were extracted from time points without camera motion should be discarded before starting the reconstruction process. In case of very long observations, or many cameras, we recommend splitting the reconstructions into parts with approximately 3000 images each. COLMAP provides functionality to merge reconstructions that are based on the same dataset.

It is sufficient to only run the sparse reconstruction for the subsequent triangulation of 3D trajectories. At this step, each image is referenced in the reconstruction as a 3D location of the camera view point, and a sparse point cloud of the visual scene is reconstructed. Additionally, COLMAP can be used to obtain a denser representation of this scene, useful for visualization and environmental analyses. However, this is computationally demanding and requires a GPU, and we do not include environmental analyses (e.g. complexity measures or habitat segmentation).

### 5 Triangulating trajectories

From the raw Mask R-CNN predictions (binary pixel masks), animal positions have to be determined in pixel coordinate space. These positions can be either estimated as the pixel mask centroid, or an approximation of fish spine pose. For both cases, and for the subsequent generation of animal trajectories, we provide Python code (Additional file 6). The resulting pixel trajectories are very likely fractured and may contain misidentifications, making manual track corrections necessary in almost all cases. Further, trajectory ids have to be matched between the multi-view trajectories. All of these tasks can be accomplished using the interactive functionality of *'TrackUtil'* (Additional file 4).

Finally, 3D trajectories can be obtained through triangulation of the multi-view Mask R-CNN detections using the camera parameters estimated with SfM. For this purpose, we implemented *'multiviewtracks'*, a Python module available on GitHub. Examples for general usage, visualization and export of 3D models can be found in the documentary.
